# Tuberculosis resistance acquisition in space and time: an analysis of globally diverse *M. tuberculosis* whole genome sequences

**DOI:** 10.1101/837096

**Authors:** Yasha Ektefaie, Avika Dixit, Luca Freschi, Maha Farhat

**Affiliations:** University of California Berkeley, Berkeley CA, 120 Sproul Hall, Berkeley CA 94720, USA; Boston Children’s Hospital, Boston MA; Harvard Medical School, Boston MA, 300 Longwood Ave, Mailstop 3103, Boston MA 02115, USA; Boston Children’s Hospital, Boston MA, Harvard Medical School, Boston BA, 10 Shattuck St, Boston MA 02115, USA; Massachusetts General Hospital, Boston, MA, 10 Shattuck St, Boston MA 02115, USA

**Keywords:** tuberculosis, drug resistance, whole genome sequencing

## Abstract

**Background:** *Mycobacterium tuberculosis* (MTB) whole genome sequencing data can provide insights into temporal and geographic trends in resistance acquisition and inform public health interventions.

**Methods:** We curated a set of clinical MTB isolates with high quality sequencing and culture-based drug susceptibility data spanning four lineages and more than 20 countries. We constructed geographic and lineage specific MTB phylogenies and used Bayesian molecular dating to infer the most-recent-common-susceptible-ancestor age for 4,869 instances of resistance to 10 drugs.

**Findings:** Of 8,550 isolates curated, 6,099 from 15 countries met criteria for molecular dating. The number of independent resistance acquisition events was lower than the number of resistant isolates across all countries, suggesting ongoing transmission of drug resistance. Ancestral age distributions supported the presence of old resistance, ≥20 years prior, in the majority of countries. A consistent order of resistance acquisition was observed globally starting with resistance to isoniazid, but resistance ancestral age varied by country. We found a direct correlation between country wealth and resistance age (*R^2^*= 0.47, P-value= 0.014). Amplification of fluoroquinolone and second-line injectable resistance among multidrug-resistant isolates is estimated to have occurred very recently (median ancestral age 4.7 years IQR 1.9-9.8 prior to sample collection). We found the sensitivity of commercial molecular diagnostics for second-line resistance to vary significantly by country (P-value <0.0003)

**Interpretation:** Our results highlight that both resistance transmission and amplification are contributing to disease burden globally but are variable by country. The observation that wealthier nations are more likely to have old resistance suggests that programmatic improvements can reduce resistance amplification, but that fit resistant strains can circulate for decades subsequently.

**Funding:** This work was supported by the NIH BD2K grant K01 ES026835, a Harvard Institute of Global Health Burke Fellowship (MF), Boston Children’s Hospital OFD/BTREC/CTREC Faculty Career Development Fellowship and Bushrod H. Campbell and Adah F. Hall Charity Fund/Charles A. King Trust Postdoctoral Fellowship (AD).

**Research in context:** *Evidence before this study:* Acquisition and spread of drug-resistance by *Mycobacterium tuberculosis* (MTB) varies across countries. Local factors driving evolution of drug resistance in MTB are not well studied.

*Added value of this study:* We applied molecular dating to 6,099 global MTB patient isolates and found the order of resistance acquisition to be consistent across the countries examined, *i.e.* acquisition of isoniazid resistance first followed by rifampicin and streptomycin followed by resistance to other drugs. In all countries with data available there was evidence for transmission of resistant strains from patient-to-patient and in the majority for extended periods of time (>20 years). Countries with lower gross wealth indices were found to have more recent resistance acquisition to the drug rifampicin. Based on the resistance patterns identified in our study we estimate that commercial diagnostic tests vary considerably in sensitivity for second-line resistance diagnosis by country.

*Implications of all available evidence:* The longevity of resistant MTB in many parts of the world emphasizes its fitness for transmission and its continued threat to public health. The association between country wealth and recent resistance acquisition emphasizes the need for continued investment in TB care delivery and surveillance programs. Geographically relevant diagnostics that take into account a country’s unique distribution of resistance are necessary.

## Introduction

Tuberculosis (TB) defines a global epidemic that takes more lives than any other infection due to a single pathogen^1^. The emergence of multidrug-resistant (MDR) and extensively drug-resistant (XDR) resistant TB presents a major hurdle to efforts in accelerating TB decline. Halting the transmission of drug-resistant (DR) TB has been a major focus of studies addressing this hurdle^2^. But the epidemic is ultimately defined by local factors that remain understudied in many parts of the world^3^. The study of geographic and temporal heterogeneity of the DR-TB epidemic can provide insights into these local factors as key drivers of MDR-TB prevalence and persistence in the community, including programmatic and bacterial factors. This understanding is key to future disease control and prevention of antibiotic resistance development.

Over the past decade, increased uptake of molecular and whole genome sequencing (WGS) technologies, and their application to *Mycobacterium tuberculosis* (MTB) clinical isolates has offered novel insights into pathogen biology and diversity in the context of human infection^4–7^. The application of WGS has allowed us to better understand the genetic determinants of drug resistance (DR) within MTB^8^. The detection of these genetic determinants using molecular technologies that include WGS is now increasingly adopted for TB resistance diagnosis in many parts of the world^9^ and is beginning to replace the more biohazardous and time consuming culture based drug susceptibility tests (DST). The study of isolates sampled from epidemiological outbreaks or from the same host over time has allowed the estimation of MTB’s molecular clock rate, or temporal rate of accumulation of fixed genome-level variation^10, 11^. The application of this rate to new WGS data from isolates collected for surveillance has helped improve transmission inference and molecular dating of specific evolutionary events such as resistance acquisition or lineage divergence^10, 12–13^.

We sought to use a large clinical collection of MTB WGS and resistance phenotype data to study how, when, and where resistance was acquired on a global scale. Using a Bayesian implementation of coalescent theory, we estimate and compare dates of resistance acquisition for MDR/XDR isolates across 15 different countries. We use the recency of resistance acquisition as a measure of fitness of the circulating strains in their respective environments and study the effect of country wealth, as a proxy for TB control programme funding, on the recency of resistance acquisition at a macro level. We also assess the distribution of unexplained MTB phenotypic resistance across 20 countries, to evaluate the accuracy and geographic heterogeneity of molecular detection of common MTB genetic resistance determinants, and discuss implications for DR-TB control.

## Methods

Further details available in the supplementary material.

### Data and quality control

We compiled a 10,299 MTB WGS dataset with culture based DST (phenotypic) data using public databases (Patric^14^, ReSeqTB^15^) and literature curation^11–13, 16–26^. A summary table with the phenotypic data is available online at https://github.com/farhat-lab/resdata.

### Genomic analysis/variant calling

We used a previously validated genomic analysis pipeline for MTB described by Ezewudo *et al*.^27^ with modifications as detailed in the supplement.

### Drug resistance definitions

Drugs were labelled as follows: isoniazid (INH), ethambutol (EMB), rifamycins (rifampicin or rifabutin) (RIF), streptomycin (STR), pyrazinamide (PZA), fluoroquinolones (FLQ) (includes moxifloxacin, ciprofloxacin, ofloxacin), second-line injectables (SLIs) (includes kanamycin, amikacin, capreomycin), ethionamide/prothionamide (ETH), and cycloserine (CYS). Para-aminosalicylic acid was not analysed due to the paucity of data. Isolates not tested for susceptibility to both INH and RIF were excluded from the assessment of DR frequency by country and lineage. Isolates resistant to both INH and RIF were labelled MDR. Those resistant to INH, RIF, FLQ and SLIs were labelled XDR.

### Estimating resistance acquisition dates

Isolates were separated into 179 groups corresponding to a single drug, lineage and source country, referred to hereafter as a ‘group’. Genetic diversity was computed as the average pairwise genetic distance within a group. To accurately date resistance acquisition, a drug-geography-lineage group was analysed only if it consisted of at least 10 isolates, ≥20% of isolates were susceptible and ≥1 isolate was resistant. To exclude isolates that only represent outbreak settings and didn’t carry more long-term information about resistance, we excluded groups with a genetic diversity score <1 standard deviation from the mean genetic diversity score measured across all groups. Supplementary methods detail the phylogeny construction and the estimation of the age of the most recent *susceptible* common ancestor (MRSCA) in years prior to isolation of the clinical sample(s).

### Distribution of Resistance Mutations

We compared the expected sensitivity and specificity of mutations captured by commercial diagnostics (summarized from the literature in Table S9) and those based on more extensive lists of mutations in DR genes that can be captured using targeted or whole genome sequencing. We used three mutations lists for the latter (1) a set of 267 common resistance-associated mutations that we previously determined using randomForests^28^ designated “RF-select WGS test”, (2) a set of mutations determined using direct association^29^ designated “DA-select WGS test”, and (3) any non-synonymous mutation or noncoding mutation in known DR regions (Table S10) in a “all WGS test”. We excluded previously described neutral/lineage associated mutations^9^.

### Code

All code used in the analysis is publically accessible at https://github.com/farhat-lab/geo_dist_tb.

## Results

### Data and global lineage distribution

Of the 10,299 MTB clinical isolates with WGS and culture-based DST data available, 9,385 passed sequence quality criteria and of these 8,550 had country of origin data (Figure 1). The four major MTB lineages, 1-4, were well represented. A relatively high proportion, 42%, of United Kingdom (UK) isolates (n= 1873) belonged to Lineages-1 & 3 (Figure 2A). Overall, the non-Europe-America-Africa Lineages-1,3, and 2, comprised 40% of European isolates (n=3956) and 7% of North and South American isolates (n=1297).

**Figure 1:**
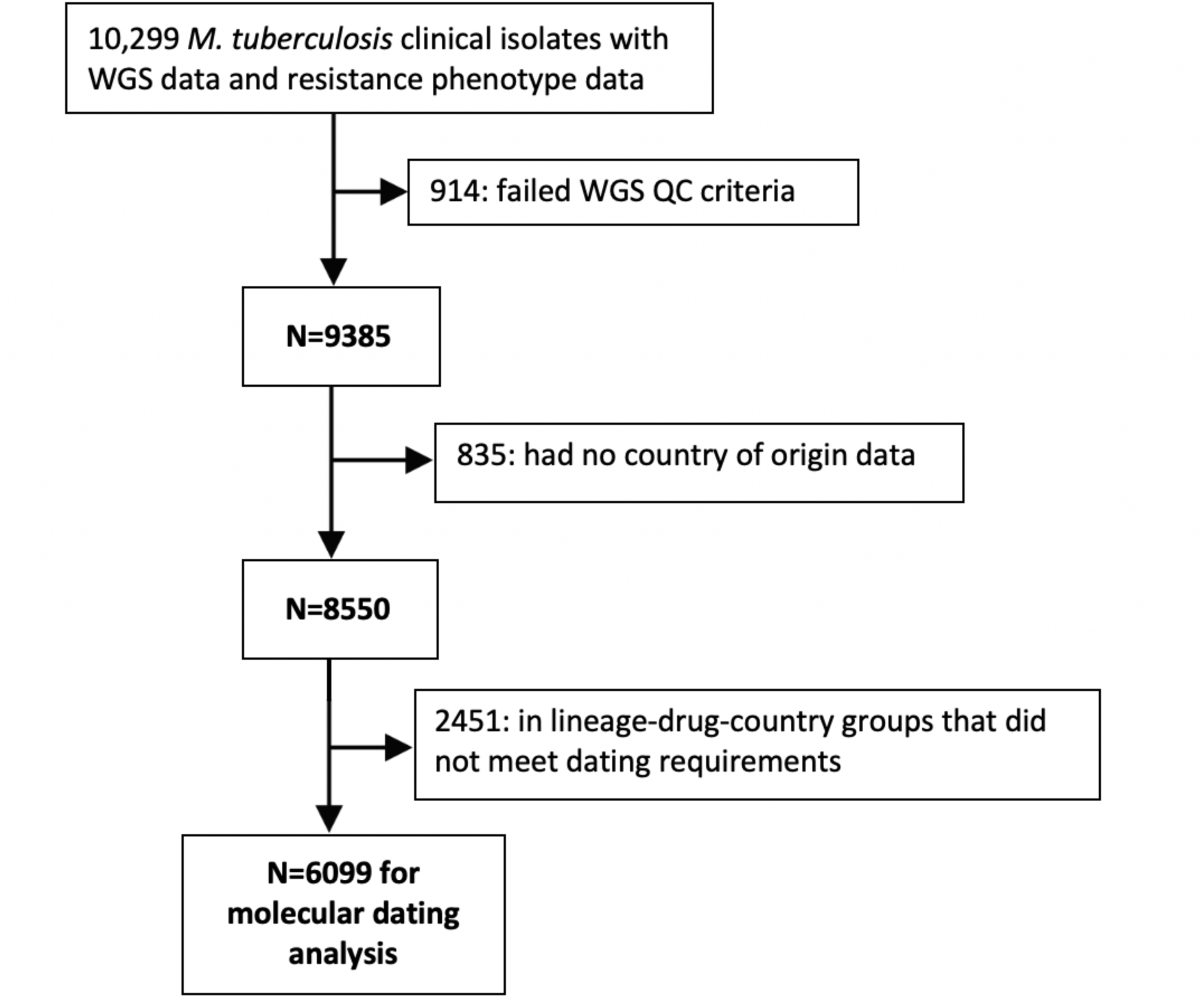
Flow diagram showing process of identification and exclusion of genomic data included in the study. WGS: Whole genome sequencing, QC: Quality control.

**Figure 2:**
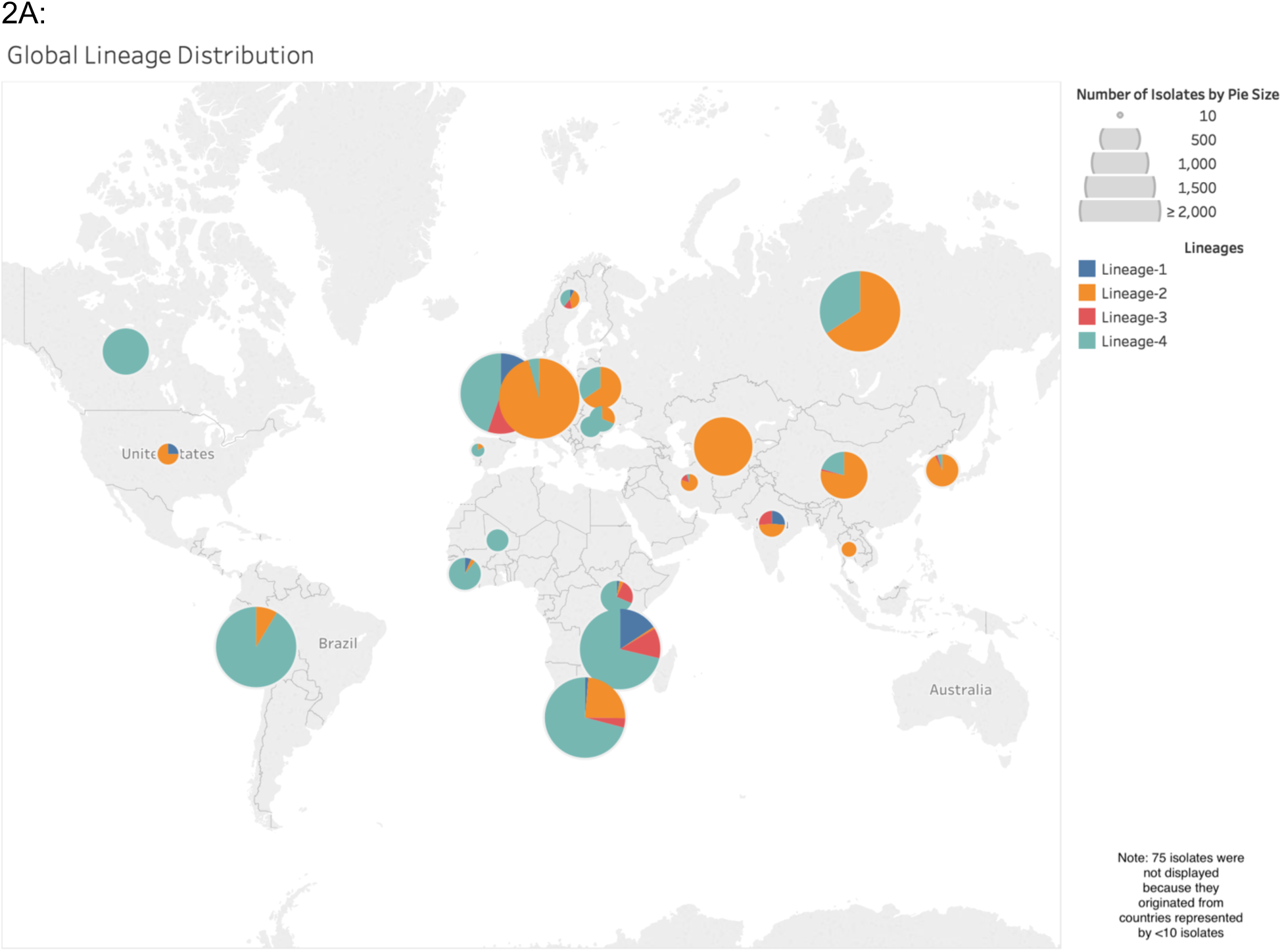
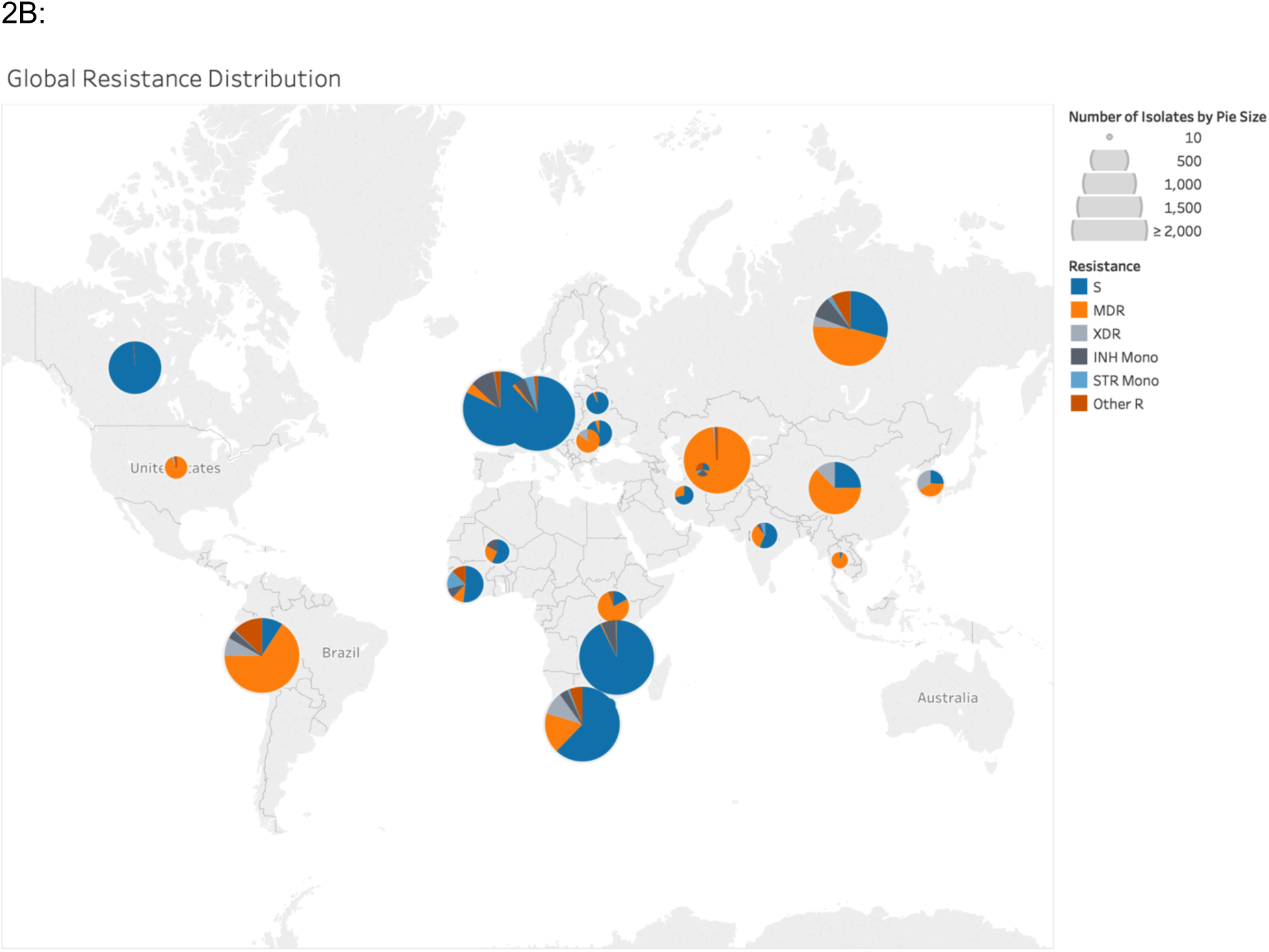
Global Distribution of *M. tuberculosis* in the study sample. Counts from countries represented by fewer than 10 isolates (n=75) not shown. **A:** Lineage distribution (n=8477). Pie charts represent the proportion of each lineage among isolates available from each country. Size of the pie is proportional to the number of isolates from each country. Detailed counts are in Table S1. **B:** Drug resistance distribution (n=7834). Pie charts provide the distribution of resistance patterns (S: Susceptible, MDR: Multidrug-resistant, XDR: Extensively drug resistant, INH Mono: mono resistant to isoniazid, STR Mono: mono resistant to streptomycin, Other R: resistance other than defined categories) by country. 75 isolates originated Pie size is proportional to the number of isolates in each country (Table S2).

**Table 1:** Sensitivity and Specificity of commercial and WGS based tests for resistance diagnosis. . Abbreviations defined in “drug resistance definitions” section of methods.

| Drug | commercial test |  | RF-select WGS test |  | DA-select WGS test |  | all WGS test* |  |
| --- | --- | --- | --- | --- | --- | --- | --- | --- |
|  | Sensitivity | Specificity | Sensitivity | Specificity | Sensitivity | Specificity | Sensitivity | Specificity |
| INH | 83% (2759/3306) | 93% (4834/5201) | 88% (2900/3306) | 92% (4780/5201) | 89% (2956/3306) | 92% (4776/5201) | 92% (3029/3306) | 64% (3314/5201) |
| RIF | 90% (2354/2624) | 93% (5361/5786) | 91% (2385/2624) | 92% (5341/5786) | 91% (2395/2624) | 92% (5338/5786) | 92% (2405/2624) | 89% (5178/5786) |
| FLQ | 53% (452/854) | 94% (2626/2790) | 51% (439/854) | 95% (2637/2790) | 51% (440/854) | 95% (2641/2790) | 57% (488/854) | 86% (2406/2790) |
| SLI | 56% (517/921) | 86% (2179/2547) | 58% (535/921) | 84% (2136/2547) | 57% (524/921) | 86% (2185/2547) | 64% (594/921) | 80% (2027/2547) |
| PZA |  |  | 65% (862/1324) | 95% (4660/4907) | 75% (996/1324) | 91% (4485/4907) | 78% (1030/1324) | 89% (4384/4907) |
| EMB |  |  | 79% (1476/1863) | 86% (4702/5441) | 75% (1389/1863) | 88% (4772/5441) | 85% (1589/1863) | 74% (4042/5441) |
| STR |  |  | 76% (1601/2112) | 86% (3204/3726) | 77% (1627/2112) | 87% (3223/3726) | 91% (1928/2112) | 71% (2657/3726) |
| Legend |  |  |  |  |  |  |  |  |
| * | classifying FLQ and SLI as susceptible if no INH and no RIF resistance mutation found |  |  |  |  |  |  |  |
| Sensitivity | Percent of resistant isolates classified as resistant |  |  |  |  |  |  |  |
| Specificity | Percent of susceptible isolates classified as susceptible |  |  |  |  |  |  |  |

### Phenotypic resistance distribution

Of the 8,550 isolates, 568 isolates lacked either INH or RIF DST data. Out of the remaining 7,909 isolates, 5,022 were pan-susceptible, 2887 were resistant to one or more drugs (DR) and of these 1937 were resistant to INH and RIF (MDR) and 288 were MDR and resistant to an SLI and a FLQ (XDR). The 8,550 isolates originated from 52 countries. Of these, 23 countries were represented by >10 isolates with resistance data, 21/23 were found to have MDR isolates and 9/21 had XDR (Figure 2B). We compared the MDR frequency in our WGS based sample with the WHO reported MDR/RIF resistance (RR) rates for the latest year available^30^. Out of the 21 countries, the confidence interval for the MDR-TB proportion in our sample overlapped with that of the WHO in 4 (19%) countries, was higher in 14 (67%) and lower in 3 (14%) of countries (Table S4). MDR rates by lineage were 3% for Lineage-1 (n=439), 48% for Lineage-2 (n=1085), 4% for Lineage-3 (n=760) and 23% for Lineage-4 (n=3358).

### Molecular dating of Resistance Acquisition

Of the 8,550 isolates, 2,451 isolates appeared in groups that did not meet our dating requirements (Methods). The remaining 6,009, included 1,547 isolates resistant to one or more drugs and were grouped into 179 country/lineage/drug combinations. We estimated 4,869 MRSCA dates for 10 drugs across these 179 groups. The number of independent resistance acquisition events *i.e.* unique MRSCA dates, was consistently lower than the total number of dated resistance isolates suggesting ongoing transmission of drug resistant isolates (Table S11). We estimated a lower bound on the burden of resistance due to transmission ranging by country from ≥14% to ≥52% pooled across drugs (Methods, Table S11). The proportion of INH or RIF resistance attributed to transmission was the highest among the 10 drugs at ≥43% and ≥46% respectively pooled across countries (pooled from Table S11).

We examined the relative order of phenotypic resistance acquisition on a global scale. For INH, we found that resistance to INH on average developed before resistance to other drugs (Figure 3A-B). Median MRSCA for INH was 11.4 years prior to isolation (IQR 6.3-16.2) *vs.* 7.6 years (IQR 3.0 – 16.0) for RIF, Wilcoxon P-value <10^-14^. Median MRSCA ages for RIF and STR resistance (7.6 years, IQR 3.0 – 16.0 and 7.7 years, IQR 3.4 – 13.0 respectively) were second oldest and not statistically significant from each other (Wilcoxon P-value 0.31). The dating supported that EMB resistance followed the acquisition of RIF (Wilcoxon P-value < 10^-6^) at a median MRSCA age of 5.0 years prior (IQR 2.1 – 12.5), and that this was followed by resistance to PZA, ETH, FLQ, SLIs, or CYS (Figure 3A), amongst which MRSCA ages did not significantly differ (Figure 3B). We found no significant correlation between the median MRSCA dates and the drug’s date of introduction into clinical use with *R^2^*= 0.04 (F-test with 1 DF P-value= 0.60, Table S12).

**Figure 3A:**
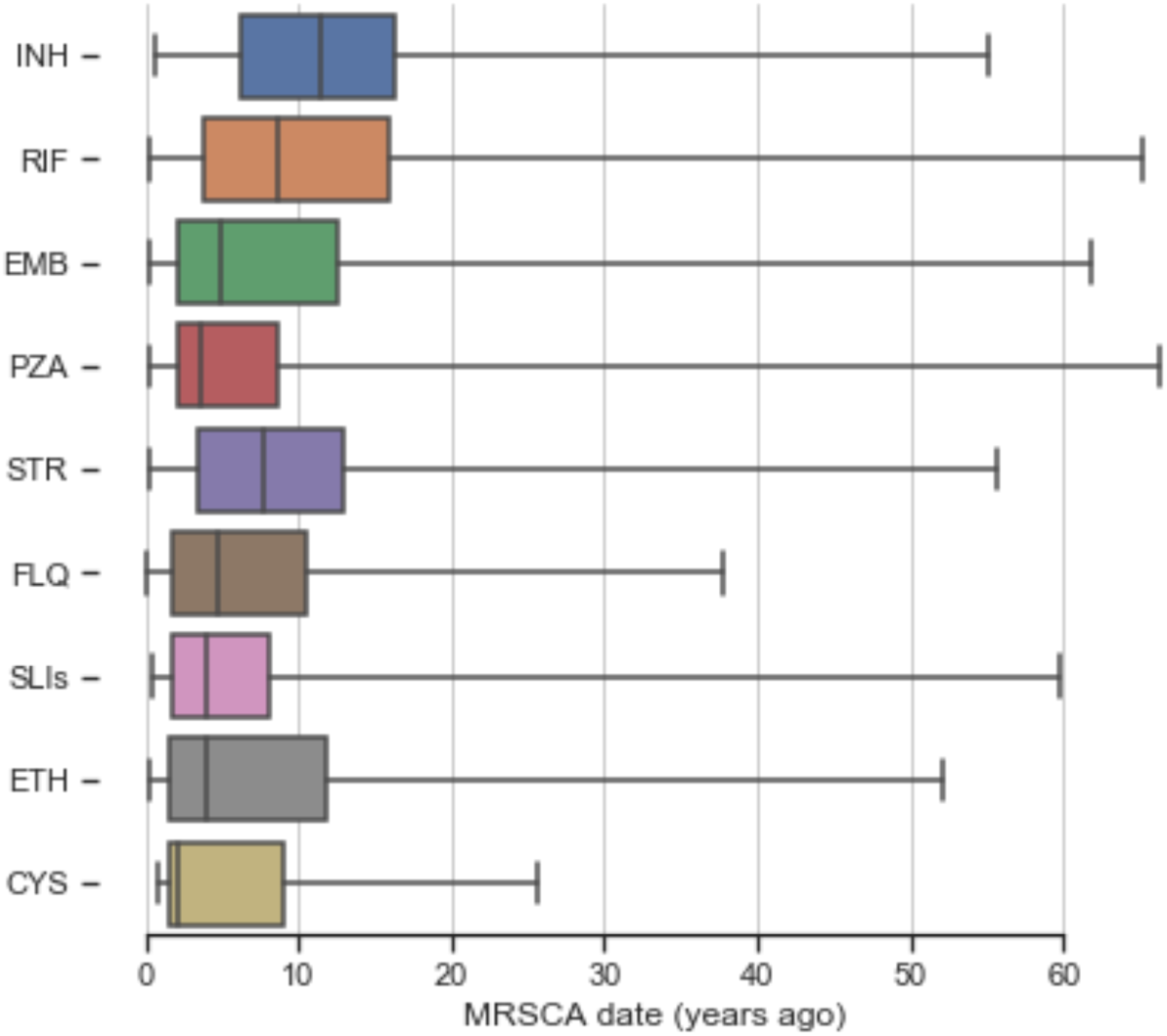
MRSCA distribution by drug (n=4844). Boxplots showing range of MRSCA distribution globally for nine anti-tubercular drugs (MRSCA: Most recent susceptible common ancestor, INH: isoniazid, RIF: rifampicin, EMB: ethambutol, PZA: pyrazinamide, STR: streptomycin, FLQ: fluoroquinolones, SLIs: second-line injectables, ETH: ethionamide, CYS: cycloserine)

**Figure 3B:**
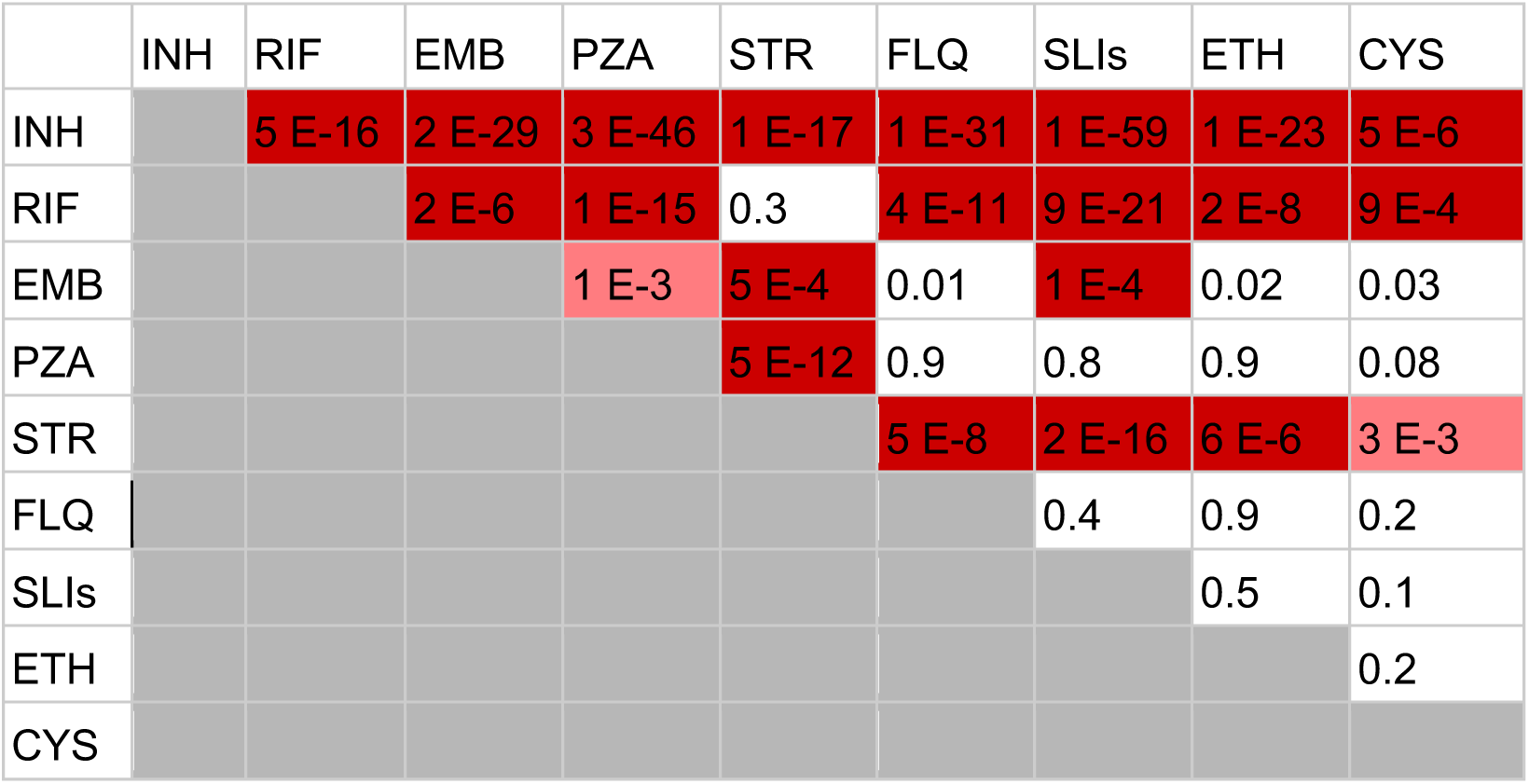
Pairwise Wilcoxon rank sum tests comparing MRSCA ages by drug category. Dark red indicates P-value <0.001 (Bonferroni threshold); pink indicates P-value <0.01; white indicates P-value ≥0.01.

We assessed the frequency of recent resistance amplification to PZA, EMB, FLQs and SLIs among MDR, *i.e.* to pre-XDR/XDR, within five years of sample isolation. Among the 11 countries with both MDR and pre-XDR/XDR isolates, we identified four countries (Peru, Russia, Sierra Leone, South Africa) with recent resistance amplification to PZA and EMB (>1% of MDR). The rates of recent amplification ranged from 2% (95% CI 1% - 4%) for PZA in Russia to 33% (95% CI 26% - 41%) for EMB in South Africa (Figure 5). Peru, Romania and South Africa were also measured to have recent resistance amplification to FLQs and SLIs (Figure 5). The median MRSCA age for FLQ or SLI resistance acquisition among MDR isolates was 4.7 years (IQR 1.9-9.8) prior to sample collection.

We found RIF to have the highest proportion of old resistance (MRSCA >20 years prior to isolation) at 17%, 197/1184 out of the total dated RIF resistance acquisition events. Old resistance was well distributed geographically and for RIF occurred in 9 of 12 countries with available dating data (Figure 4). Old FLQ resistance constituted 8% (24/311) of total dated isolates and spanned 6 of the 7 countries with available data.

**Figure 4:**
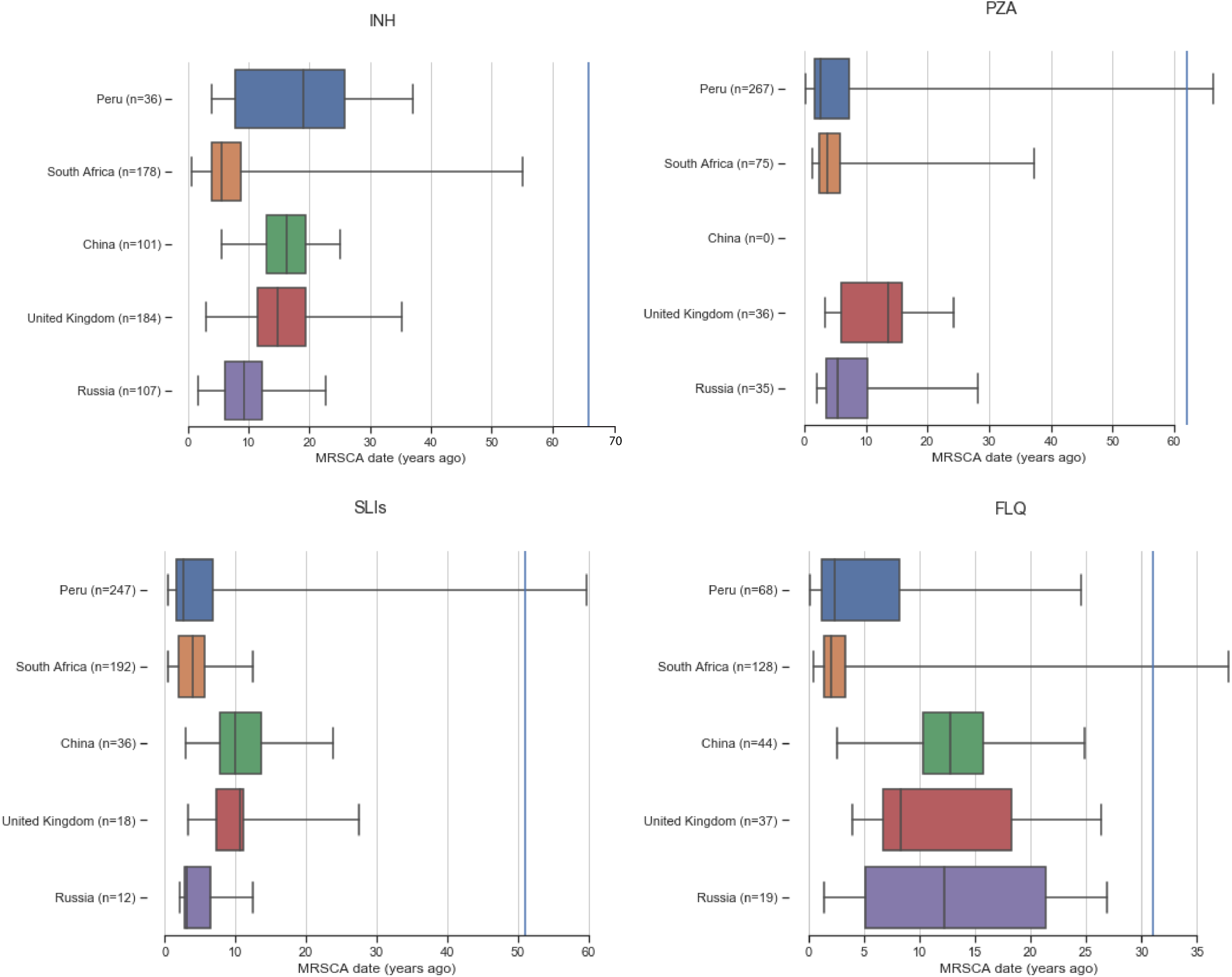
Box and whisker plots summarizing the MRSCA distribution per country. (Blue vertical line indicates year when drug was introduced (Table S5))

**Figure 5:**
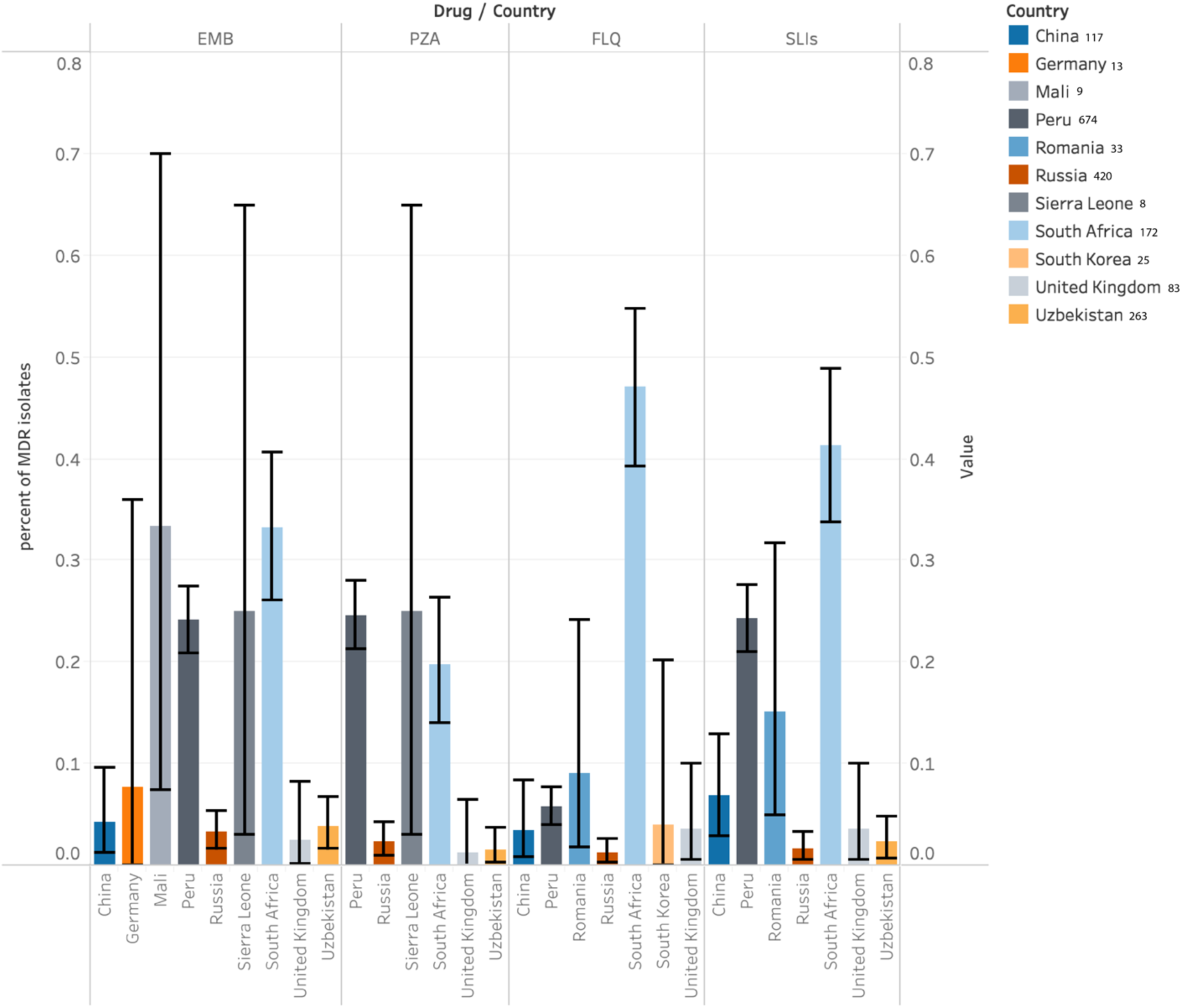
**Proportion of MDR isolates with recent amplification of resistance** to ethambutol (EMB), pyrazinamide (PZA), fluoroquinolones (FLQ) or second-line injectables (SLIs) by country (MRSCA age estimate <5 years ago). The legend lists the number of multidrug-resistant isolates analysed from each country. Error bars indicate 95% confidence intervals. Full data given in Supplementary Table 3. Four countries displayed a measurable proportion of recent FLQ and SLI amplification (95% CI does not include 0) – South Africa, Peru, Romania and China.

We compared the geographic distribution of MRSCA ages restricting to four key drug classes, namely INH, RIF, SLIs and FLQs, and the five countries with the largest number of resistant isolates (Figure 4). MRSCA ages did not differ between the UK and China across all four drug classes. These two countries had the oldest median MRSCA across the five countries and four drug classes except for INH. MRSCA ages in the UK were a median of 13.0 years (IQR 10.7-19.0) for RIF, a median of 10.6 years (IQR 7.3-11.2) for SLIs and a median of 8.4 years (IQR 6.7-18.4) for FLQs (Figure 4). South Africa most consistently had the youngest median MRSCA for the four drug classes, but its MRSCA distribution was not significantly different from that of Peru (for FLQs and SLIs) and Russia (for SLIs) (Table S3). A similar geographic/age pattern was observed for the drugs PZA and EMB across these five countries (Table S3).

We examined if the geographic resistance age differences correlated with resources available for TB control programs using the gross domestic product per capita as a proxy. We found GDP to correlate significantly with an older RIF MRSCA date with *R^2^*= 0.47 (F-test with 1 DF P-value= 0.014) (Figure 6 & Table S8).

**Figure 6:**
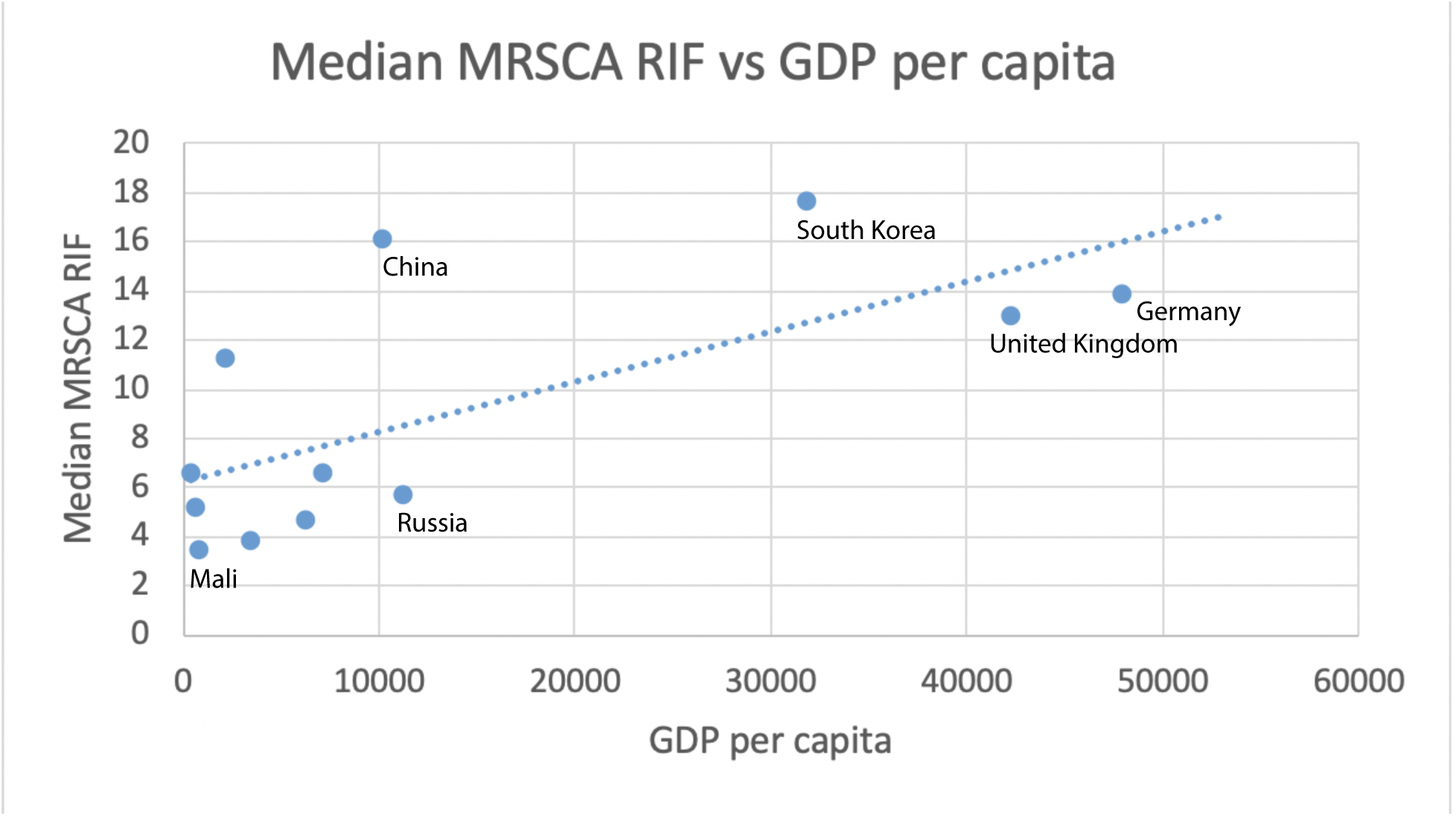
Median Rifamycin (RIF) most-recent-common-susceptible-ancestor (MRSCA) date vs Gross Domestic Product (GDP) per capita for 12 countries. Data plotted is provided in Supplementary Table 7 and includes other drugs than RIF. (*R^2^* = 0.47, F-test P-value (1 DF) =0.014).

### Distribution of Resistance Mutations

We assessed the frequency of 267 resistance mutations previously determined to be important for resistance prediction^28^ and their geographic distribution among the 8,550 isolates with country of origin and WGS data meeting quality criteria (Figure 1). Resistance mutation prevalence varied significantly by country. The most frequent INH causing mutation^31^, *katG* S315T was more frequent among phenotypically INH resistant isolates by DST (pheno-R) in Russia (84%, n=526) than in Peru (67%, n=760) (Fisher P-value 1×10^-12^). The second most common INH resistance mutation −15 C>T *fabG1/inhA* promoter was more prevalent among INH pheno-R Peruvian isolates (20%) than in Russian isolates (8%) (Fisher P-value 7×10^-9^). Twenty four of the 267 resistance mutations (9%) varied geographically to a larger extent than the mutation *fabG/inhA* promoter −15C>T (standard deviation 11%, frequency range 0-39%, Table S13). The mutation I491F was recently described to be common in Estawini^32^ and is not detectable by line-probe or GeneXpert commercial molecular diagnostics. In our sample that did not contain data from Estawini, we calculated a standard deviation of 1% for the global frequency of I491F (range 0% - 4%) among RIF pheno-R isolates.

We calculated the proportion of pheno-R isolates that can be captured by the Hain Line-probe or GeneXpert commercial molecular diagnostics due to the presence of one or more mutations in their pooled target regions for the drugs INH, RIF, SLIs, and FLQ (Tables 1 and S14). Sensitivity was highest for RIF (90% of 2624) and lowest for FLQs (51% of 854). Specificity was consistently high (lowest for SLIs at 86%) (Table 1). Second-line sensitivity of commercial diagnostics differed significantly across countries (Table S14). FLQ sensitivity in Peru was 38% (n= 121) and 77% in South Africa (n=111) (Fisher P-value 1×10^-6^). A similarly low sensitivity for SLI resistance was seen in Peru compared with South Africa (Fisher P-value 3×10^-4^) (Table S14).

We examined if expanding the resistance mutation list to variants previously characterized in diverse global MTB genomic datasets using direct association^29^ or random forests^28^ can improve sensitivity and specificity in a “select WGS test.” The select WGS test improved sensitivity slightly for INH and SLIs with relatively preserved specificity (Table 1). In addition select WGS test allowed for prediction of resistance to other drugs not tested by commercial diagnostics: PZA, EMB and Streptomycin. For comparison, we assessed if including any non-silent variant in the resistance regions (excluding a select number of known lineage markers) was indeed inferior to the more informed ‘select WGS test” reported previously. We found that this “all WGS test” only modestly improved sensitivity and at the expense of a larger decrease in specificity (Figure 7).

## Discussion

Using 8,550 clinical MTB sequences with culture-based DST, we examined geographic trends in the DR-TB epidemic. Geographically, MTB lineages 1-4 were each represented in the continents of Europe, Asia and Africa providing evidence of disease spread across borders, likely driven by human migration^30^. We found MDR-TB in nearly every country represented by more than 10 isolates. XDR isolates were found in half of these countries and spanning all five major continents. Lineage-2 had the highest percentage of MDR isolates in our sample followed by Lineages-4, 3 and 1. Using this diverse sample we dated more than 4869 resistance phenotypes across 4 lineages and 15 countries.

We found a consistent order of resistance acquisition globally among drug classes. The development of INH resistance was previously found to be a sentinel event heralding the development of MDR^33^. Our results corroborate these findings using phenotypic resistance data and across a larger geographically diverse sample. After INH, we find that MTB acquires resistance to RIF/STR then EMB followed by PZA, ETH, FLQ, SLIs, or CYS. We found no correlation between the age of resistance acquisition and the year of clinical introduction of the drug but there may be multiple other causes for the observed order of resistance acquisition. Differences in mutation rates across drug targets or resistance genes have been postulated but shown to be an unlikely explanation for INH resistance arising first^34, 35^. Pharmacokinetic difference may result in higher risk for under-dosing^36^ for some drugs and earlier resistance acquisition. Bacterial fitness costs are also variable across resistance mutations. For INH resistance, mutations like *katG* S315T carry a low fitness cost and likely contribute to resistance arising earliest for this drug^33, 37, 38^. The order of drug administration can explain dating differences between first-line (INH, RIF, EMB, PZA) and second (ETH, FLQ, SLIs) or third-line (CYS) resistance, as second-line drugs are usually only administered after resistance to first-line drugs is ascertained. Acquisition of resistance to INH first then RIF may also relate to their use for treatment of latent TB infection, leading to more exposure and selection pressure overall. However, because adoption of INH preventative therapy for latent TB remains low in many parts of the world, we expect it to be a lesser contributor to INH and RIF resistance rates^39^. Lastly, the observation of contemporaneous acquisition of RIF and STR resistance is likely best explained by the effects of Category II TB treatment initially recommended in 1991^40^. Category II is no longer recommended by the WHO but consists of adding streptomycin to the first-line drug regimen after treatment failure. Our dating supports that streptomycin resistance amplified among patients failing due to recent RIF resistance and/or MDR acquisition.

Published evidence supports that most resistant cases of MTB result from recent resistance acquisition in the host or are related to transmission^41^. Reactivation of resistant MTB disease acquired remotely (>2 years prior) is much less likely^42^. Thus the identification of isolates with old resistance suggests high fitness for continued transmission between human hosts. Most countries with available data had isolates with resistance dated more than 20 years prior. This is also supported by our phylogenetic assessment where we estimate a lower bound of TB resistance due to transmission to range between 14-52% across countries with available data. As our approach cannot distinguish between resistance importation through human migration after transmission outside of the country and new resistance acquisition, these figures are underestimates of the true resistance burden due to transmission. Mathematical models of TB rates have previously predicted transmission to be a major driver of observed resistance rates^43^, we present here WGS based evidence of the high burden of resistance transmission. Mathematical models have also emphasized that drug resistant strain fitness is a key parameter that dictates how the resistance epidemic will unfold. Our results support that >14-52% of isolates are fit and successfully transmitting patient-to-patient and in most countries there have been uninterrupted chains of resistance transmission for >20 years. These data emphasize the need to contain resistance transmission through improved diagnosis, treatment and other preventative strategies such as infection control and vaccine development.

In addition to transmission, we find evidence for recent resistance amplification, especially to second-line drugs mediating the transition from MDR to XDR-TB. XDR has considerably worse treatment outcome than susceptible TB and incurs more than 25 times the cost^44^. We estimate that half of FLQ and SLI resistant isolates had acquired resistance within 4.7 years of isolation despite the promotion of directly-observed-therapy (DOT) by the WHO since 1994. As most FLQ and SLI resistant isolates are also MDR, our results also emphasize the need for better regimens to treat MDR that can prevent resistance amplification. By country, we found a significant correlation between the estimated age of resistance acquisition and per capita GDP, with more affluent countries having older ages of resistance. This unexpected correlation is likely driven by a combination of factors but the routine use of DST and close patient monitoring in the more well-resourced health systems are likely important contributors. Specifically, we found the UK and China to have the oldest resistance ages across the drugs. The Chinese national TB program budget was increased from $98 million in 2002 to $272 million in 2007^45^ and a new policy for free TB diagnostics tests and drug use was introduced in 2004^58, 46^. This increased investment can explain the observed low rates of recent resistance acquisition in China^30, 47^.

Likely due to geographic differences in MTB lineage, transmission and resistance acquisition rates, we find 10% of assayed resistance mutations to have high geographic variance. We also found commercial diagnostics to vary in sensitivity for second-line drugs. Given recent reports about the accuracy of WGS for confirming susceptibility of MTB^29^, we measured improvements in resistance sensitivity offered by including mutations outside of regions targeted by commercial diagnostics through direct association. This offered modest improvements in sensitivity with little to no change in specificity. We found a considerable number of indeterminate mutations in resistance regions, that when included with simple direct association improve sensitivity but at the expense of loss of specificity. The study of these variants through statistical models will likely further inform their diagnostic use in the future^28, 48^.

Our study has several limitations including the oversampling of DR isolates as evidenced by our comparison with WHO reported MDR rates. We tried to control for this by dating only in countries with at least 20% susceptible isolates and limiting dating of low diversity samples that represent unique outbreak settings and lack long term information about resistance acquisition. This may have resulted in underestimation of rates of recent resistance acquisition but despite this we were able to document recent resistance acquisition in many countries. Molecular dating is also reliant on the accurate estimation of the phylogenetic tree of MTB isolates and the molecular clock assumption. We thus used a rigorous approach to phylogenetic estimation and dating despite its computational and time cost^49^. Our analysis also assumes the accuracy of culture based phenotypic DST, even though test to test variability is known to exist. We justify this as our data was curated from ReseqTB^15^ and studies were phenotypic testing was performed in national or supranational laboratories with rigorous quality control.

Overall, our results support that DR rates are fuelled by both recent resistance acquisition and ongoing transmission, and suggest the need for better detection, treatment and health system investment. In the future, reassessment of these patterns will be enabled by the sharing of systematically collected isolate data, data that is increasingly generated as by-products of TB surveillance and resistance diagnosis^29^.

## Authors’ contributions

Yasha Ektefaie conducted the data analysis, drafted and revised the manuscript.

All authors provided key edits to the manuscript.

Additionally:

Luca Freschi contributed to the data analysis.

Avika Dixit contributed to the data analysis.

Maha Farhat conceptualized the study, supervised the data analysis, reviewed, wrote and edited the manuscript.

## Supplementary Methods

### Data curation and quality control

The isolate metadata including their geographic locations were downloaded using metatools_ncbi (https://github.com/farhat-lab/metatools_ncbi). We also performed literature curation to fill the gaps in the NCBI geographic location data. The resulting table of the geographic locations of the isolates is available in Supplementary File 1. We excluded isolates that did not meet WGS quality control criteria as detailed below, had no geographic information or were not tested for phenotypic resistance to one or more drugs.

### Genomic analysis/variant calling

Briefly, reads were trimmed using PRINSEQ^1^ setting average phred score threshold to 20. Raw read data was confirmed to belong to MTB complex using Kraken^2^. Isolates with <90% mapping were excluded. Reads were aligned to H37Rv (GenBank NC000962.3) reference genome using BWA MEM^3^. Duplicate reads were removed using PICARD^4^. We excluded any isolates with coverage <95% of known drug resistance regions (*katG*, *inhA* & its promoter, *rpoB*, *embA, embB, embC* & *embB* promoter, *ethA*, *gyrA* and *gyrB*, *rrs*, *rpsL*, *gid*, *pncA*, *rpsA*, *eis* promoter) at 10x or higher. Variants were called using Pilon^5^ that uses local assembly to increase indel (insertions and deletions) call accuracy. This deviates from Ezewudo *et al.* that uses Samtools for variant calling^6^. The reference allele was implied if allele frequency was <75% or the Pilon filter was not PASS. Low confidence coordinates were filtered from all strains if >95% of strains did not have coverage of (trimmed reads) at least 10x at that site. Isolate lineage belonging to one of the seven main MTB lineages was confirmed using the Coll *et al.* SNP barcode^7^.

### Drug resistance definitions and comparison with WHO reported resistance proportions

The ‘Mono’ resistant designation was given to isolates that were resistant to only one specific drug and susceptible to all others that were tested, with the exception of the INH-mono resistant category that encompassed any isolates that were resistant to INH and/or STR but not to others that were tested. The ‘Other-R’ category was reserved for isolates that were resistant to some drugs but were neither INH or STR mono resistant, nor MDR or XDR. Isolates were labelled susceptible if they were susceptible to all drugs tested.

To compute exact binomial confidence intervals for MDR proportions by country we used the python library statsmodel^8^. To assess overlap with World Health Organization (WHO) estimates we determined whether the confidence intervals of our proportion intersected with that of the WHO. We labelled our estimate as high if our confidence interval was higher than the WHO, low if it was lower, and the same if they intersected.

### Estimating resistance acquisition dates and lower bounds of resistance transmission

A maximum likelihood tree was generated for each group via RAxML 8.2.11^9^ with H37Rv (NC000962.3) as the outgroup, starting from a neighbour-joining seed tree and assuming a generalized time reversible (GTR) nucleotide substitution model with the Γ distribution used to model site rate heterogeneity^10^. We bootstrapped the maximum likelihood tree 1000 times. The maximum likelihood tree was dated using BEAST v1.10.4^11^ assuming a relaxed molecular clock with a log normal distribution and a mean rate of 0.5 SNP per genome per year based on prior published data^12^. Sumtrees.py from the DendroPy library^13^ was then used to combine the output from the bootstrap analysis and that of BEAST to get our final dated phylogenetic tree with nodal bootstrap support.

We dated the most recent common ancestor between all the resistant isolates and their most closely related susceptible isolate. Accordingly, the dates of resistance acquisition will be referred to as the estimated age of the most recent *susceptible* common ancestor (MRSCA) in years prior to isolation of the clinical sample(s) throughout the text. We excluded resistant isolates with MRSCAs inferred at nodes with less than 50 bootstrap support.

We calculated the number of phylogenetically inferred resistance acquisition events (*N_aq_*) per country and lineage as the number of unique MRSCAs identified. This was compared with the total resistant isolates that could be dated (*N_td_*). Phylogenetically inferred unique resistance acquisition events for a particular country may be related to either *in host* evolution of new resistance or due to human migration/importation from another country with the latter still possibly related to transmission elsewhere. Thus, the following quantity represents a lower bound on the burden of resistance due to transmission for a particular country:

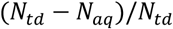

To estimate the order of resistance acquisition for different drugs we pooled the MRSCA dates by drug across countries and lineages. We compared the medians of the MRSCA distributions and performed pairwise Wilcoxon Rank Sum tests to assess for statistical significance, correcting for multiple testing using the Bonferroni approach. Interquartile ranges were calculated using the python package numpy^14^.

We correlated the median MRSCA date per drug pooled across countries against the date of drug introduction using linear regression as implemented in Microsoft Excel version 16.25.

### Distribution of resistance mutations

We measured resistance mutations’ geographic variance by calculating the proportion of resistant isolates with the resistance mutation per country, excluding countries with less than 10 resistant isolates for a particular drug. We computed the standard deviation of this distribution of proportions across countries. To test cross-country differences in the proportion of INH phenotypically resistant isolates that contained specific mutations, we used a Fisher-test using the python library scipy^15^.

### GDP/Programmatic Spending

To correlate countries’ gross domestic product (GDP) per capita against resistance acquisition dates, GDP per capita data for 2019 was gathered from the International Monetary Fund (IMF)^16^ and plotted against the median MRSCA date for RIF using Microsoft Excel version 16.25. F-value and significance was calculated via Anova^17^ in Excel.

## Supplementary Material

**Supplementary Table 1.** Global Lineage Distribution.

|  | Lineage-1 | Lineage-2 | Lineage-3 | Lineage-4 | Lineage-5 | Lineage-6 | Lineage-7 | Total |
| --- | --- | --- | --- | --- | --- | --- | --- | --- |
| Belarus | 0 | 87 | 0 | 46 | 0 | 0 | 0 | 135 |
| Canada | 0 | 0 | 0 | 17 | 0 | 0 | 0 | 165 |
| China | 0 | 132 | 2 | 35 | 0 | 0 | 0 | 170 |
| Germany | 0 | 22 | 0 | 1 | 0 | 0 | 0 | 857 |
| India | 5 | 9 | 5 | 0 | 0 | 0 | 0 | 52 |
| Iran | 0 | 16 | 3 | 1 | 0 | 0 | 0 | 22 |
| Malawi | 208 | 10 | 162 | 945 | 0 | 0 | 0 | 1427 |
| Mali | 0 | 0 | 0 | 37 | 0 | 0 | 0 | 37 |
| Moldova | 0 | 16 | 0 | 35 | 0 | 0 | 0 | 51 |
| Netherlands | 7 | 19 | 0 | 70 | 0 | 0 | 0 | 98 |
| Peru | 0 | 82 | 0 | 869 | 0 | 0 | 0 | 1098 |
| Portugal | 0 | 2 | 0 | 10 | 0 | 0 | 0 | 13 |
| Romania | 0 | 0 | 0 | 33 | 0 | 0 | 0 | 33 |
| Russia | 0 | 525 | 1 | 273 | 0 | 0 | 0 | 868 |
| Sierra Leone | 3 | 2 | 0 | 41 | 4 | 10 | 0 | 79 |
| South Africa | 8 | 144 | 23 | 427 | 0 | 0 | 0 | 974 |
| South Korea | 0 | 37 | 1 | 2 | 0 | 0 | 0 | 80 |
| Swaziland | 0 | 9 | 0 | 0 | 0 | 0 | 0 | 10 |
| Sweden | 2 | 11 | 4 | 11 | 0 | 0 | 0 | 28 |
| Thailand | 0 | 2 | 0 | 0 | 0 | 0 | 0 | 17 |
| Turkmenistan | 0 | 10 | 0 | 0 | 0 | 0 | 0 | 11 |
| Uganda | 2 | 3 | 18 | 50 | 0 | 0 | 0 | 80 |
| United Kingdom | 222 | 102 | 560 | 716 | 9 | 4 | 1 | 1873 |
| United States of America | 1 | 3 | 0 | 0 | 0 | 0 | 0 | 34 |
| Uzbekistan | 0 | 20 | 0 | 0 | 0 | 0 | 0 | 265 |
| Unknown | 37 | 81 | 35 | 792 | 1 | 0 | 0 | 1749 |
| **Total** | **495** | **1344** | **814** | **4411** | **14** | **14** | **1** | **10226** |

**Supplementary Table 2.** Global Resistance Distribution and Other Resistance Count.

| country | S | MDR | XDR | INH_MONO | STR_MONO | OTHER_R | NO_DATA | SUM | SUM_W_DATA | MDR/OTHER_R |
| --- | --- | --- | --- | --- | --- | --- | --- | --- | --- | --- |
| Belarus | 28 | 1 | 0 | 1 | 0 | 0 | 105 | 135 | 30 | NA |
| Canada | 163 | 0 | 0 | 2 | 0 | 0 | 0 | 165 | 165 | NA |
| China | 46 | 117 | 23 | 0 | 0 | 0 | 7 | 170 | 163 | NA |
| Germany | 749 | 13 | 0 | 40 | 34 | 14 | 7 | 857 | 850 | 0.928571429 |
| India | 22 | 13 | 0 | 2 | 2 | 0 | 13 | 52 | 39 | NA |
| Iran | 15 | 6 | 0 | 0 | 0 | 0 | 1 | 22 | 21 | NA |
| Malawi | 1302 | 7 | 0 | 90 | 1 | 3 | 24 | 1427 | 1403 | 2.333333333 |
| Mali | 20 | 9 | 0 | 6 | 0 | 0 | 2 | 37 | 35 | NA |
| Moldova | 38 | 2 | 0 | 0 | 0 | 0 | 11 | 51 | 40 | NA |
| Netherlands | 68 | 0 | 0 | 5 | 20 | 0 | 5 | 98 | 93 | NA |
| Not Provided | 179 | 646 | 138 | 90 | 6 | 109 | 719 | 1749 | 1030 | 5.926605505 |
| Peru | 94 | 674 | 86 | 34 | 5 | 134 | 157 | 1098 | 941 | 5.029850746 |
| Romania | 0 | 33 | 6 | 0 | 0 | 0 | 0 | 33 | 33 | NA |
| Russia | 261 | 420 | 42 | 80 | 17 | 76 | 14 | 868 | 854 | 5.526315789 |
| Sierra Leone | 41 | 8 | 0 | 7 | 13 | 10 | 0 | 79 | 79 | 0.8 |
| South Africa | 612 | 172 | 102 | 34 | 8 | 56 | 92 | 974 | 882 | 3.071428571 |
| South Korea | 16 | 25 | 21 | 0 | 0 | 0 | 39 | 80 | 41 | NA |
| Swaziland | 8 | 1 | 0 | 0 | 0 | 1 | 0 | 10 | 10 | 1 |
| Thailand | 1 | 15 | 0 | 0 | 0 | 0 | 1 | 17 | 16 | NA |
| Turkmenistan | 3 | 1 | 0 | 3 | 1 | 3 | 0 | 11 | 11 | 0.333333333 |
| Uganda | 10 | 45 | 0 | 1 | 0 | 2 | 22 | 80 | 58 | 22.5 |
| United Kingdom | 1525 | 83 | 4 | 187 | 5 | 50 | 23 | 1873 | 1850 | 1.66 |
| United States of America | 0 | 29 | 1 | 0 | 0 | 1 | 4 | 34 | 30 | 29 |
| Uzbekistan | 0 | 263 | 3 | 1 | 0 | 1 | 0 | 265 | 265 | 263 |
| Total | 5201 | 2583 | 426 | 583 | 112 | 460 | 1246 | 10185 | 8939 | 5.615217391 |

Abbreviations: RIF (Rifamycin), ETH (Ethionamide), FQ (Fluoroquinolones), SLIS (Second Line Injectable), “\_” (AND—EX: RIF\_ISONIAZID\_ETH = Isolate resistant to Rifamycin, Isoniazid, and Ethionamide)

| Other Resistance Categories | Count |
| --- | --- |
| RIF_ | 78 |
| PYRAZINAMIDE_ | 65 |
| ETHAMBUTOL_ | 44 |
| ISONIAZID_ETHAMBUTOL_STREPTOMYCIN_ | 29 |
| ISONIAZID_ETHAMBUTOL_ | 28 |
| ISONIAZID_PYRAZINAMIDE_ | 21 |
| ISONIAZID_PYRAZINAMIDE_STREPTOMYCIN_ | 16 |
| ISONIAZID_ETHAMBUTOL_PYRAZINAMIDE_STREPTOMYCIN_ | 11 |
| RIF_ETHAMBUTOL_ | 11 |
| RIF_STREPTOMYCIN_ | 10 |
| ISONIAZID_STREPTOMYCIN_ETH_ | 9 |
| ISONIAZID_ETHAMBUTOL_FQ_ | 9 |
| SLIS_ | 8 |
| ISONIAZID_STREPTOMYCIN_FQ_ | 8 |
| ETHAMBUTOL_FQ_ | 7 |
| FQ_ | 7 |
| ISONIAZID_ETHAMBUTOL_STREPTOMYCIN_FQ_ | 7 |
| ETHAMBUTOL_STREPTOMYCIN_ | 6 |
| ISONIAZID_ETH_ | 6 |
| STREPTOMYCIN_FQ_ | 5 |
| ISONIAZID_ETHAMBUTOL_PYRAZINAMIDE_ETH_ | 5 |
| ISONIAZID_FQ_ | 4 |
| ISONIAZID_ETHAMBUTOL_STREPTOMYCIN_ETH_ | 4 |
| ISONIAZID_ETHAMBUTOL_PYRAZINAMIDE_STREPTOMYCIN_SLIS_ETH_ | 3 |
| ISONIAZID_ETHAMBUTOL_PYRAZINAMIDE_STREPTOMYCIN_SLIS_ | 3 |
| ISONIAZID_PYRAZINAMIDE_ETH_ | 3 |
| ETHAMBUTOL_PYRAZINAMIDE_ | 3 |
| RIF_PYRAZINAMIDE_STREPTOMYCIN_ | 2 |
| RIF_ETHAMBUTOL_PYRAZINAMIDE_STREPTOMYCIN_FQ_ | 2 |
| ISONIAZID_ETHAMBUTOL_PYRAZINAMIDE_STREPTOMYCIN_ETH_ | 2 |
| PYRAZINAMIDE_STREPTOMYCIN_ | 2 |
| RIF_ETHAMBUTOL_STREPTOMYCIN_ | 2 |
| ISONIAZID_SLIS_ | 2 |
| PYRAZINAMIDE_SLIS_ | 2 |
| ISONIAZID_ETHAMBUTOL_PYRAZINAMIDE_SLIS_ETH_ | 2 |
| ISONIAZID_ETHAMBUTOL_PYRAZINAMIDE_ | 2 |
| PYRAZINAMIDE_FQ_ | 1 |
| ISONIAZID_PYRAZINAMIDE_FQ_SLIS_ | 1 |
| STREPTOMYCIN_SLIS_ | 1 |
| RIF_ETHAMBUTOL_FQ_ | 1 |
| ISONIAZID_ETHAMBUTOL_STREPTOMYCIN_SLIS_ETH_ | 1 |
| ISONIAZID_ETHAMBUTOL_STREPTOMYCIN_SLIS_ | 1 |
| RIF_ETHAMBUTOL_STREPTOMYCIN_SLIS_ETH_ | 1 |
| ETH_ | 1 |
| RIF_PYRAZINAMIDE_FQ_ | 1 |
| RIF_ETHAMBUTOL_PYRAZINAMIDE_FQ_ | 1 |
| RIF_ETHAMBUTOL_PYRAZINAMIDE_STREPTOMYCIN_ETH_ | 1 |
| RIF_PYRAZINAMIDE_ | 1 |
| ETHAMBUTOL_ETH_ | 1 |
| ISONIAZID_ETHAMBUTOL_ETH_ | 1 |
| RIF_ETHAMBUTOL_SLIS_ | 1 |
| ISONIAZID_PYRAZINAMIDE_STREPTOMYCIN_ETH_ | 1 |
| RIF_PYRAZINAMIDE_SLIS_ETH_ | 1 |
| RIF_ETHAMBUTOL_PYRAZINAMIDE_ | 1 |
| RIF_ETHAMBUTOL_PYRAZINAMIDE_STREPTOMYCIN_ | 1 |
| ISONIAZID_ETHAMBUTOL_FQ_SLIS_ | 1 |
| ETHAMBUTOL_STREPTOMYCIN_FQ_ | 1 |
| PYRAZINAMIDE_SLIS_ETH_PAS_ | 1 |
| ISONIAZID_STREPTOMYCIN_FQ_SLIS_ETH_PAS_ | 1 |
| ISONIAZID_STREPTOMYCIN_SLIS_PAS_ | 1 |
| ISONIAZID_ETHAMBUTOL_SLIS_ | 1 |
| RIF_ETHAMBUTOL_PYRAZINAMIDE_FQ_SLIS_ETH_ | 1 |
| RIF_FQ_SLIS_ | 1 |
| ISONIAZID_PYRAZINAMIDE_SLIS_ | 1 |
| ISONIAZID_ETHAMBUTOL_PYRAZINAMIDE_SLIS_ETH_PAS_ | 1 |
| RIF_ETHAMBUTOL_PYRAZINAMIDE_SLIS_ | 1 |
| RIF_PYRAZINAMIDE_STREPTOMYCIN_FQ_SLIS_ | 1 |
| ISONIAZID_ETHAMBUTOL_PYRAZINAMIDE_SLIS_ | 1 |
| RIF_SLIS_ | 1 |
| RIF_FQ_ | 1 |

**Supplementary Table 3.**
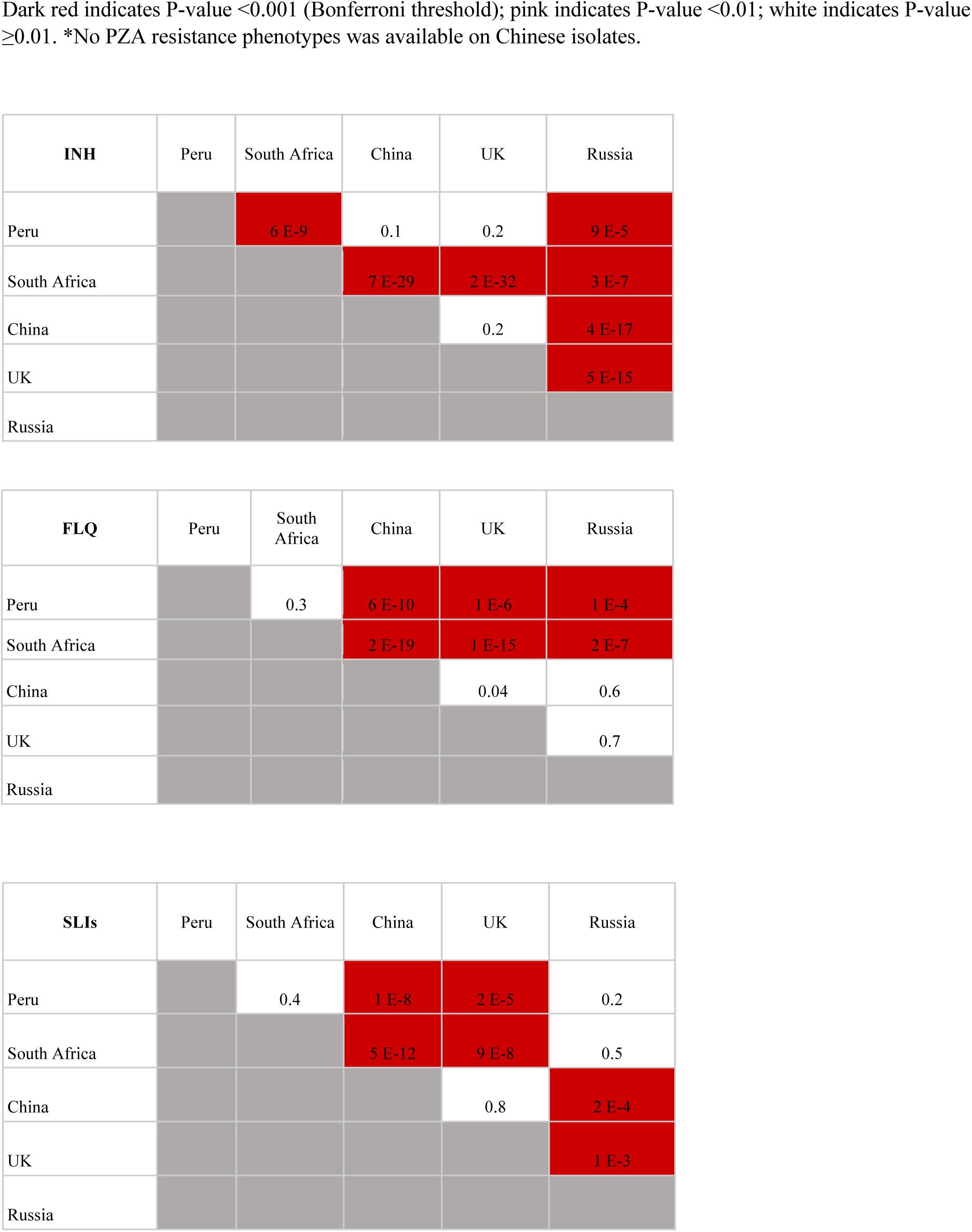
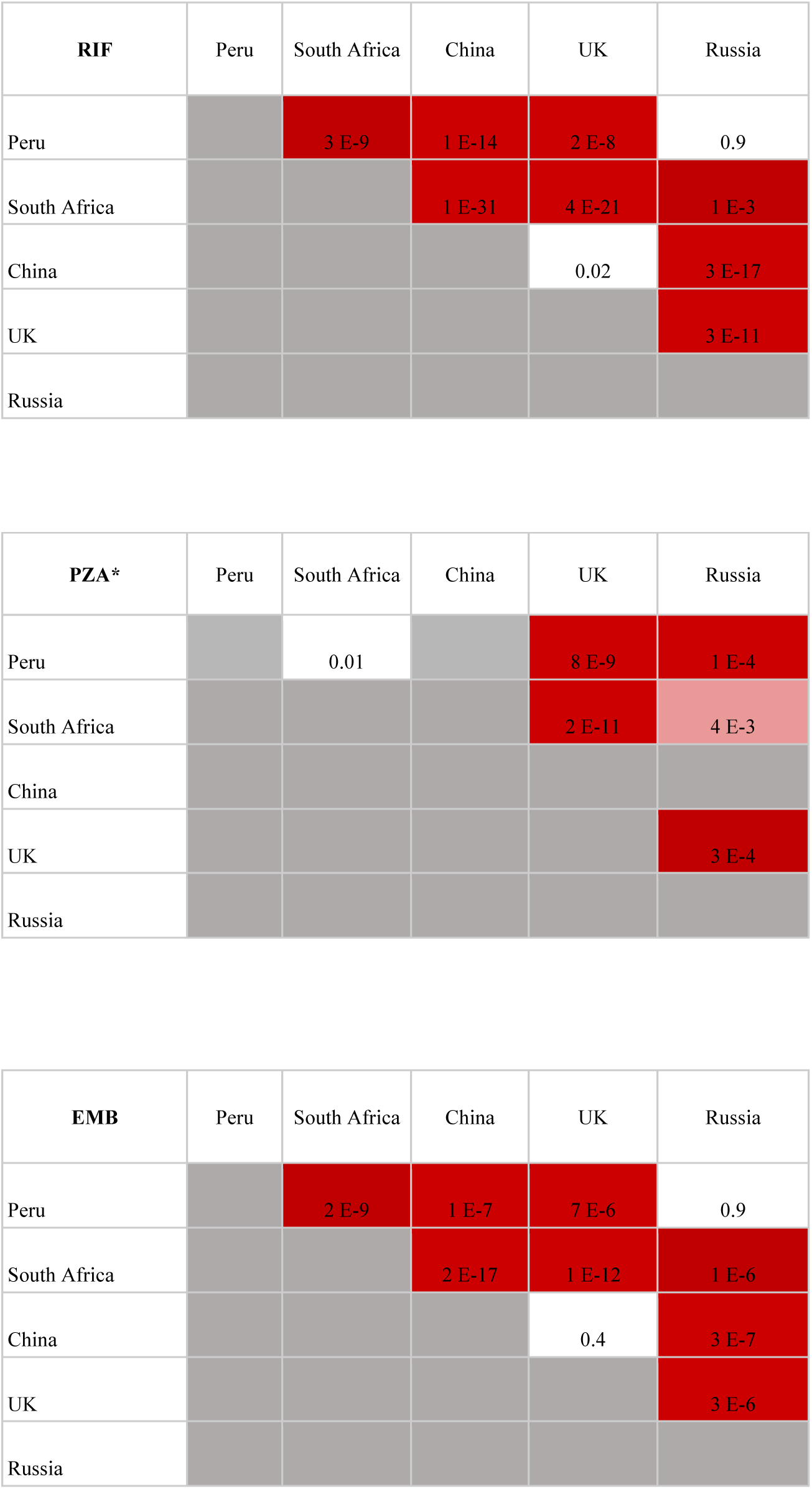
Pairwise Country MRSCA date comparison. Dark red indicates P-value <0.001 (Bonferroni threshold); pink indicates P-value <0.01; white indicates P-value ≥ 0.01. *No PZA resistance phenotypes was available on Chinese isolates.

| INH | Peru | South Africa | China | UK | Russia |
| --- | --- | --- | --- | --- | --- |
| Peru |  | 6 E-9 | 0.1 | 0.2 | 9 E-5 |
| South Africa |  |  | 7 E-29 | 2 E-32 | 3 E-7 |
| China |  |  |  | 0.2 | 4 E-17 |
| UK |  |  |  |  | 5 E-15 |
| Russia |  |  |  |  |  |

| FLQ | Peru | South Africa | China | UK | Russia |
| --- | --- | --- | --- | --- | --- |
| Peru |  | 0.3 | 6 E-10 | 1 E-6 | 1 E-4 |
| South Africa |  |  | 2 E-19 | 1 E-15 | 2 E-7 |
| China |  |  |  | 0.04 | 0.6 |
| UK |  |  |  |  | 0.7 |
| Russia |  |  |  |  |  |

| SLIs | Peru | South Africa | China | UK | Russia |
| --- | --- | --- | --- | --- | --- |
| Peru |  | 0.4 | 1 E-8 | 2 E-5 | 0.2 |
| South Africa |  |  | 5 E-12 | 9 E-8 | 0.5 |
| China |  |  |  | 0.8 | 2 E-4 |
| UK |  |  |  |  | 1 E-3 |
| Russia |  |  |  |  |  |

| **RIF** | Peru | South Africa | China | UK | Russia |
| --- | --- | --- | --- | --- | --- |
| Peru |  | 3 E-9 | 1 E-14 | 2 E-8 | 0.9 |
| South Africa |  |  | 1 E-31 | 4 E-21 | 1 E-3 |
| China |  |  |  | 0.02 | 3 E-17 |
| UK |  |  |  |  | 3 E-11 |
| Russia |  |  |  |  |  |

| **PZA\*** | Peru | South Africa | China | UK | Russia |
| --- | --- | --- | --- | --- | --- |
| Peru |  | 0.01 |  | 8 E-9 | 1 E-4 |
| South Africa |  |  |  | 2 E-11 | 4 E-3 |
| China |  |  |  |  |  |
| UK |  |  |  |  | 3 E-4 |
| Russia |  |  |  |  |  |

| **EMB** | Peru | South Africa | China | UK | Russia |
| --- | --- | --- | --- | --- | --- |
| Peru |  | 2 E-9 | 1 E-7 | 7 E-6 | 0.9 |
| South Africa |  |  | 2 E-17 | 1 E-12 | 1 E-6 |
| China |  |  |  | 0.4 | 3 E-7 |
| UK |  |  |  |  | 3 E-6 |
| Russia |  |  |  |  |  |

**Supplementary Table 4.** MDR Frequency Comparison: Comparison of MDR Frequency between our estimate based on WGS data (WGS MDR Frequency) and estimate based on WHO data (WHO MDR Frequency). Number of isolates with resistance data per country is provided in the “Total Number of Isolates” column.

|  | WGS MDR Frequency | WHO MDR Frequency | Total Number of Isolates |
| --- | --- | --- | --- |
| **Belarus** | 0.03±0.07 | 0.71(0.67-0.75) | 30 |
| **Canada** | 0 | 0.01(0.01-0.02) | 165 |
| **China** | 0.72±0.07 | 0.08(0.07-0.09) | 163 |
| **Germany** | 0.02±0.01 | 0.03(0.02-0.05) | 850 |
| **India** | 0.33±0.15 | 0.05(0.04-0.06) | 39 |
| **Iran** | 0.29±0.19 | 0.02(0.01-0.02) | 21 |
| **Malawi** | 0.01±0.01 | 0.01(0.01-0.02) | 1403 |
| **Mali** | 0.26±0.14 | 0.03(0.02-0.04) | 35 |
| **Moldova** | 0.05±0.07 | 0.34(0.31-0.36) | 40 |
| **Netherlands** | 0 | 0.02(<0.01-0.01) | 93 |
| **Not Provided** | 0.63±0.03 |  | 1030 |
| **Peru** | 0.70±0.03 | 0.09 (0.09-0.1) | 941 |
| **Romania** | 1 | 0.05(0.05-0.06) | 33 |
| **Russia** | 0.49±0.03 | 0.43(0.43-0.44) | 854 |
| **Sierra Leone** | 0.10±0.07 | 0.03(0.02-0.04) | 79 |
| **South Africa** | 0.20±0.03 | 0.04(0.04-0.05) | 882 |
| **South Korea** | 0.61±0.15 | 0.04(0.03-0.04) | 41 |
| **Swaziland** | 0.10±0.19 | 0.1(0.09-0.11) | 10 |
| **Thailand** | 0.94±0.12 | 0.03(0.03-0.04) | 16 |
| **Turkmenistan** | 0.10±0.17 | 0.22(0.19-0.24) | 11 |
| **Uganda** | 0.78±0.11 | 0.02(0.02-0.03) | 58 |
| **United Kingdom** | 0.04±0.01 | 0.02(0.01-0.02) | 1850 |
| **United States of America** | 0.97±0.06 | 0.02(0.01-0.02) | 30 |
| **Uzbekistan** | 0.99±0.01 | 0.33(0.29-0.34) | 265 |

**Supplementary Table 5.** Dates Drugs Introduced.

| **Drug** | **Year Introduced** |
| --- | --- |
| Isoniazid<sup>18</sup> | 1952 |
| Rifampicin<sup>18</sup> | 1965 |
| Pyrazinamide<sup>19</sup> | 1952 |
| Streptomycin<sup>18</sup> | 1944 |
| Ethambutol<sup>18</sup> | 1965 |
| Ethionamide<sup>18</sup> | 1960 |
| Kanamycin/amikacin<sup>18</sup> | 1958 |
| Cycloserine<sup>18</sup> | 1955 |
| Capreomycin<sup>18</sup> | 1967 |
| PAS<sup>18</sup> | 1944 |
| Ofloxacin<sup>20</sup> | 1985 |
| Moxifloxacin<sup>20</sup> | 1999 |
| Ciprofloxacin<sup>20</sup> | 1986 |

**Supplementary Table 6.**
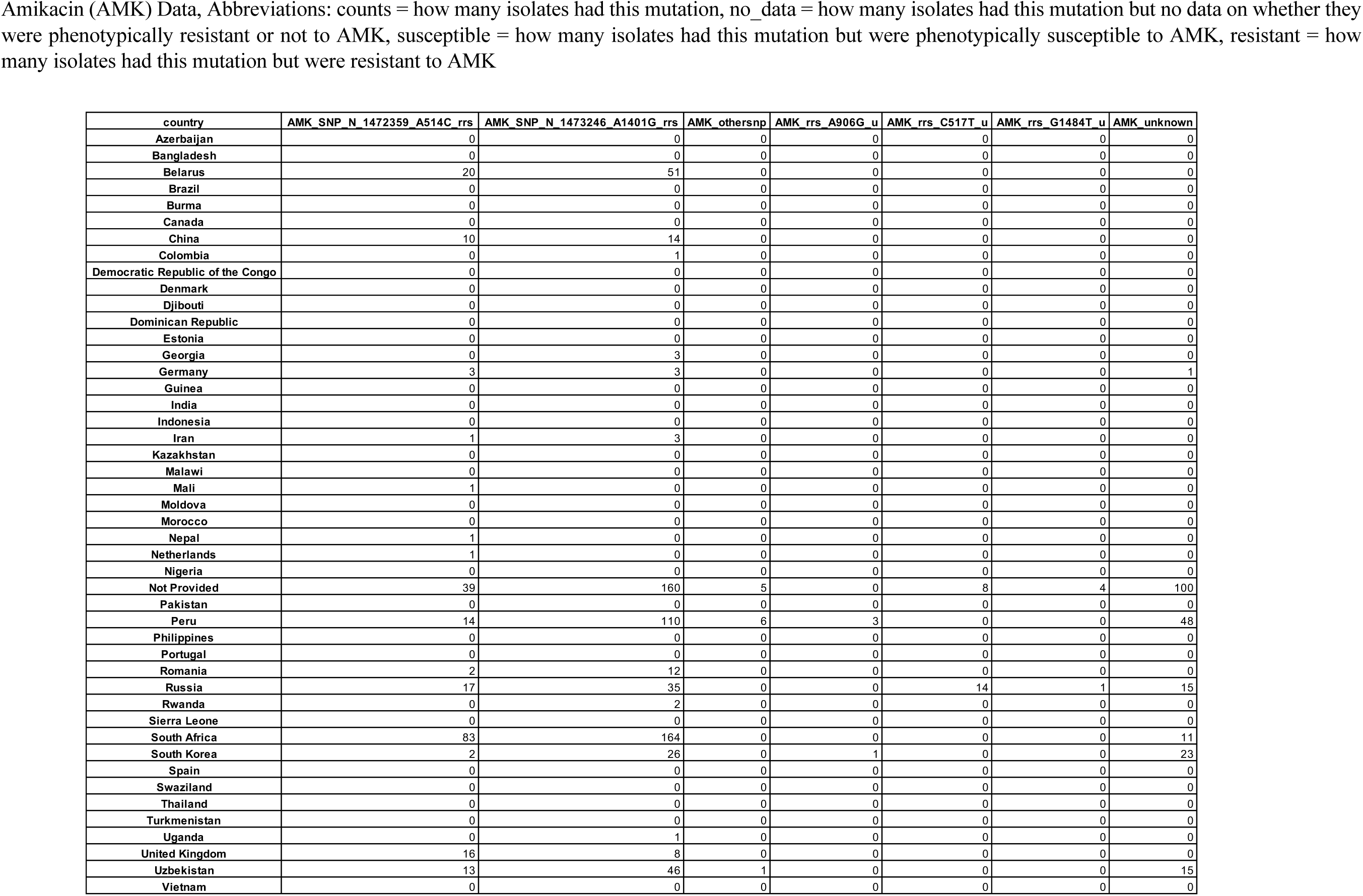
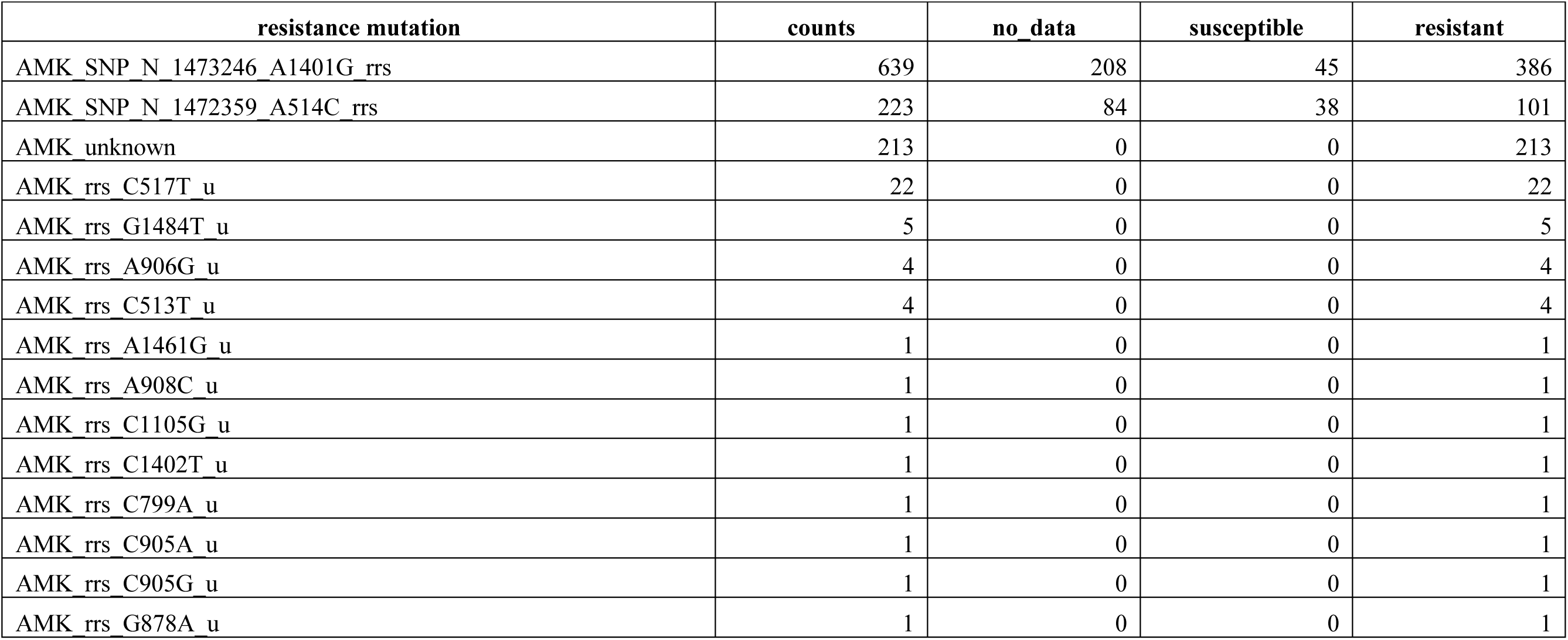
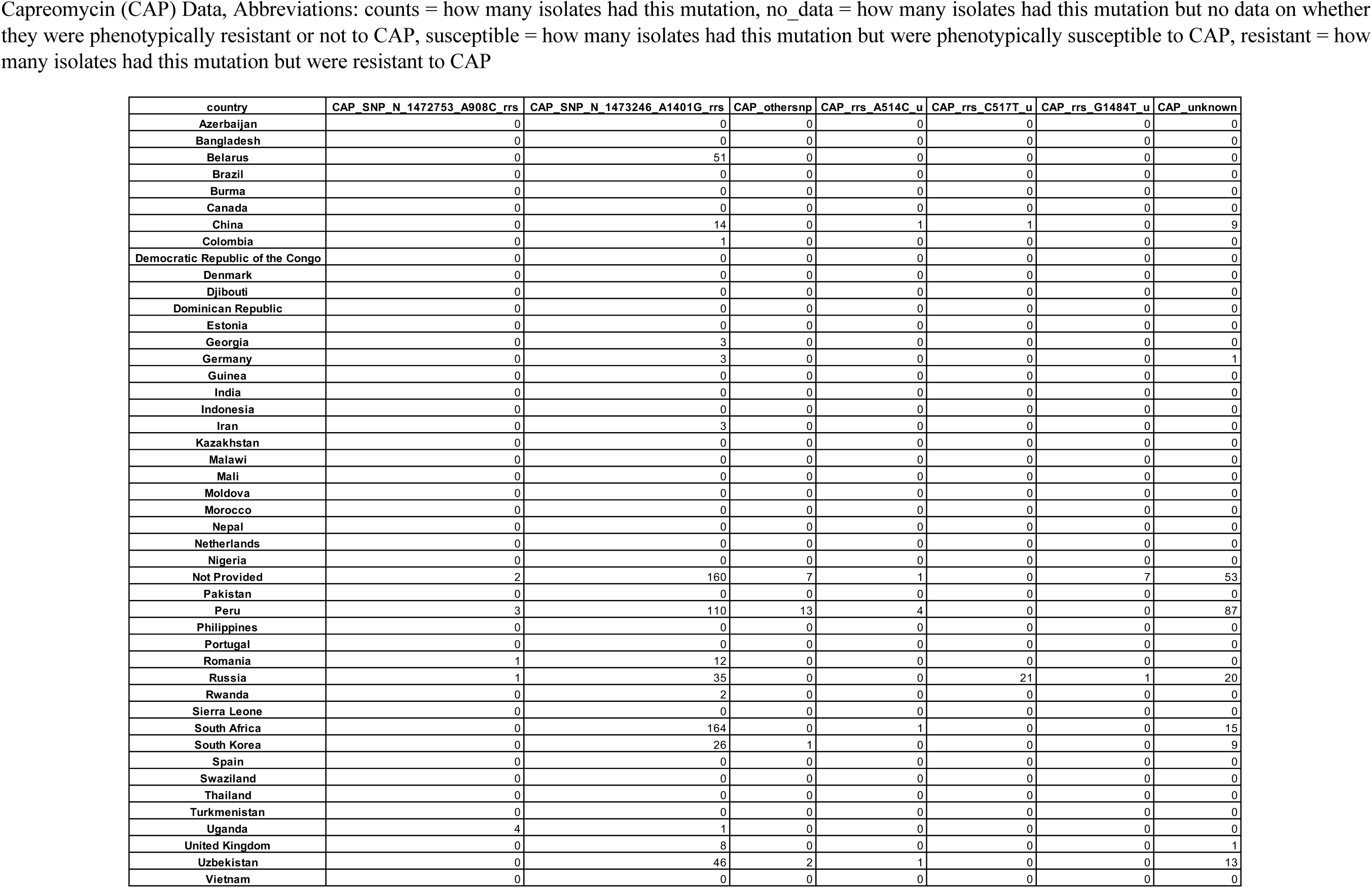
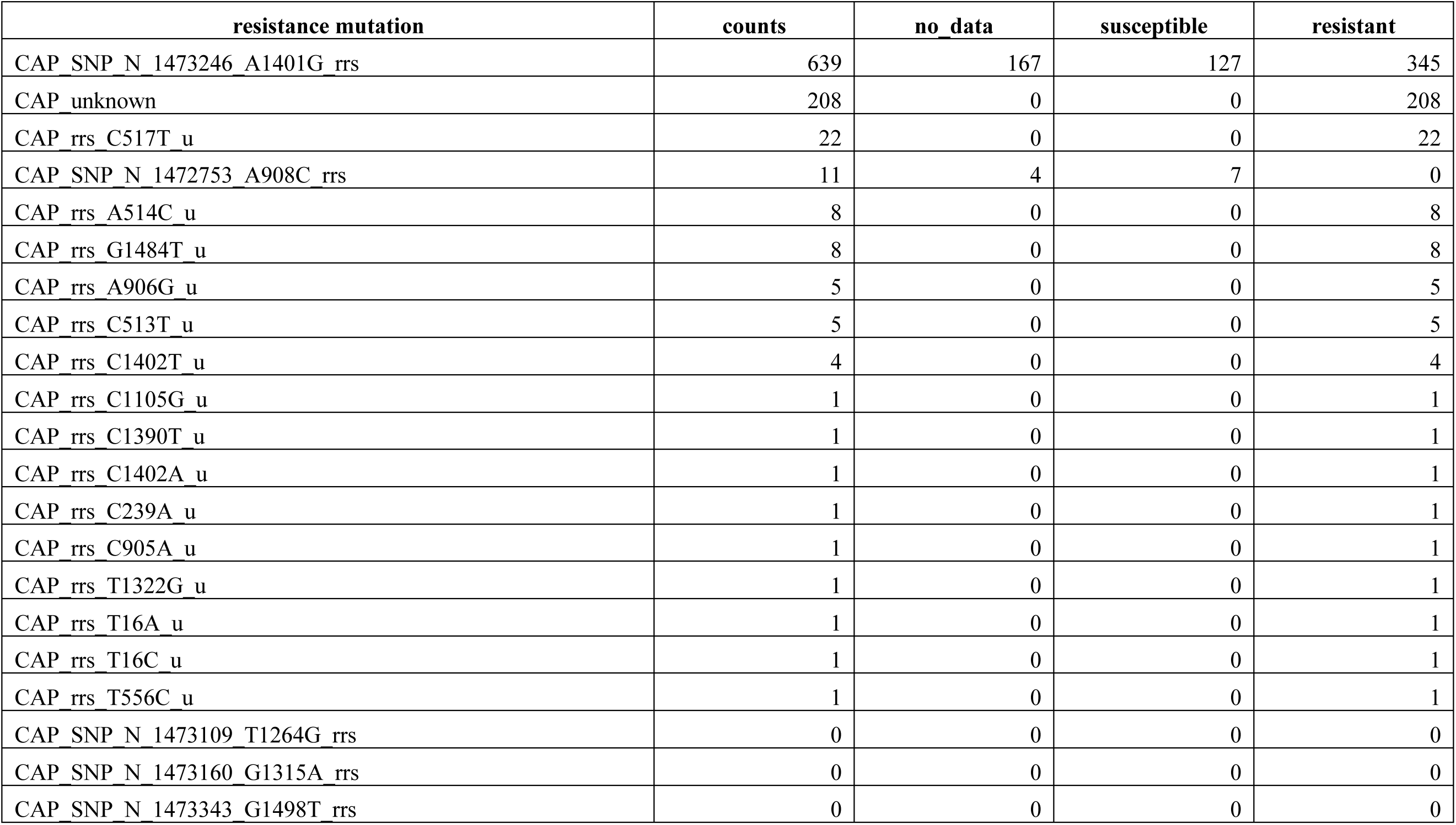
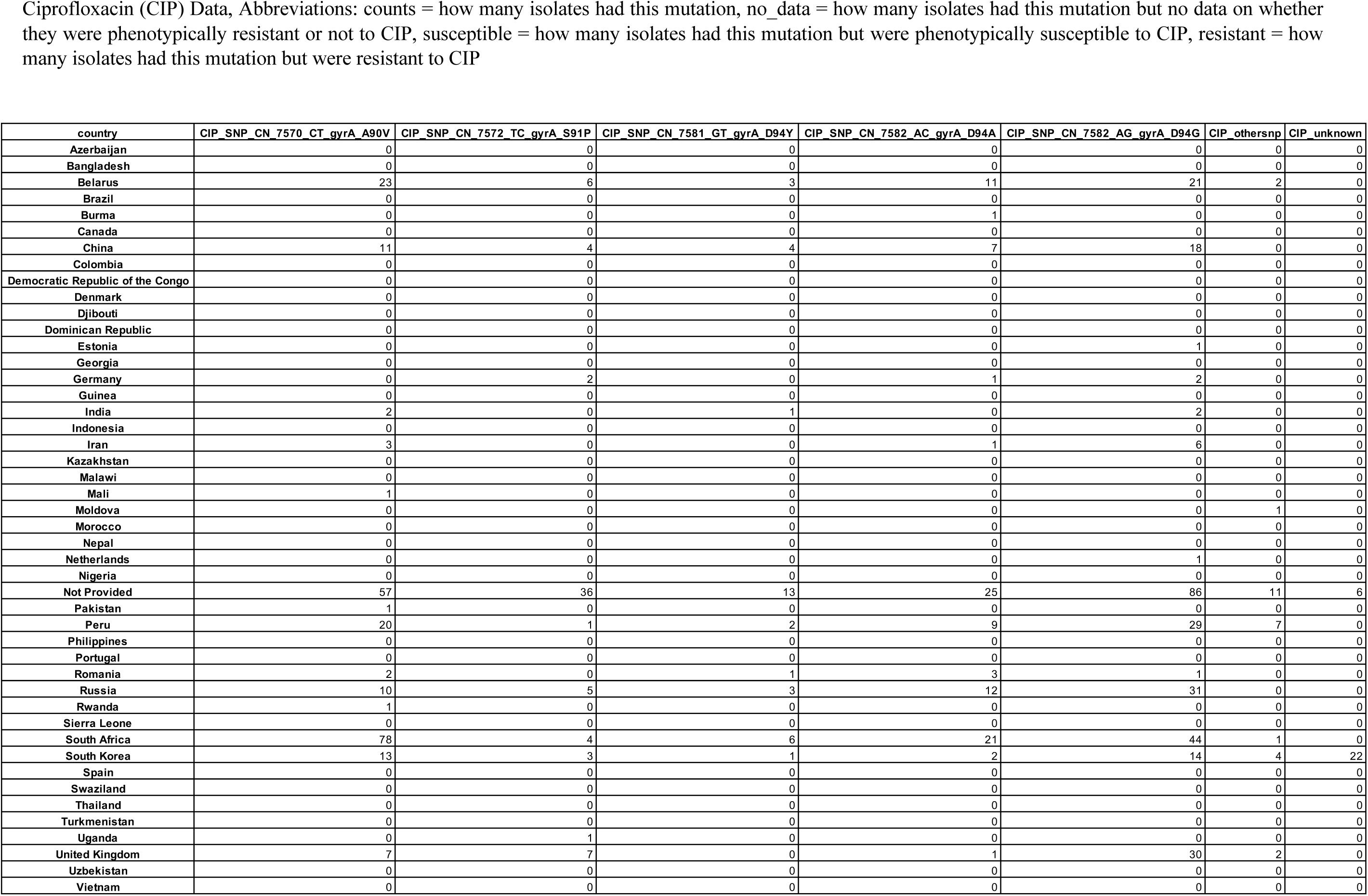
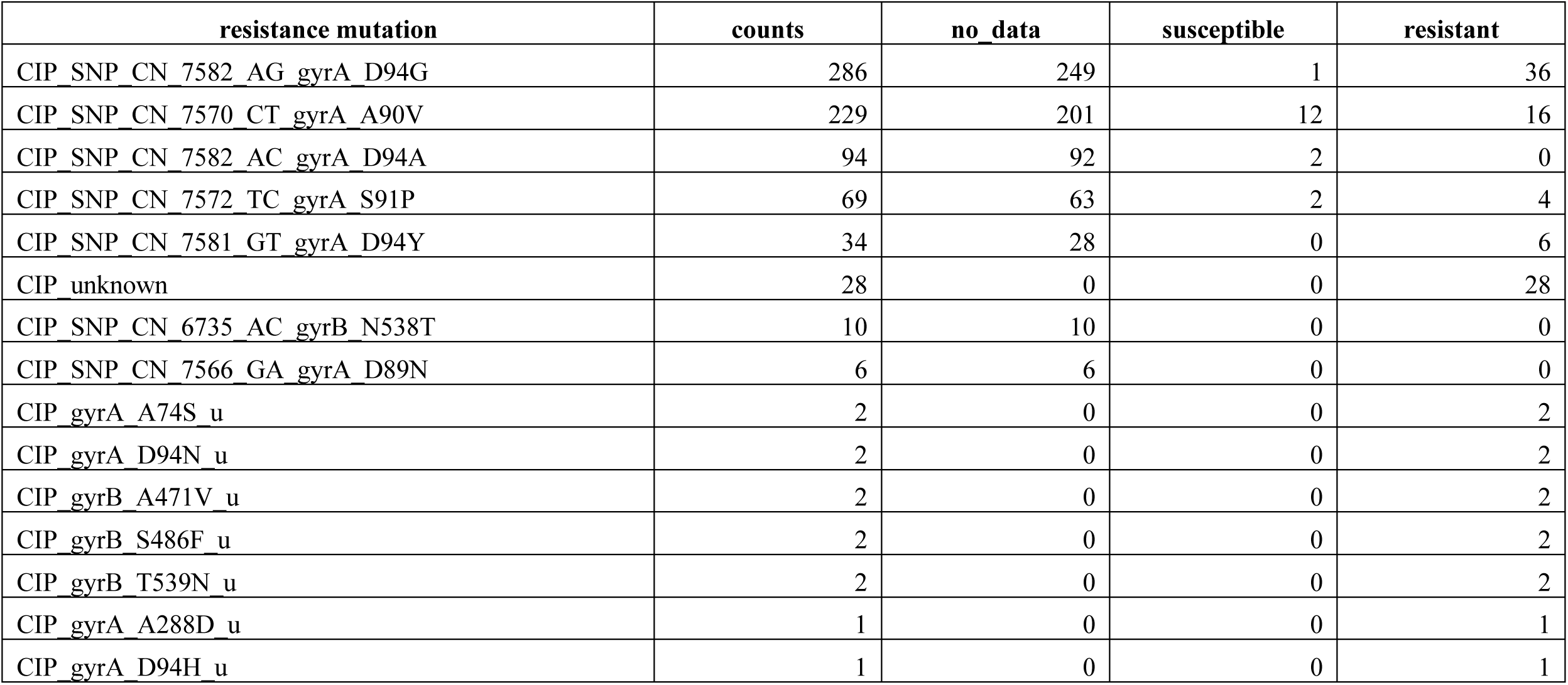
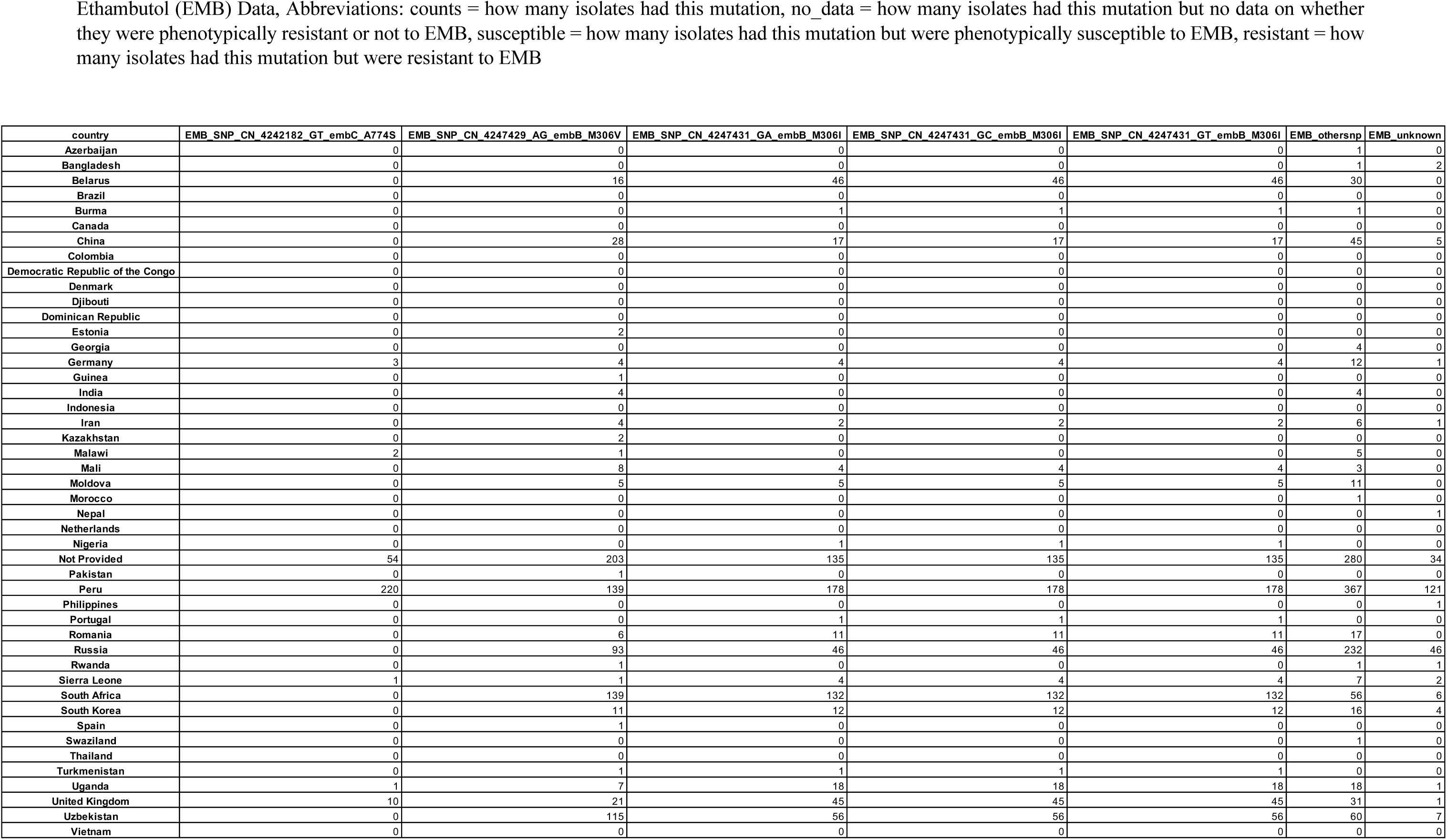
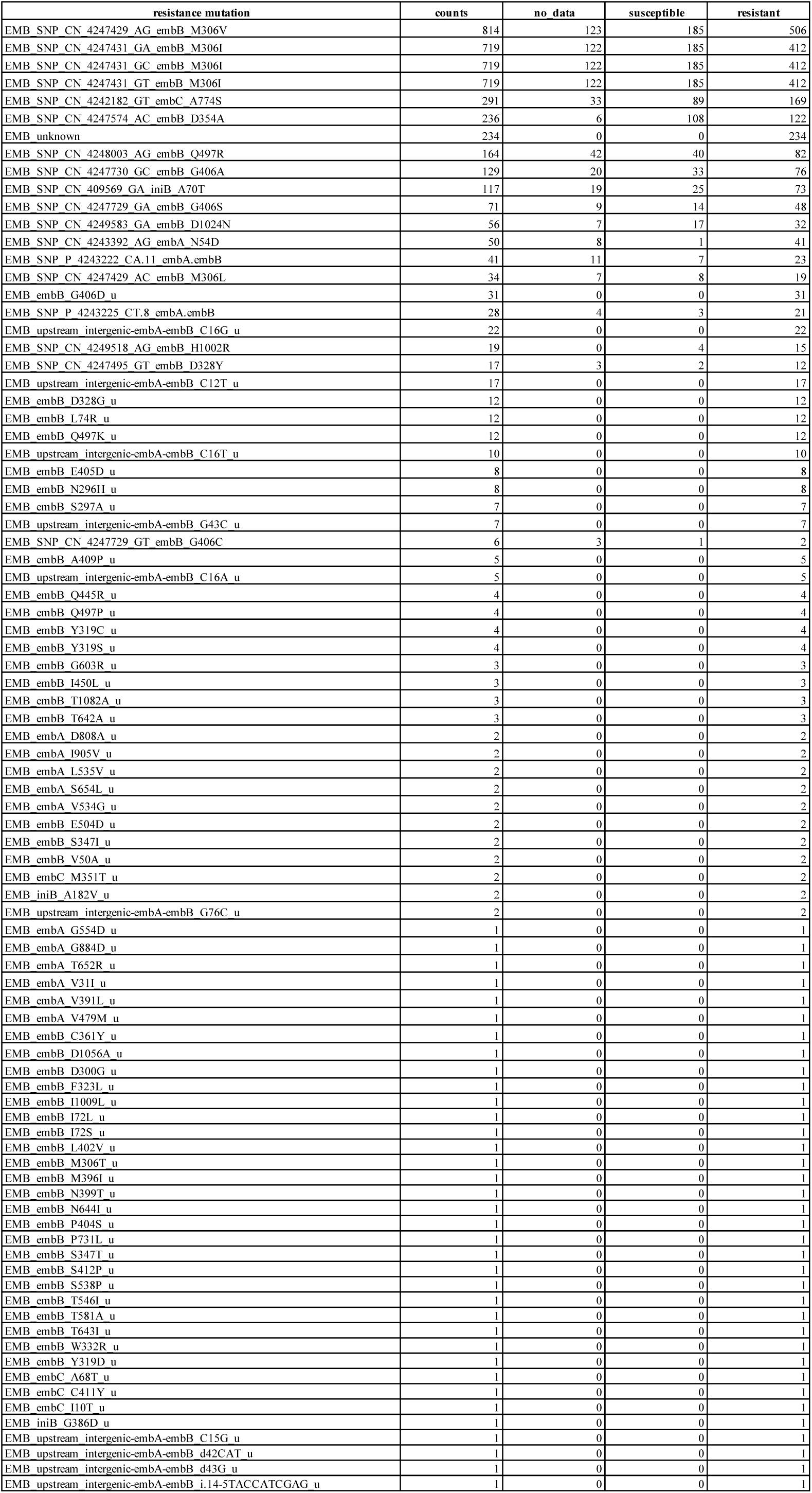
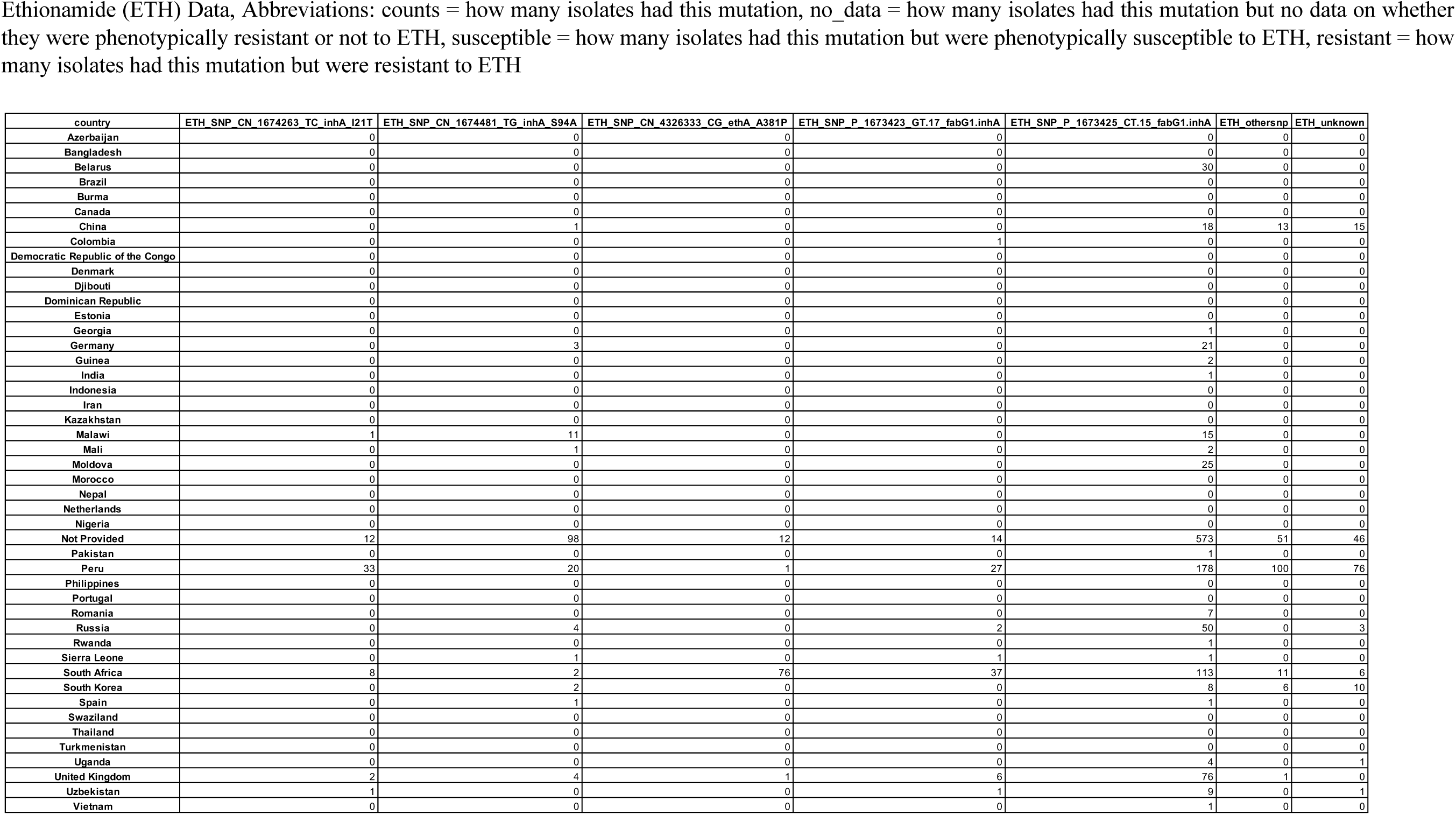
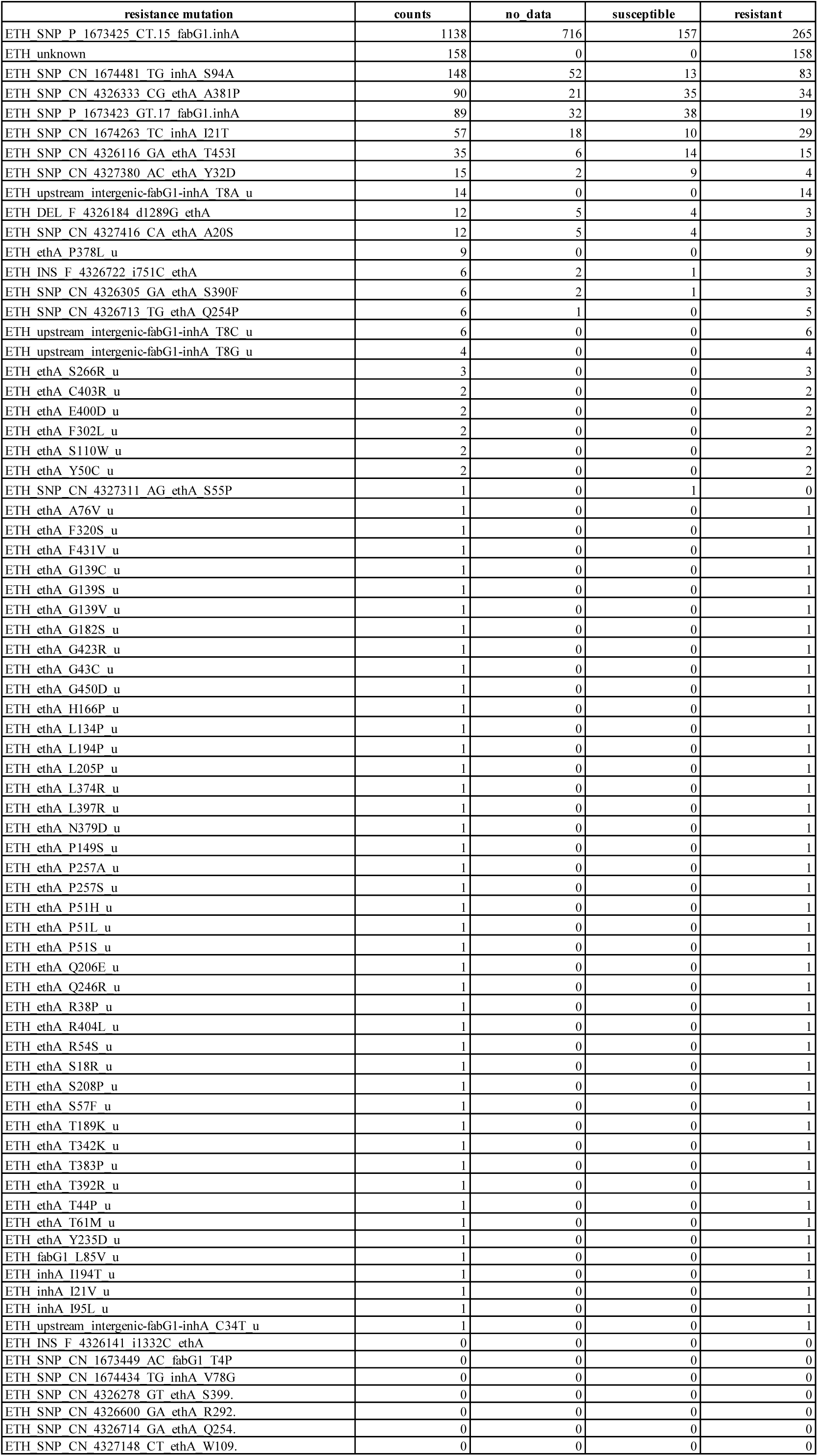
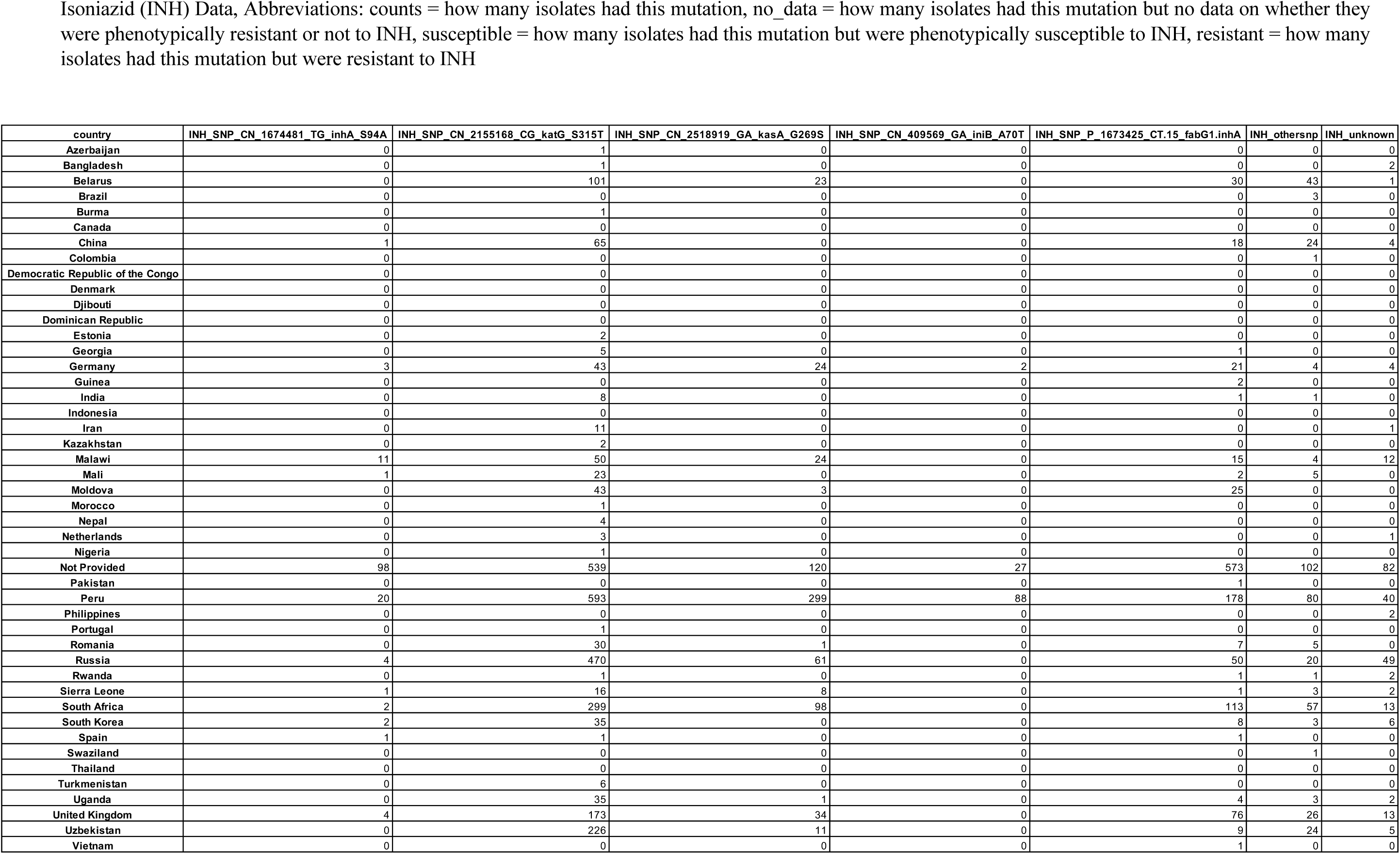
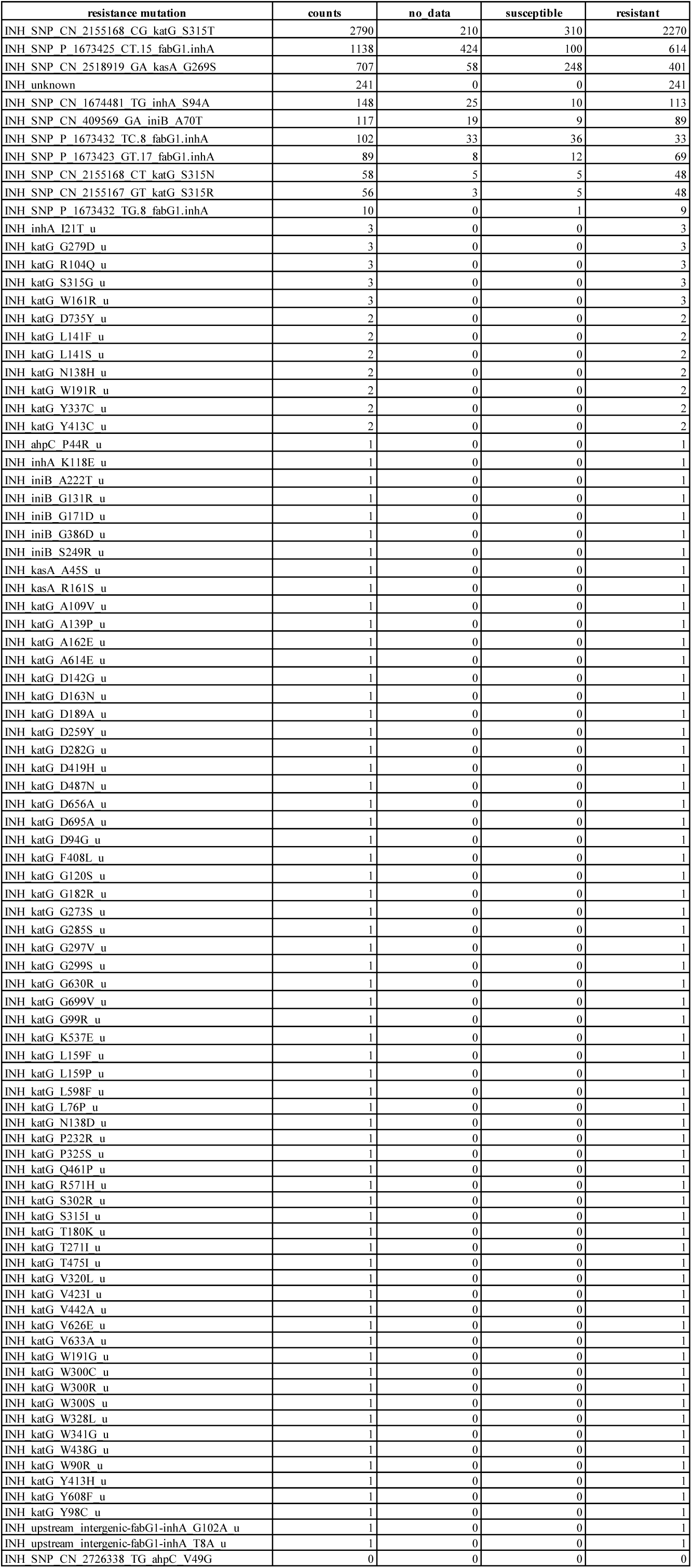
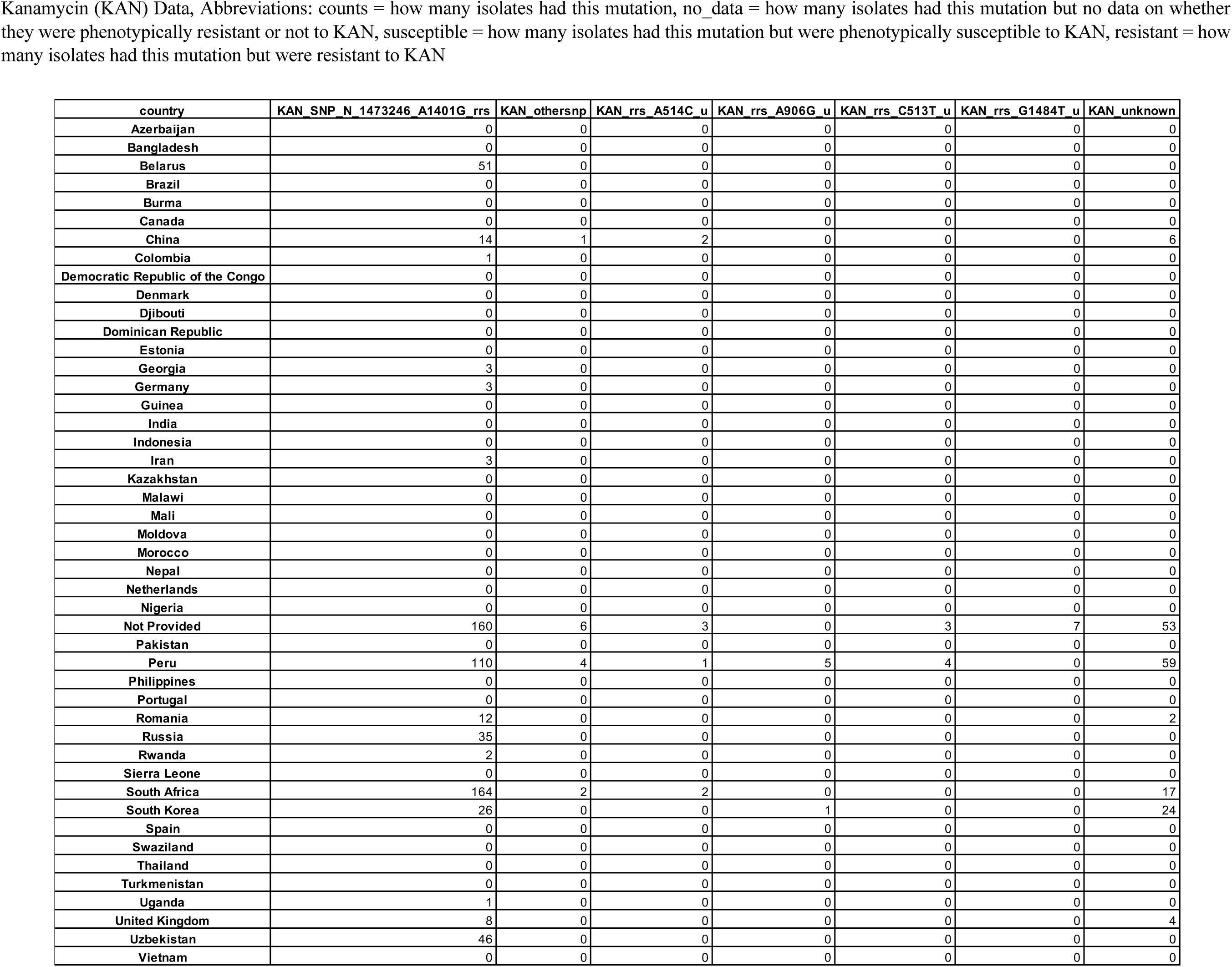
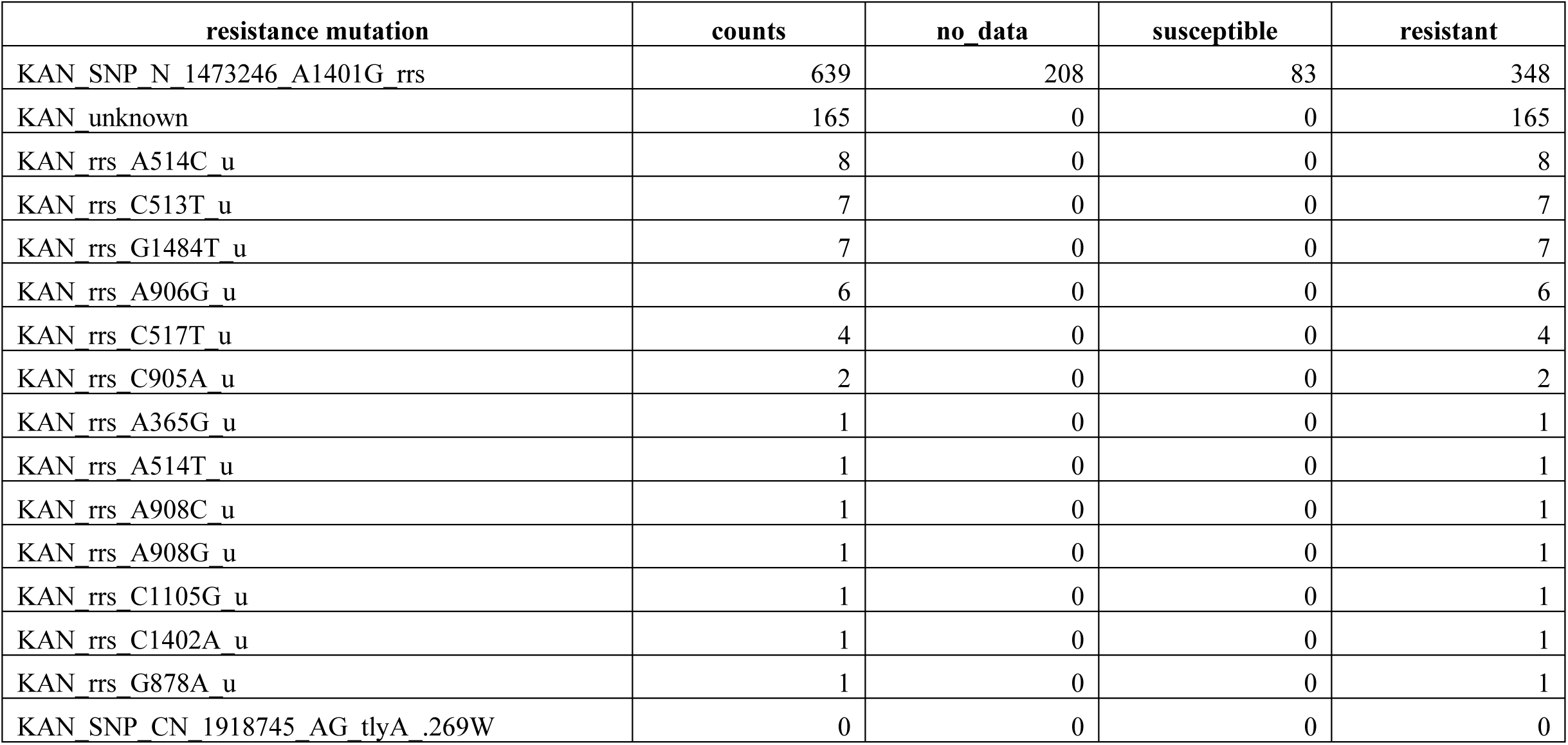
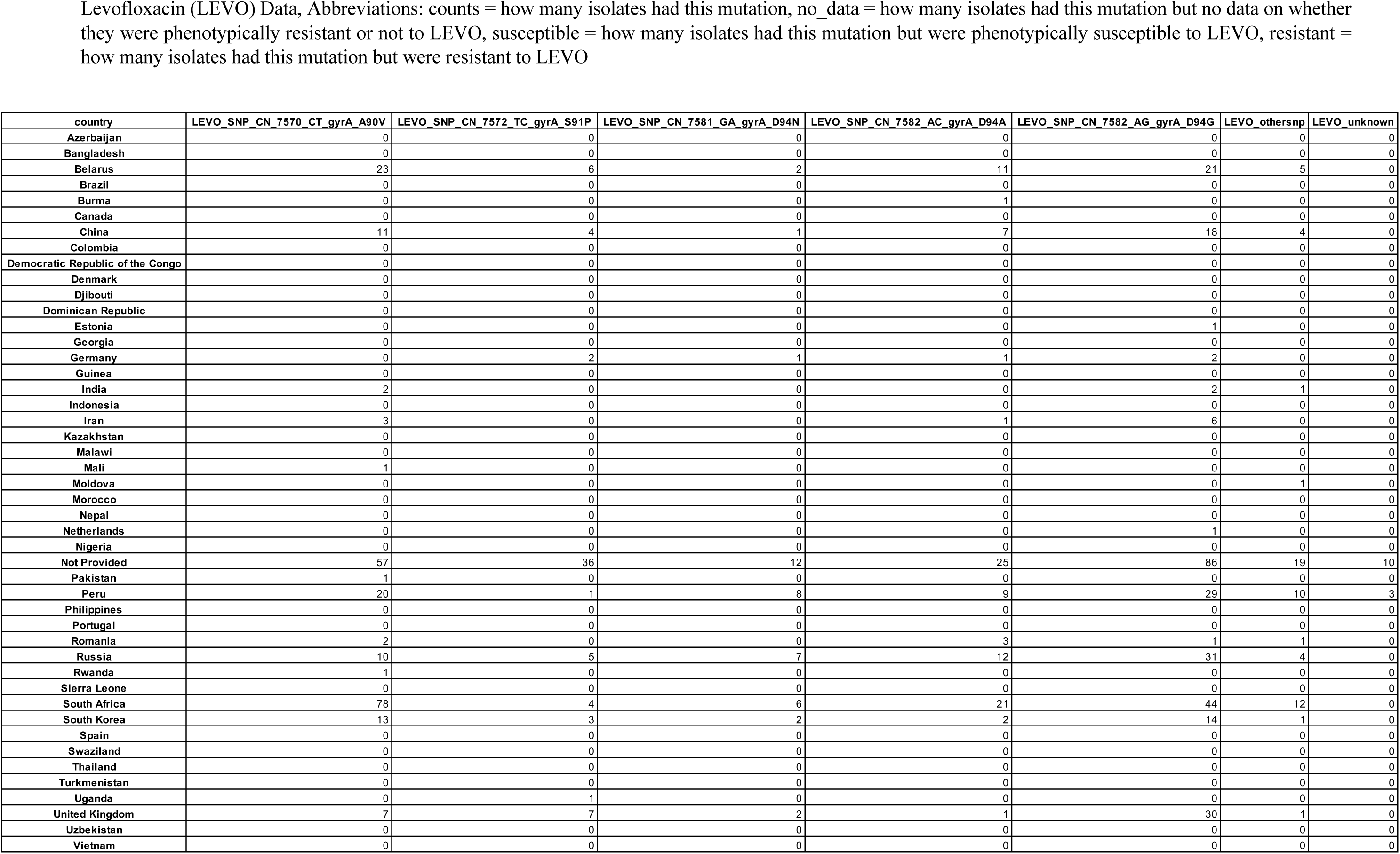
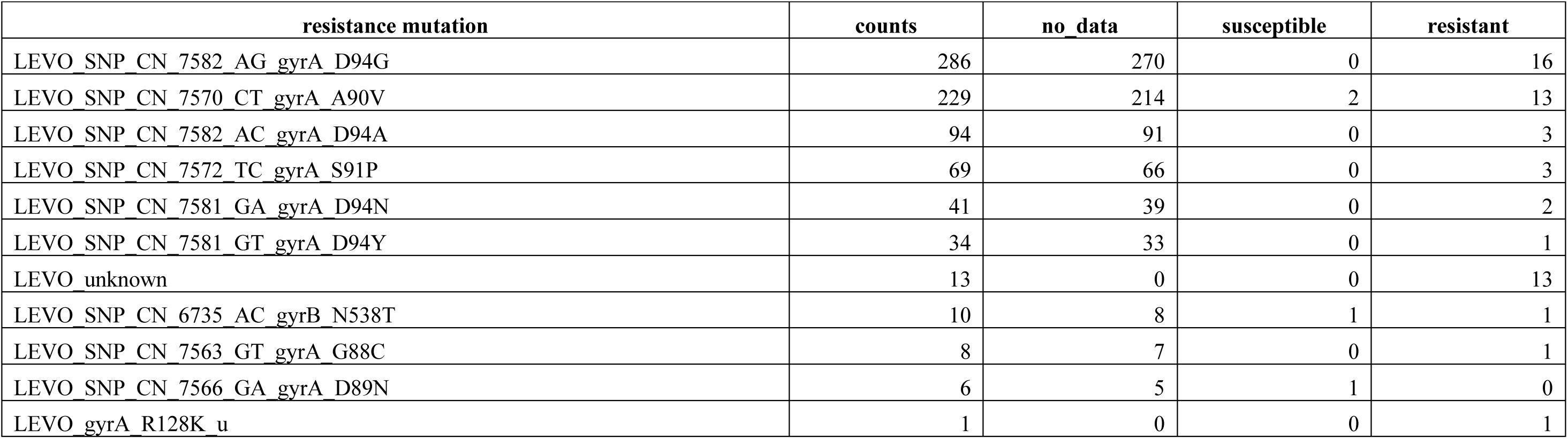
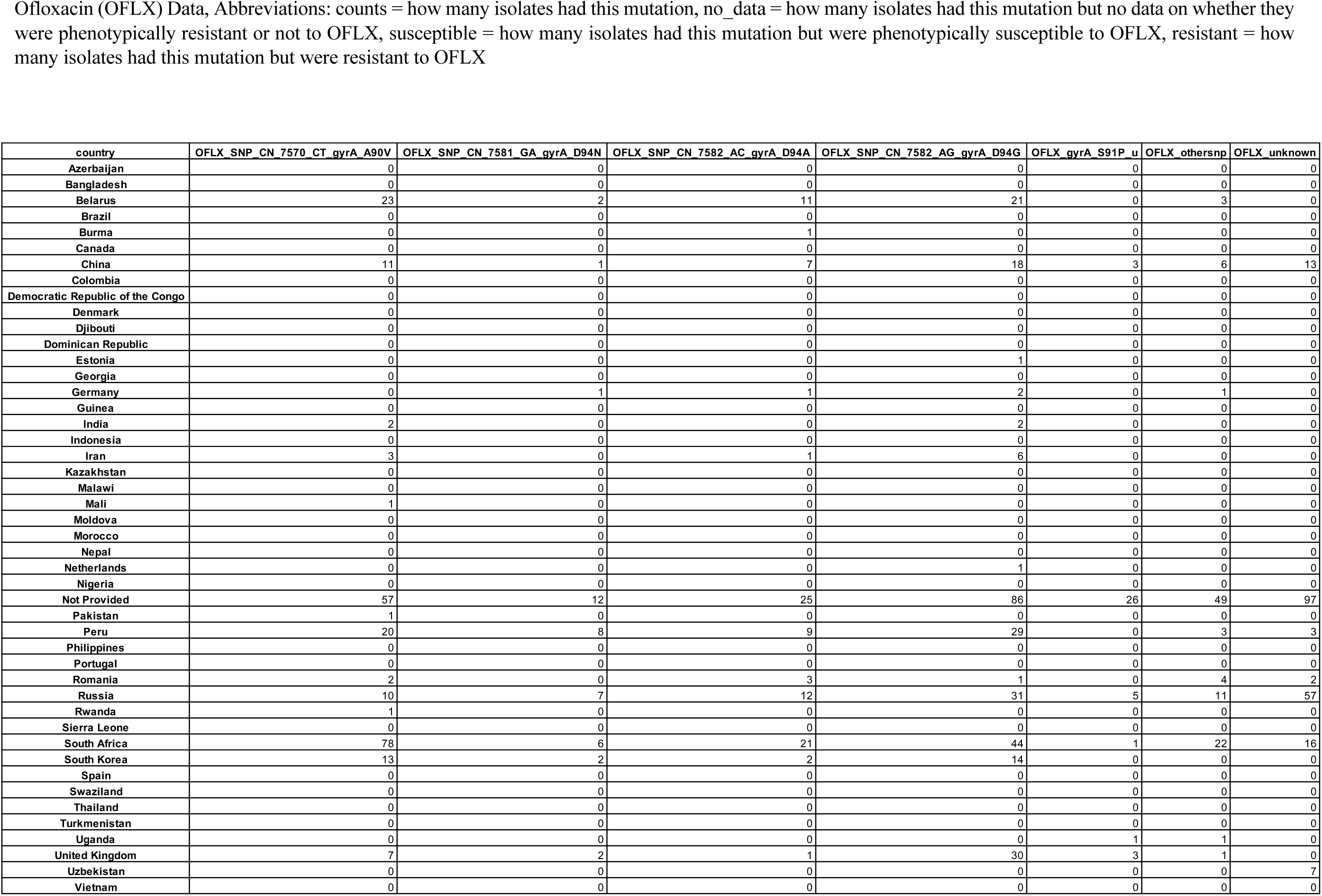
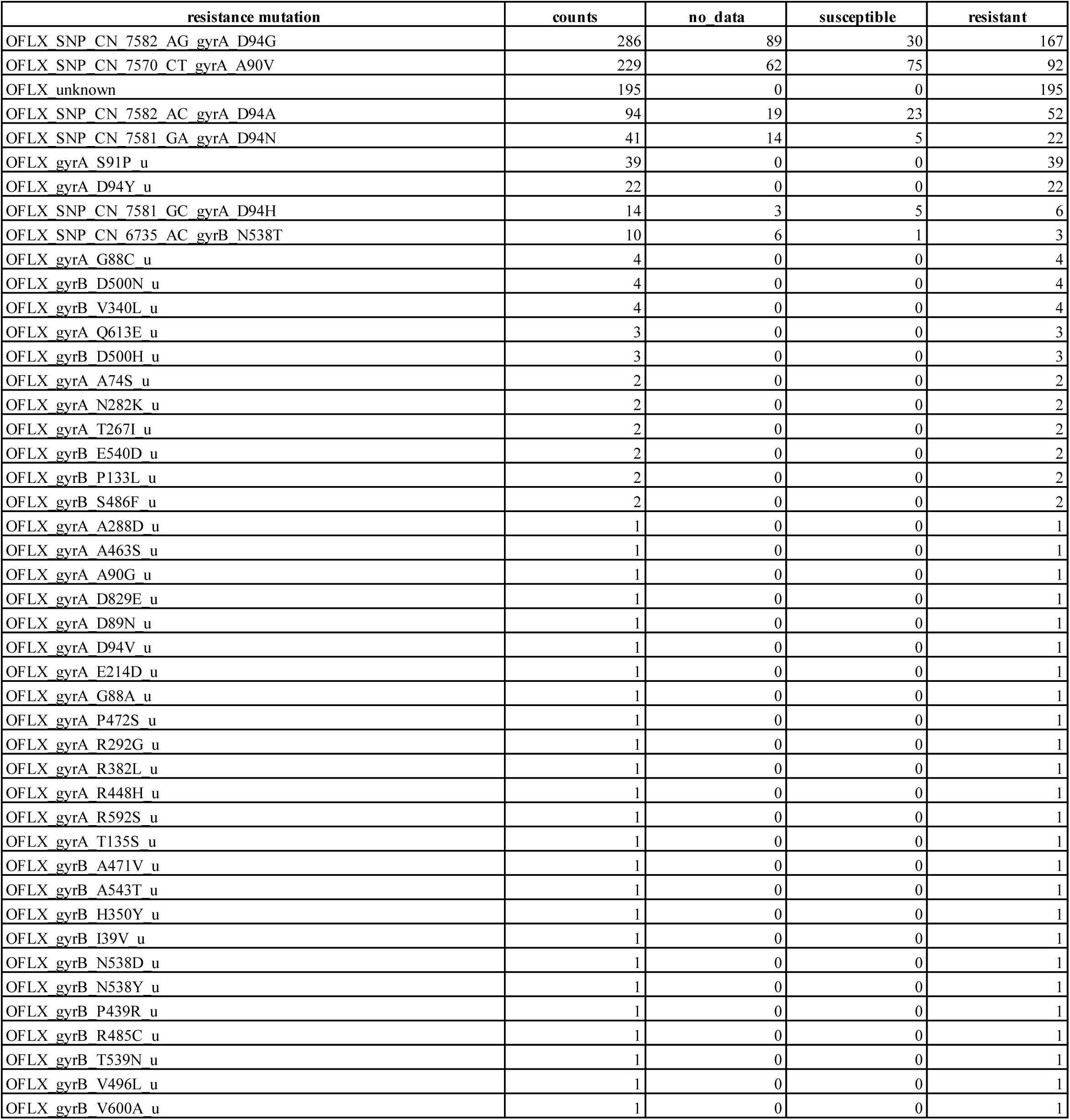
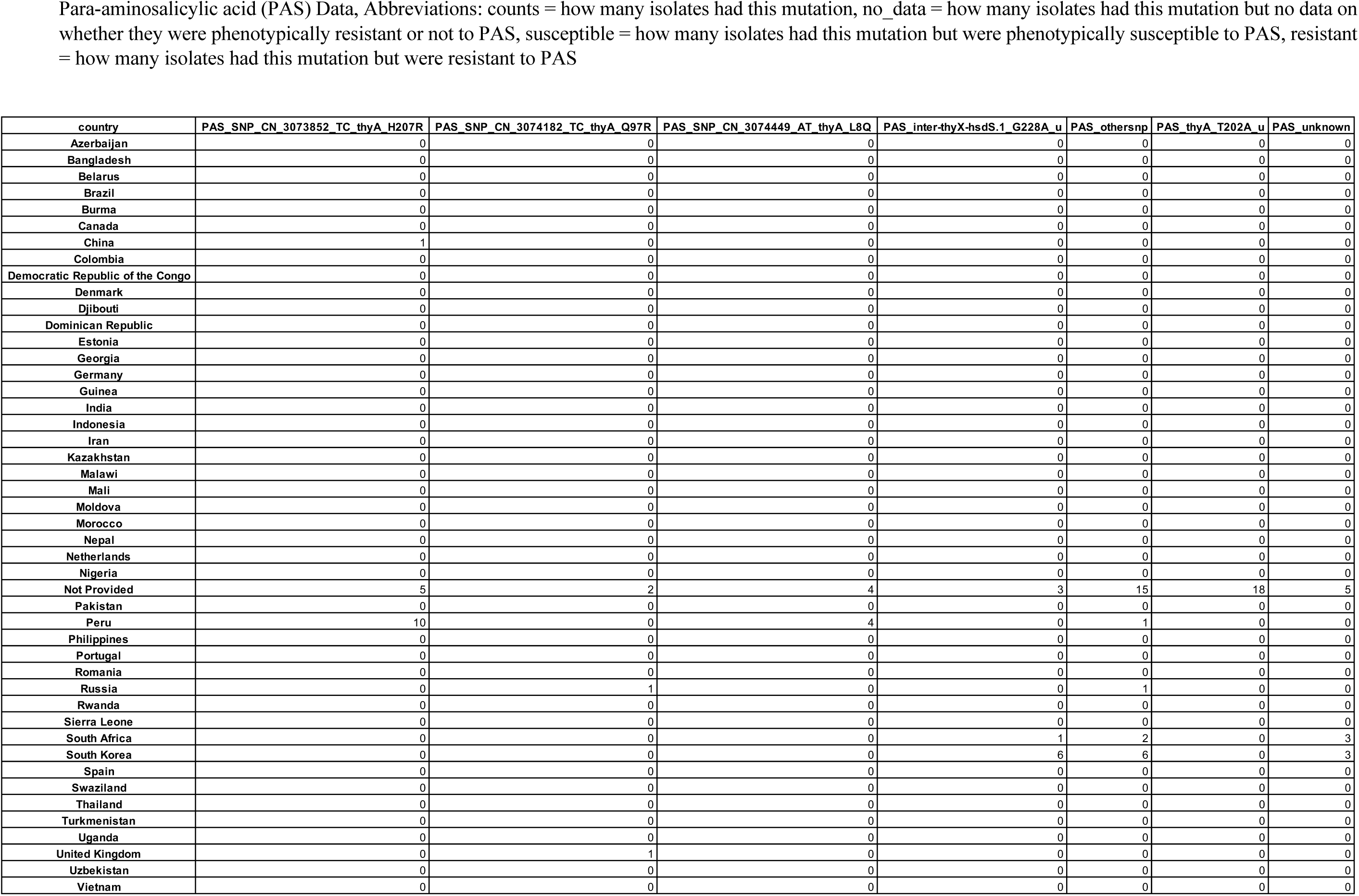
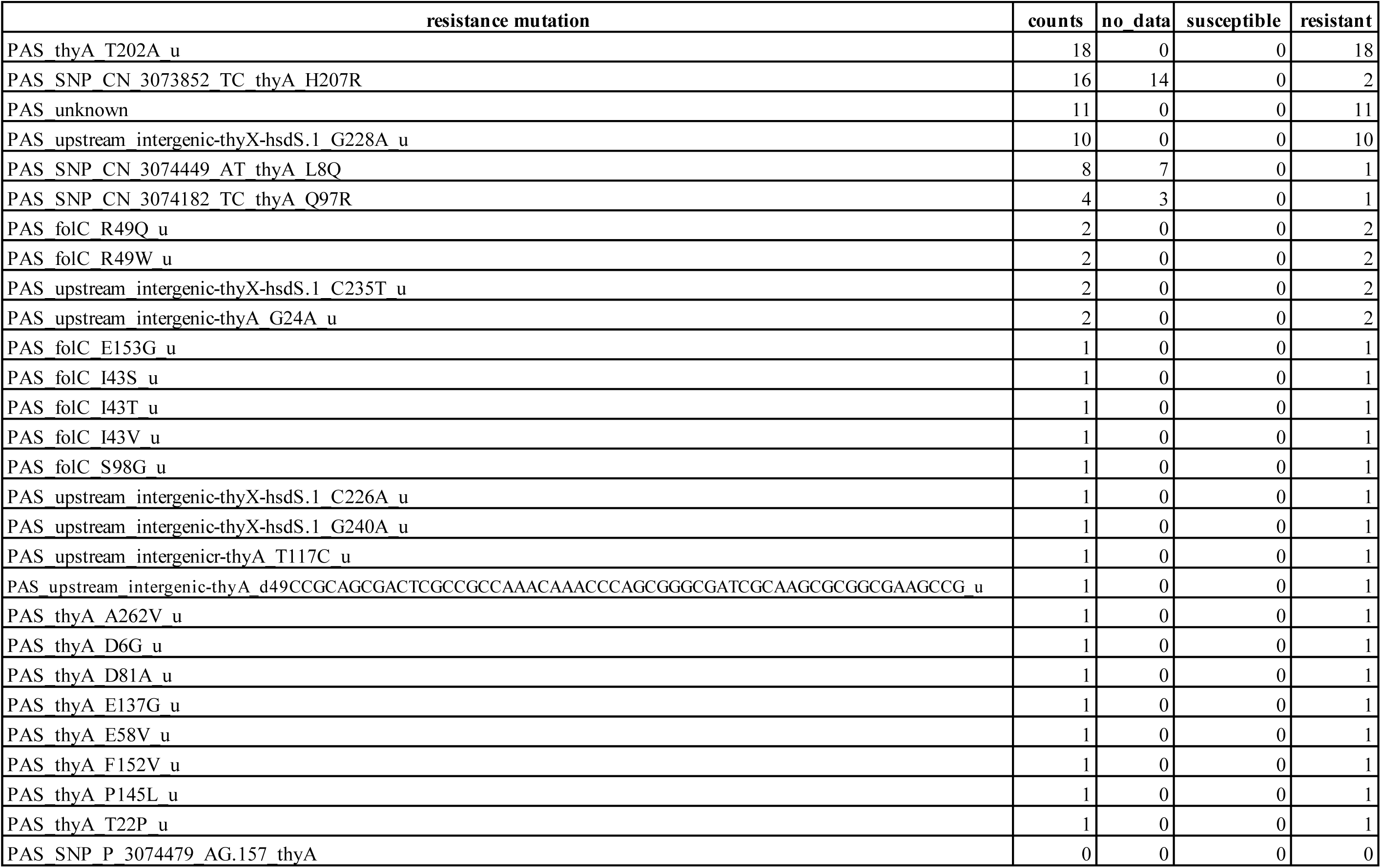
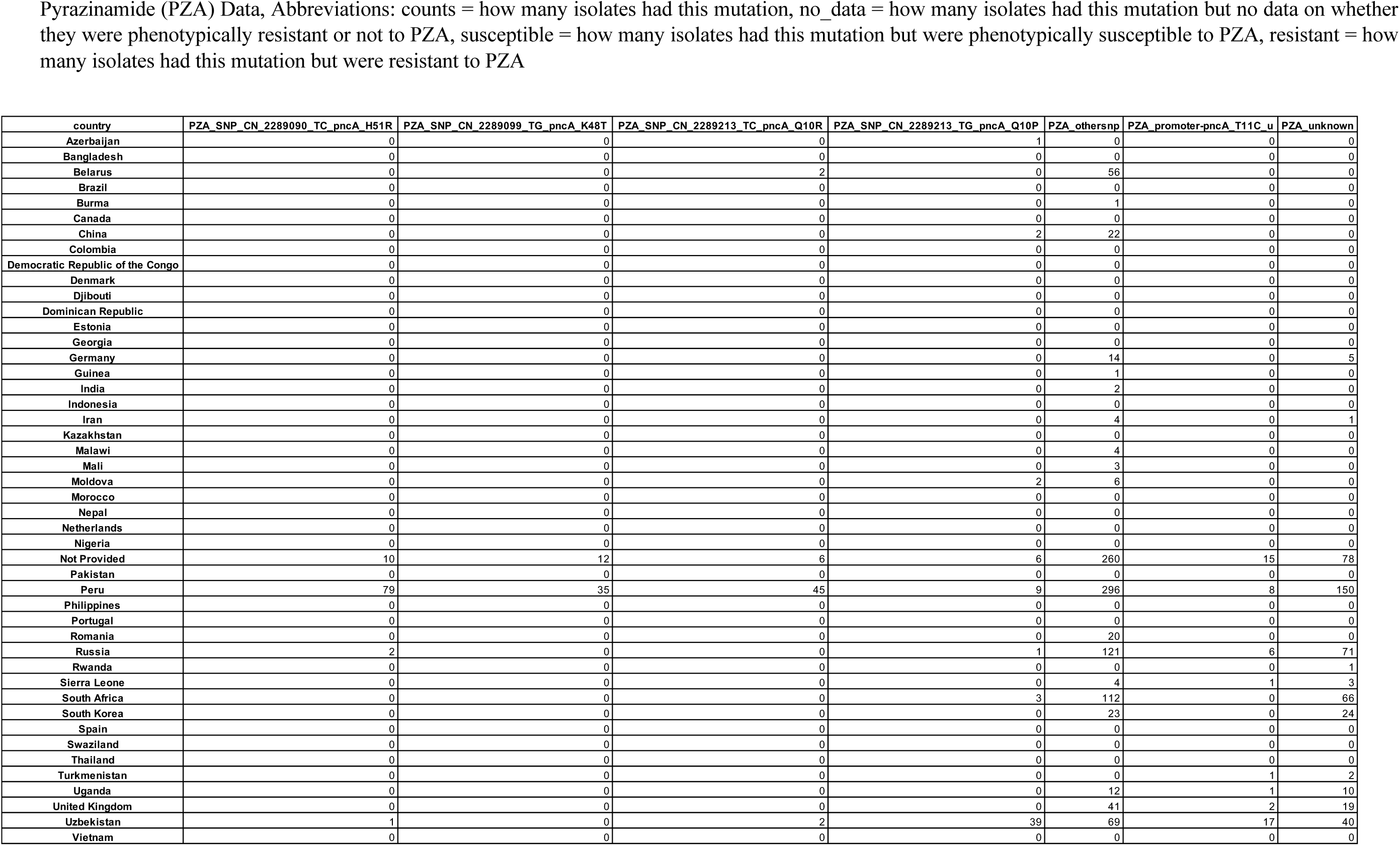
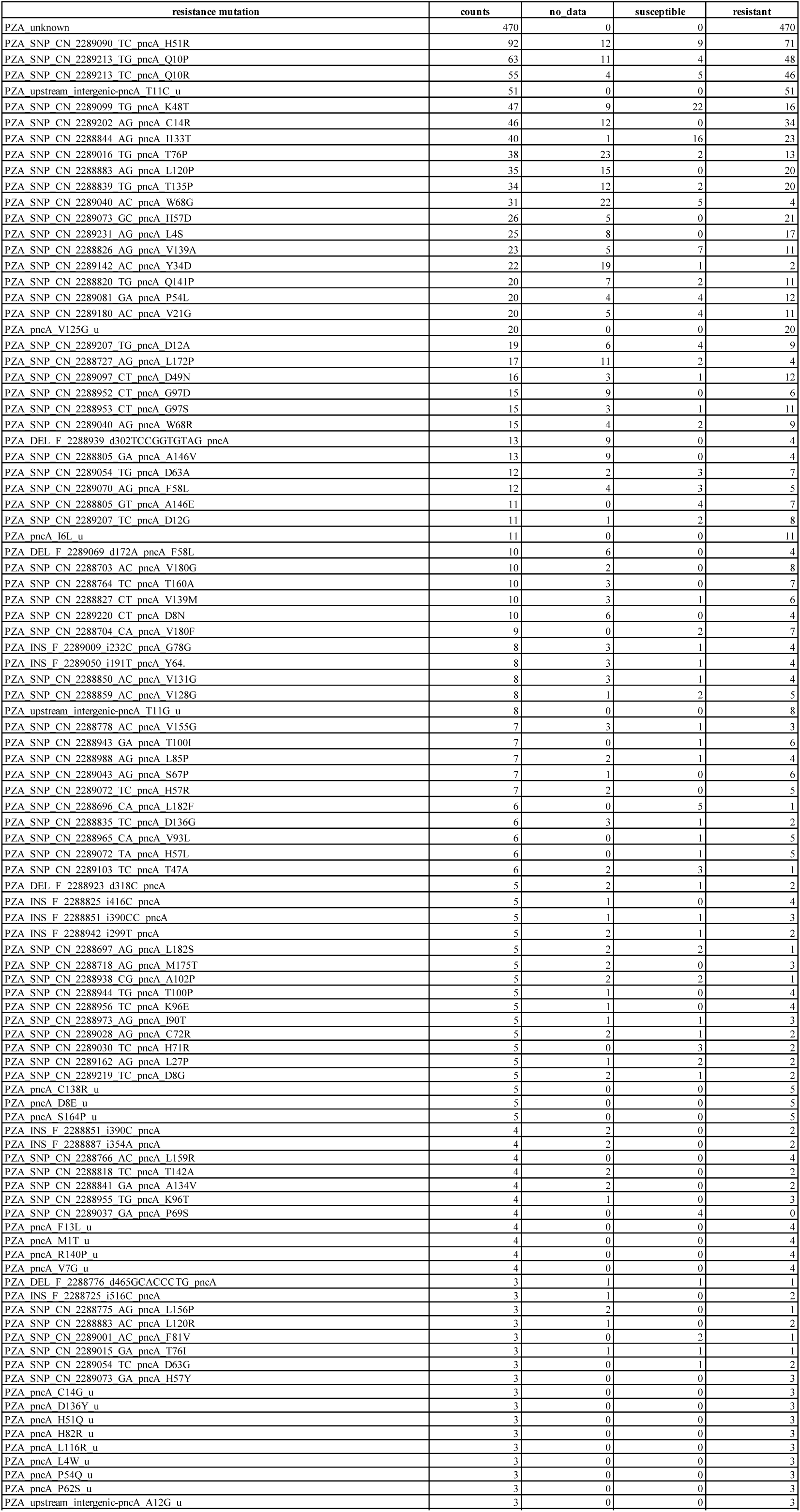
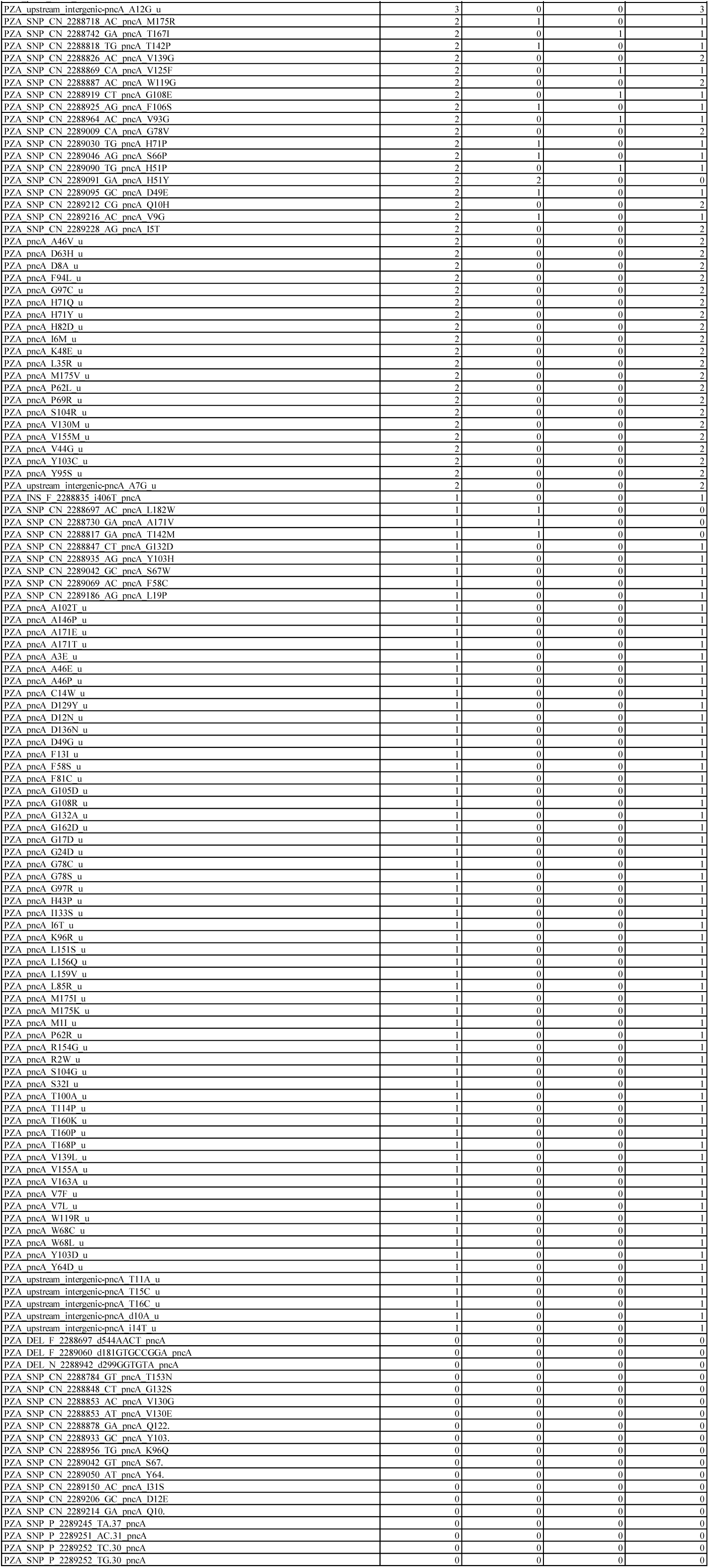
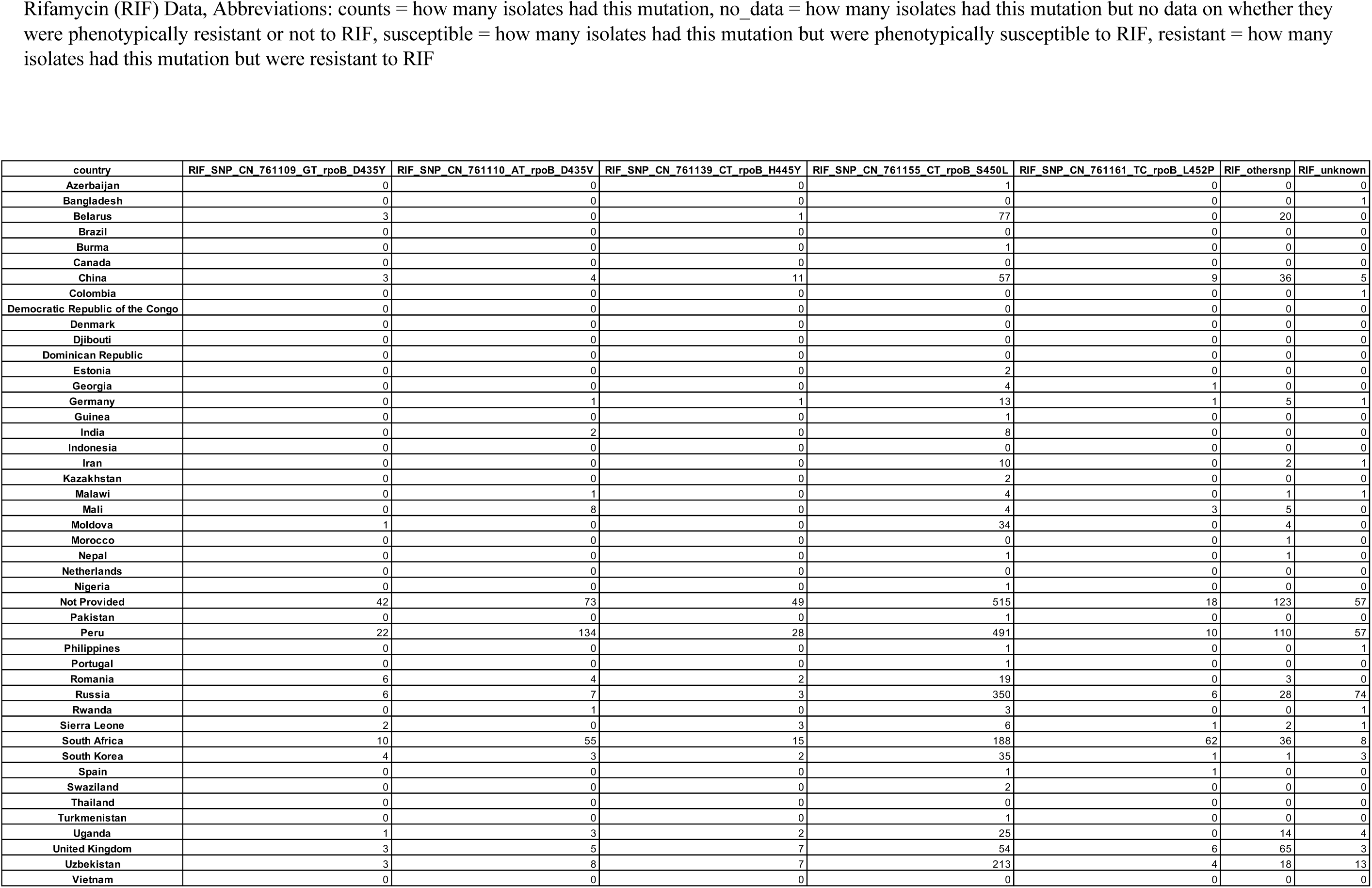
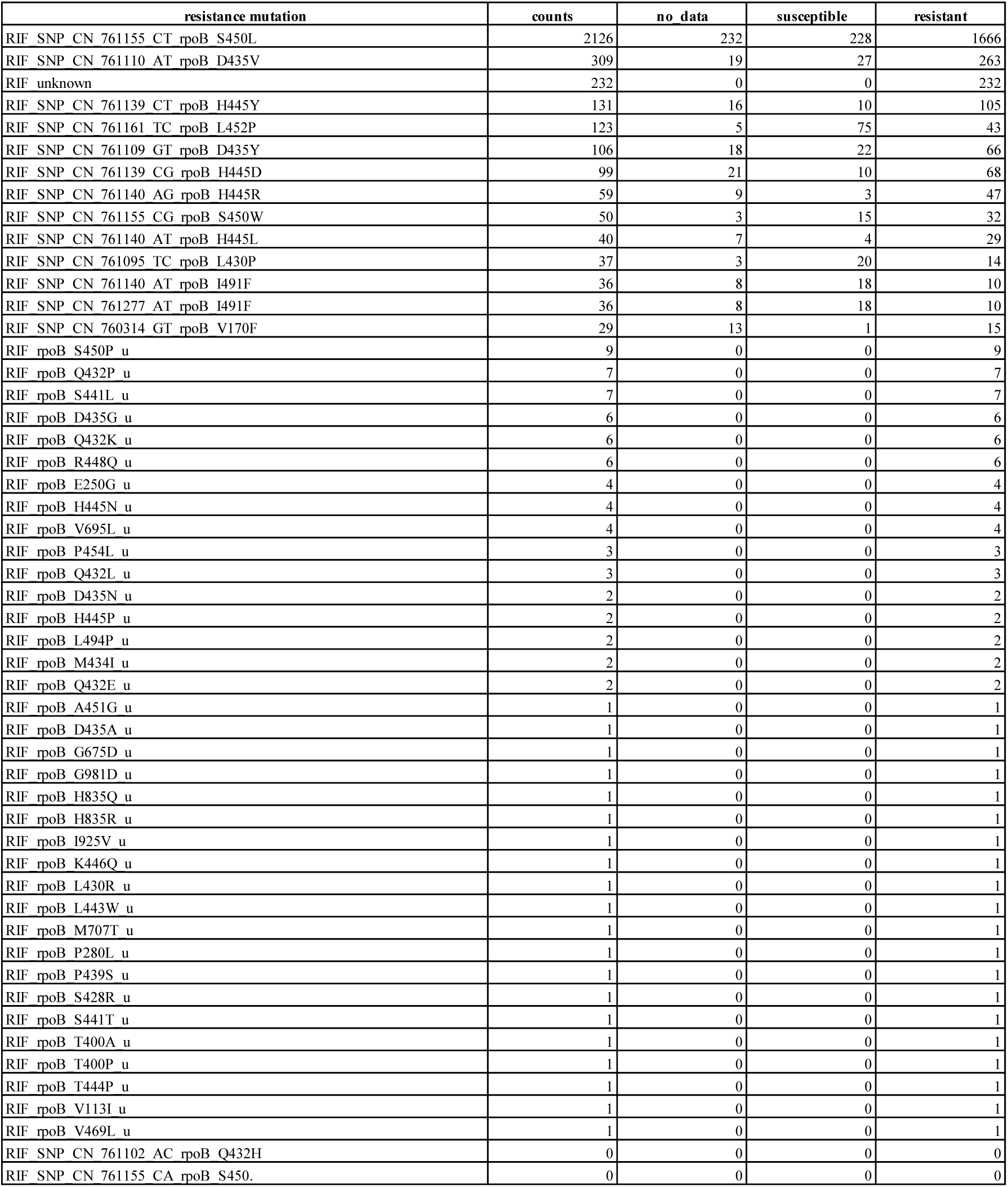
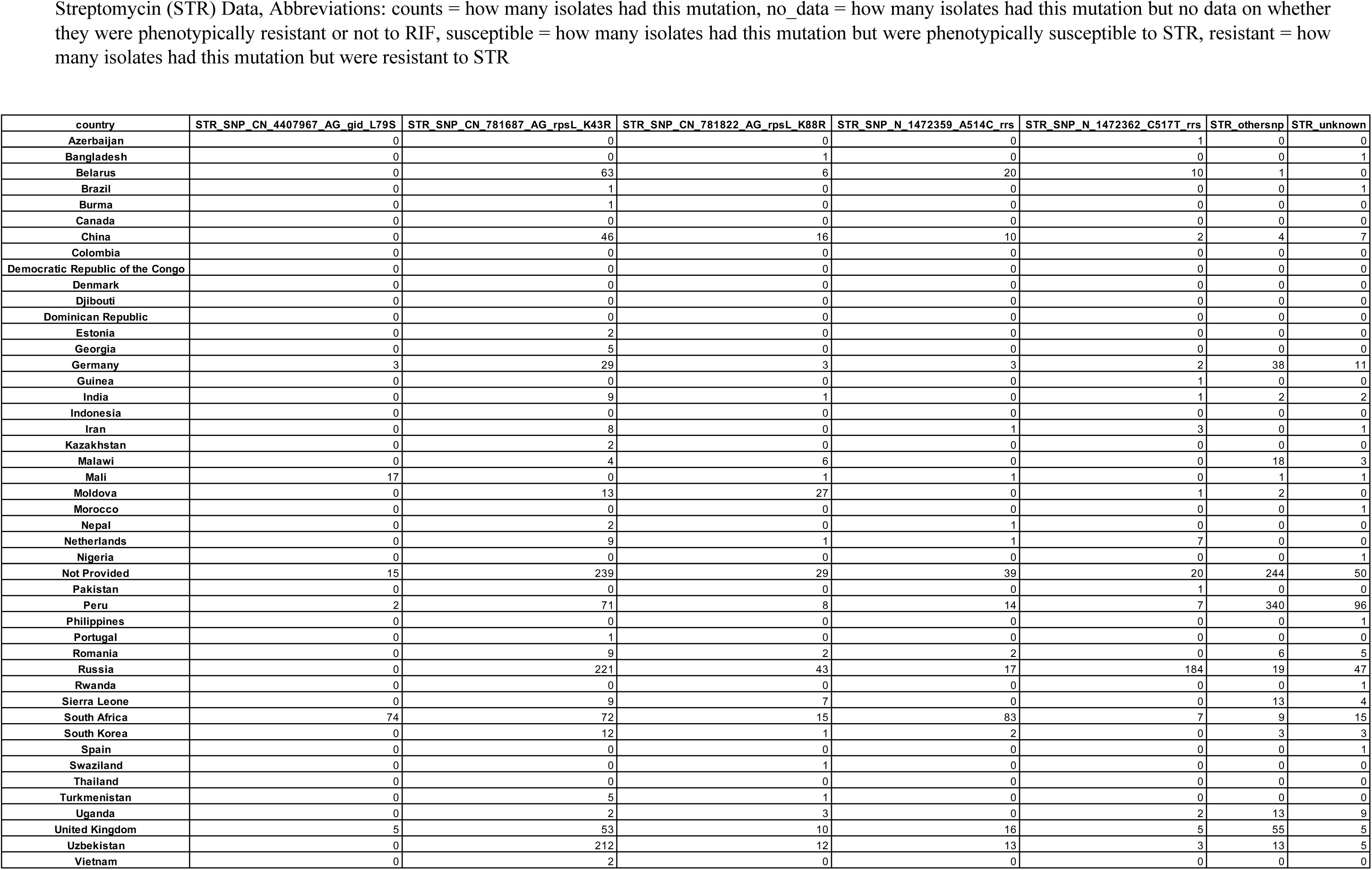
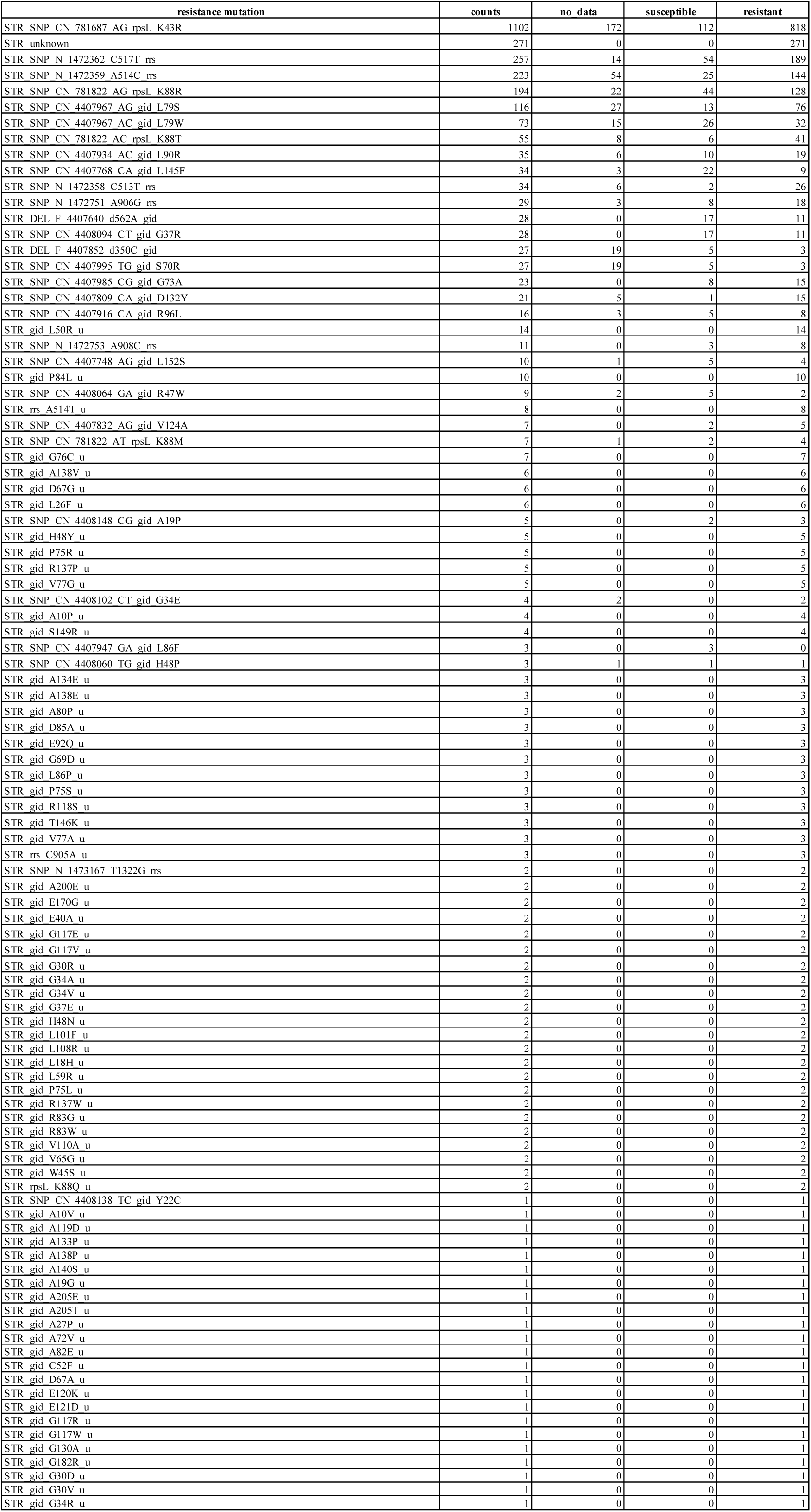
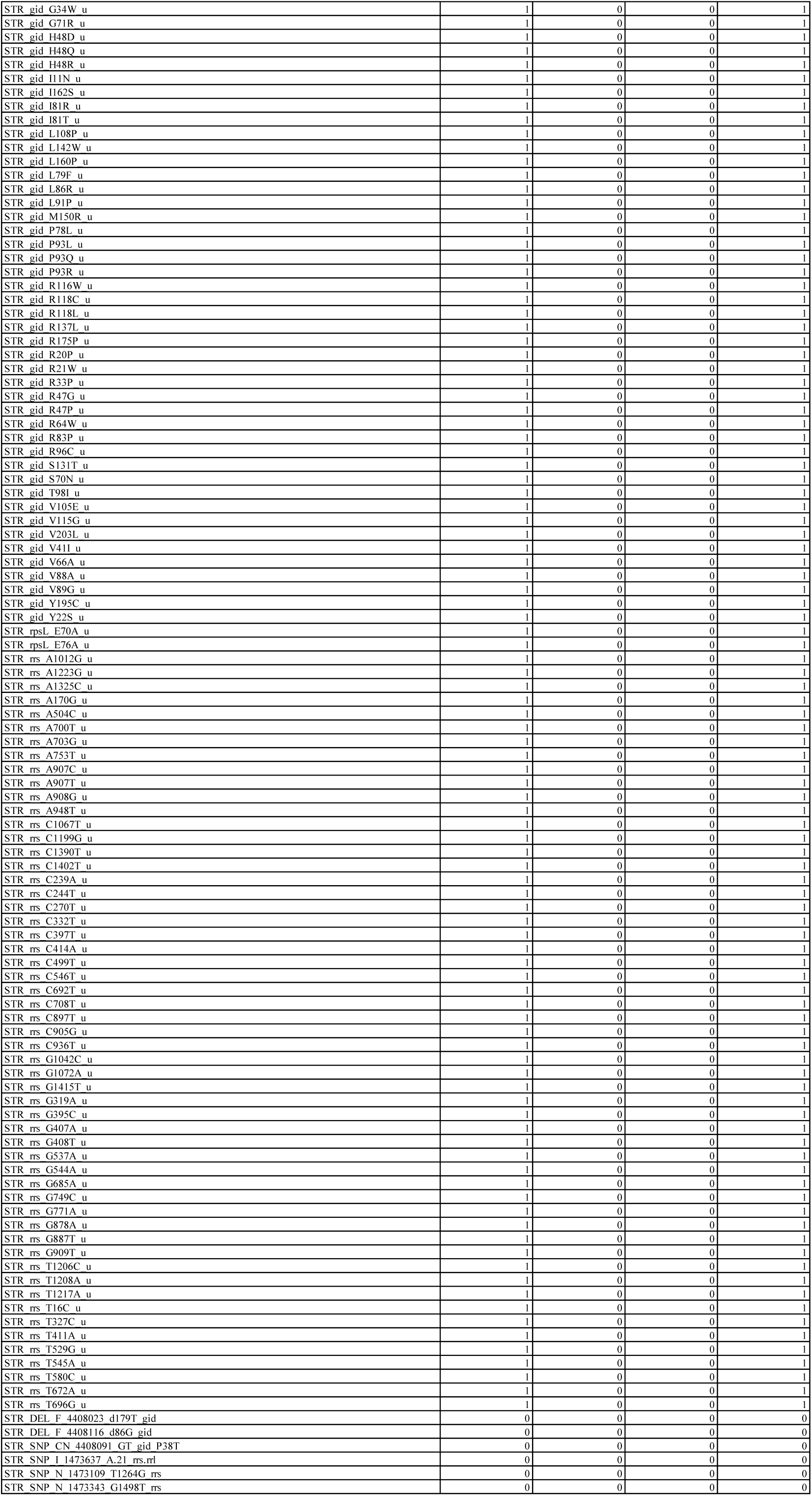
Resistance Mutation Counts Per Country and Per Isolate Phenotype.

**Supplementary Figure 7.**
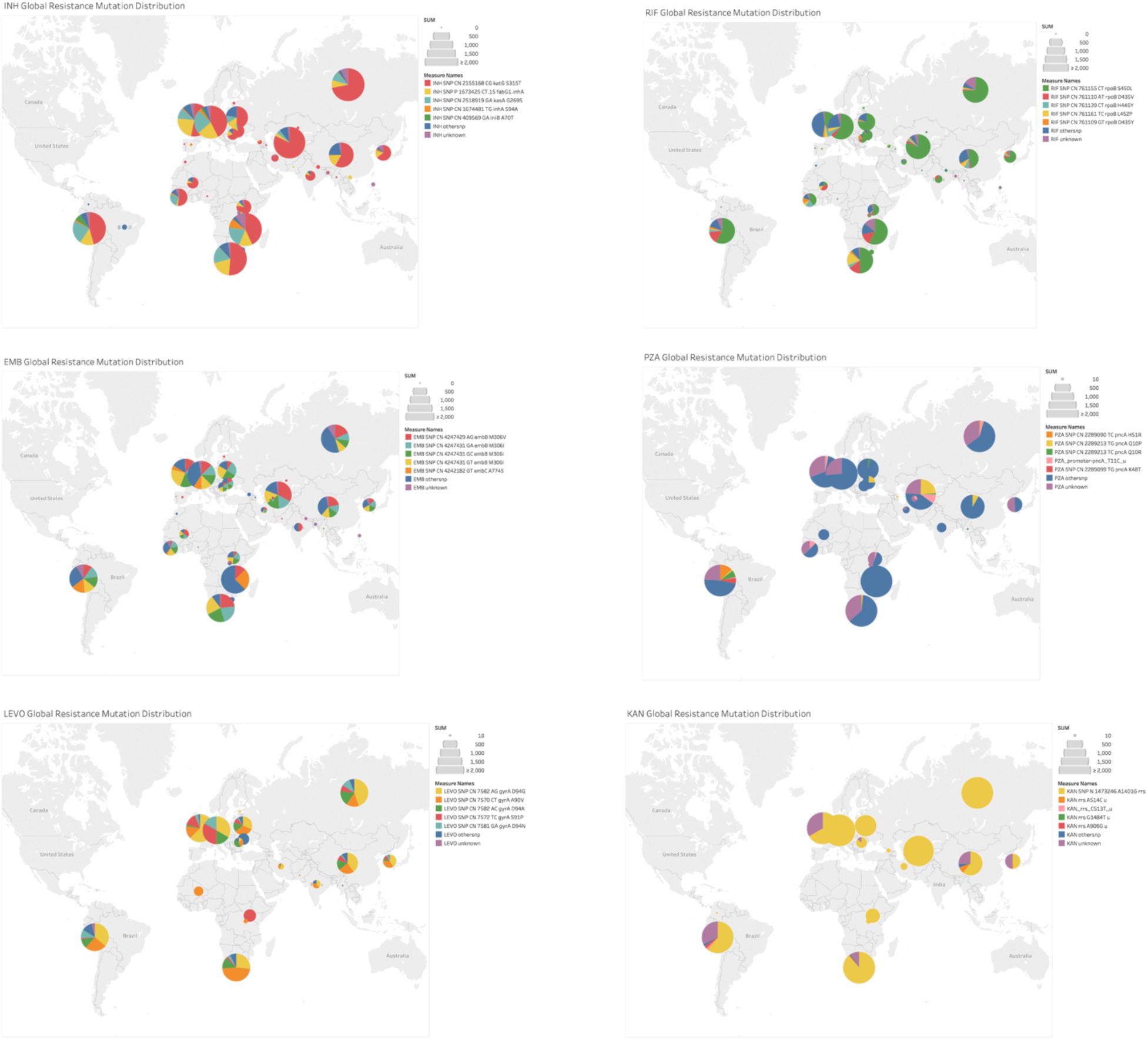
Distribution of resistance mutations for six drugs (n=9385). Pie chart size is proportional isolate number from each country. Mutations listed in order of frequency. Full data provided in Supplementary Table 6.

**Supplementary Table 8.**
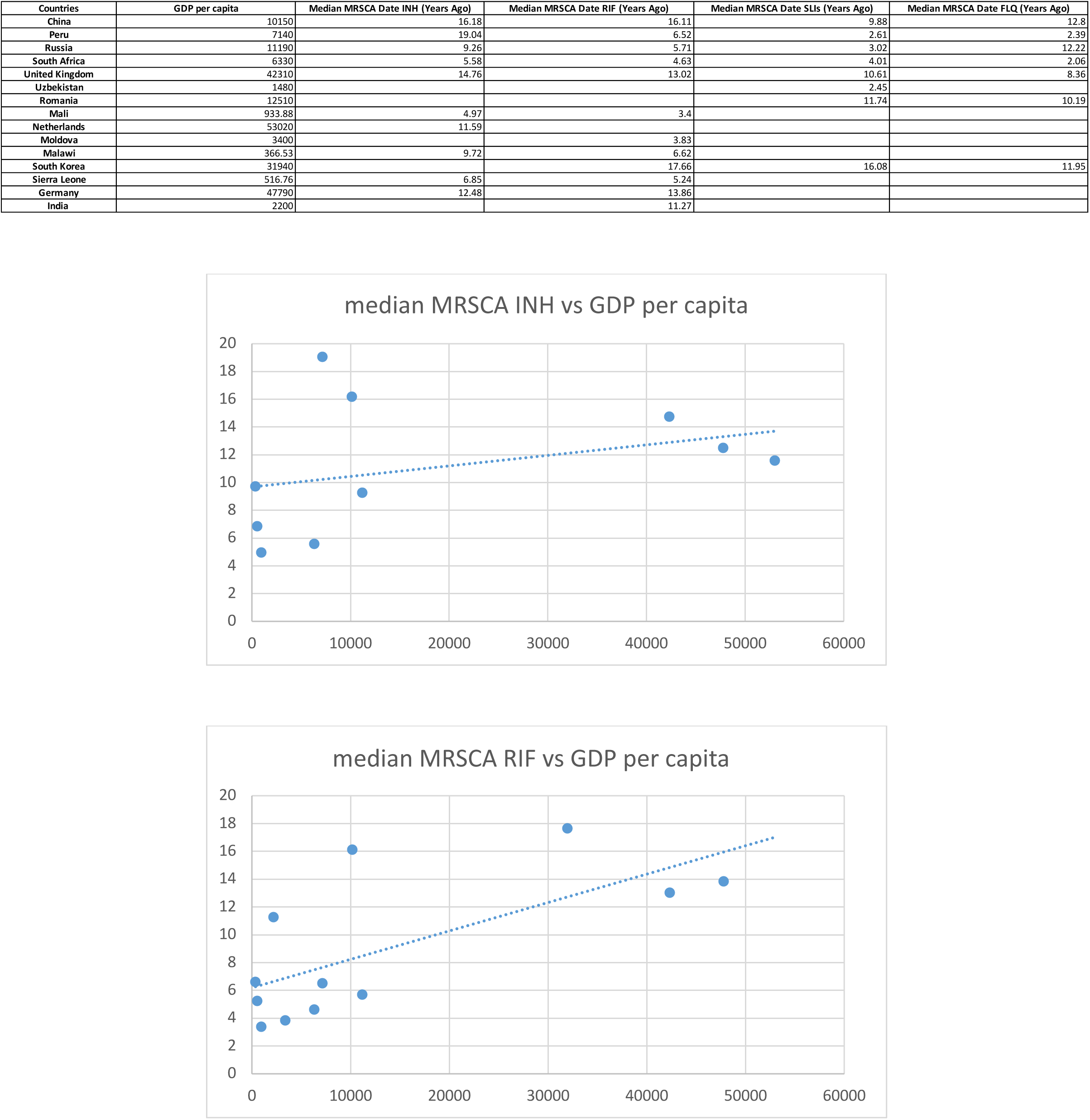
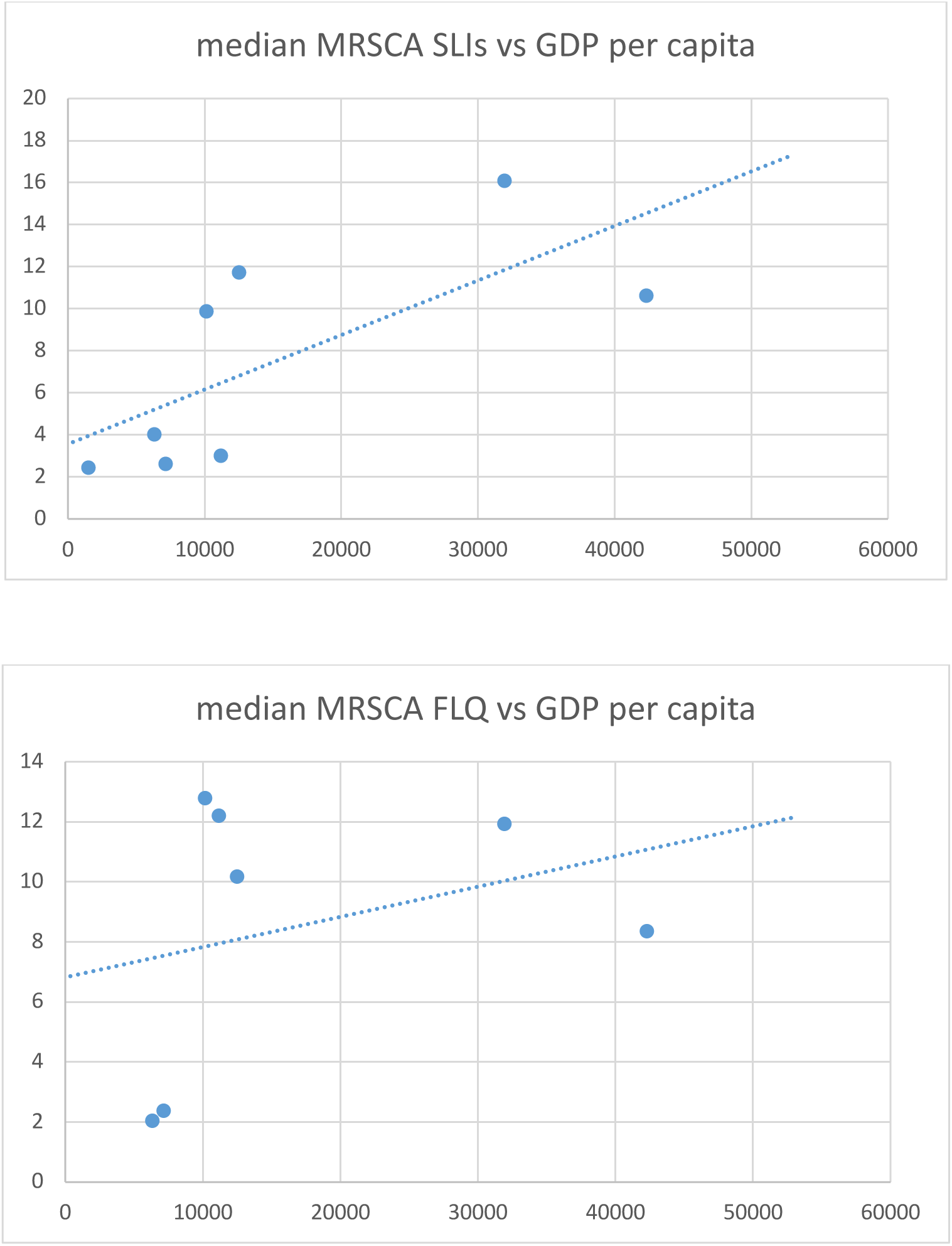
GDP Per Capita Verses Median MRSCA date for INH, RIF, SLIs, and FLQ.

**Supplementary Table 9.** **Commercial Diagnostics**^21,22,23,24,25^:

| Drug | Gene | Location |
| --- | --- | --- |
| Isoniazid | katG codon | 315 |
| Isoniazid | inhA promoter | -15,-16,-8 |
| Rifamycin | rpoB codons | 424-454 |
| Aminoglycosides | rrs | 1401,1402 |
| Aminoglycosides | eis | -10 to -14, -37 |
| Fluoroquinolones | gyrA | 89-94 |
| Fluoroquinolones | gyrB | 500-541 |

\*E-coli 505-534

**Supplementary Table 10.** Genes Searched Per Drug.

| Drug | Genes Searched |
| --- | --- |
| AMK | rrs |
| PAS | thyA, inter-thyA-Rv2765, folC, inter-thyX-hsdS.1 |
| EMB | embA, embB, embC, iniB, inter-embC-embA |
| CAP | rrs, tlyA |
| KAN | rrs, inter-eis-Rv2417c |
| CIP | gyrB, gyrA |
| INH | inhA, iniB, embB, inter-Rv1482c-fabG1, ahpC, inter-oxyR'-ahpC, inter-embC-embA, kasA, katG, fabG1 |
| STR | gid, rpsL, rrs, inter-rrs-rrl |
| RIF | rpoB |
| LEVO | gyrB, gyrA |
| ETH | inhA, inter-Rv1482c-fabG1, ethA, inter-ethA-ethR |
| OFLX | gyrB, gyrA |
| PZA | inter-pncA-Rv2044c, pncA, rpsA |

**Supplementary Table 11.**
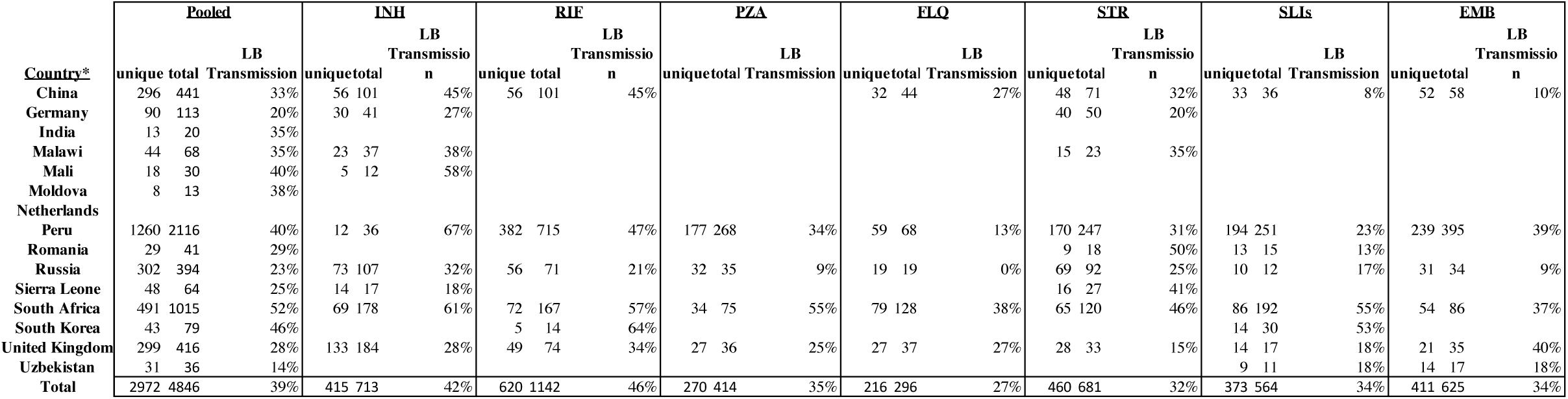
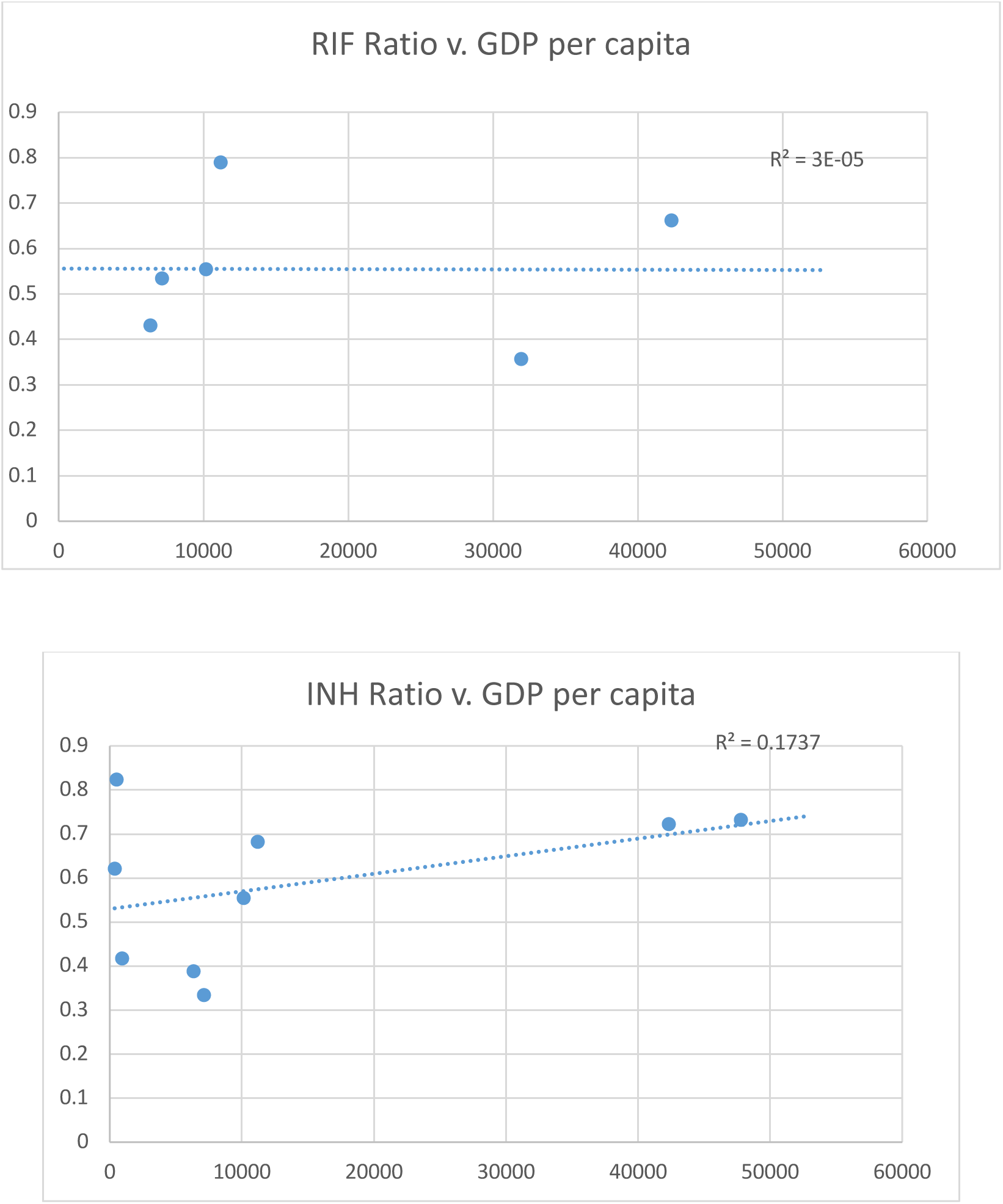
Transmission rate estimates and association with GDP by country. Transmission rate X/Y indicates X unique MRSCA dates/Y number of MRSCA dates for a specific country and drug. Legend: Unique = number of unique MRSCA dates for drug and country, total = total number of dated resistant isolates for drug and country, Lower Bound (LB) = (total-unique)/total, Note: Data for country fewer than 10 resistant isolates per drug not shown.

**Supplementary Table 12.** MRSCA age is not associated with the earliest date of drug introduction into clinical use.

| Drug | Median MRSCA | Date Introduced Years ago |
| --- | --- | --- |
| INH | 11.4 | 67 |
| RIF | 7.61 | 54 |
| PZA | 3.56 | 67 |
| FLQ | 4.81 | 34 |
| STR | 7.67 | 75 |
| SLIs | 4.02 | 61 |
| EMB | 5.01 | 54 |
| ETH | 4.01 | 59 |
| CYS | 2.05 | 64 |

**Supplementary Table 13.**
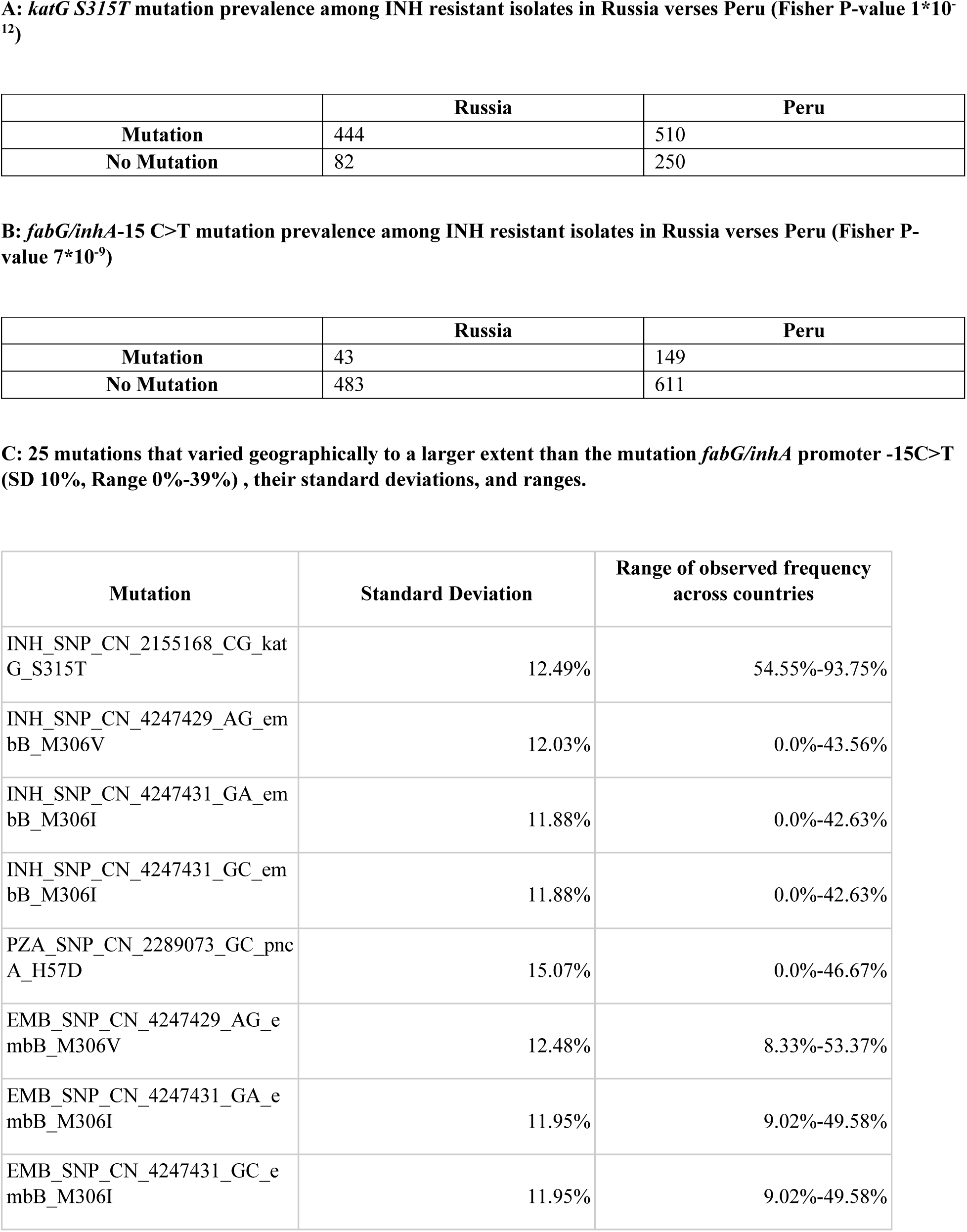
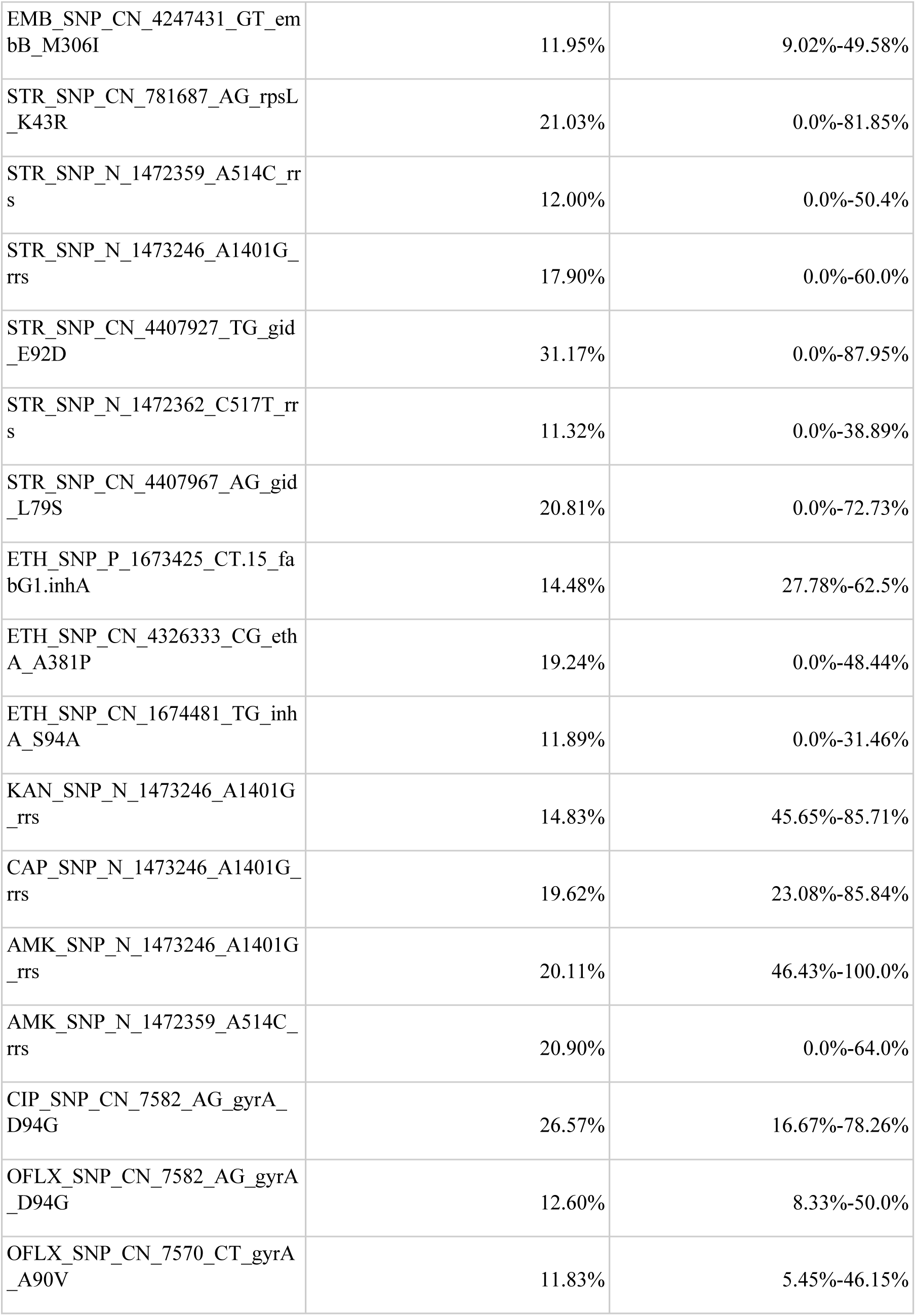
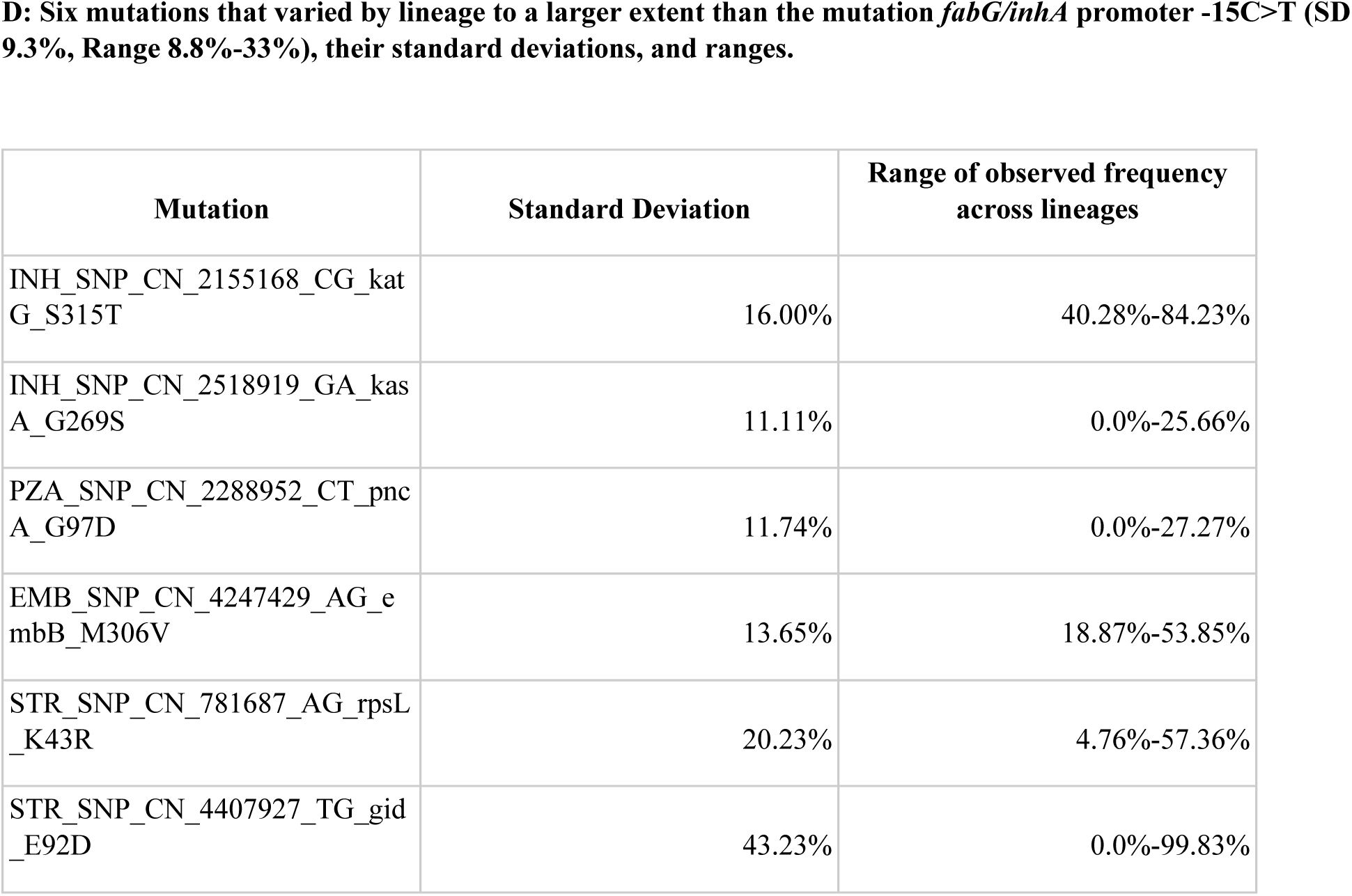
Geographic variance of resistance mutations.

**Supplementary Table 14.** Sensitivity and Specificity of commercial tests for INH, RIF, SLIS, and FLQ in five countries with the largest number of phenotyped strained: Russia, South Africa, Peru, Uzbekistan, and United Kingdom.

| Drug | <u>commercial test-Peru</u> |  | <u>commercial test-Russia</u> |  | <u>commercial test-South Africa</u> |  | <u>commercial test-Uzbekistan</u> |  | <u>commercial test-United Kingdom</u> |  |
| --- | --- | --- | --- | --- | --- | --- | --- | --- | --- | --- |
|  | Sensitivity | Specificity | Sensitivity | Specificity | Sensitivity | Specificity | Sensitivity | Specificity | Sensitivity | Specificity |
| INH | 84% (641/760) | 90% (151/168) | 87% (459/526) | 91% (281/310) | 88% (168/190) | 72% (444/619) | 90% (238/264) | NA 0/0 | 87% (245/283) | 99% (1559/1567) |
| RIF | 90% (623/692) | 89% (209/236) | 82% (347/425) | 88% (363/412) | 95% (158/166) | 70% (433/616) | 95% (248/262) | 50% (1/2) | 91% (86/95) | 99% (1742/1750) |
| FLQ | 38% (46/121) | 98% (788/801) | 38% (51/133) | 94% (275/292) | 77% (86/111) | 87% (510/585) | 0% (0/7) | 100% (213/213) | 90% (26/29) | 98% (295/301) |
| SLI | 47% (101/214) | 98% (692/708) | 77% (63/82) | 48% (163/343) | 77% (101/131) | 91% (486/536) | 79% (44/56) | 66% (109/164) | 90% (9/10) | 100% (83/83) |

